# An alternative spliceosome defined by distinct snRNAs in early zebrafish embryogenesis

**DOI:** 10.1101/858944

**Authors:** Johanna F. B. Pagano, Rob J. Dekker, Wim A. Ensink, Marina van Olst, Alex Bos, Selina van Leeuwen, Wim C. de Leeuw, Ulrike Nehrdich, Herman P. Spaink, Han Rauwerda, Martijs J. Jonker, Timo M. Breit

## Abstract

Splicing removes intronic RNA sequences from pre-mRNA molecules and enables, by alternative splicing, the generation of multiple unique RNA molecules from a single gene. As such, splicing is an essential part of the whole translation system of a cell. The spliceosome is a ribonucleoprotein complex in which five small nuclear RNAs (snRNAs) are involved; U1, U2, U4, U5, and U6. For each of these snRNAs there are variant gene copies present in a genome. Furthermore, in many eukaryotic species there is an alternative, minor spliceosome that can splice a small number of specific introns. As we previously discovered an embryogenesis-specific ribosomal system in zebrafish early embryogenesis based on variant rRNA and snoRNA expression, we hypothesized that there may also be an embryogenesis-specific spliceosome. An inventory of zebrafish snRNA genes revealed clustered and dispersed loci for all but U2 major snRNAs. For each minor spliceosome snRNA, just one gene locus was found. Since complete snRNA molecules are hard to sequence, we employed a combined PCR-sequencing approach to measure the individual snRNA-variant presence. Analysis of egg and male-adult samples revealed embryogenesis-specific and somatic-specific variants for each major snRNA. These variants have substantial sequence differences, yet none in their mRNA binding sites. Given that many of the sequence differences are found in loop structures indicate possible alternative protein binding. Altogether, with this study we established that the spliceosome is also an element of the embryogenesis-specific translation system in zebrafish.

## INTRODUCTION

Alternative splicing is fundamental for gene regulation and the generation of different transcripts and/or proteins from an individual gene in eukaryotes (1). Splicing is executed by the spliceosome and removes intronic sequences from pre-mRNA during the maturation process in which the exonic sequences eventually form the mRNA (2,3).The spliceosome is a molecular complex formed by hundreds of proteins and five essential small-nuclear RNAs (snRNAs) that are typically located in the nucleus. The size of these small RNA molecules ranges from 118 nucleotides (nt) to 191 nt. As they are uracil rich, they are called U1, U2, U4, U5 and U6 snRNAs. Next to this major spliceosome, a minor (or U12 dependent) spliceosome exists in many eukaryotic species, which is involved in the splicing of a relative small number of specific introns (4). The snRNAs involved in the minor spliceosome are: U11, U12, U4atac, and U6atac, completed by the U5 from the major spliceosome (4).

As splicing is at the core of the cellular translation system, the sequences of the involved snRNA are highly conserved across species. At the same time, many non-canonical variants and gene copies of the major snRNA genes are present within the same organisms (5–9). This raises the question why these variants exist and what role they might play in translation. Although expression of these variants has been extensively studied, there is still not a clear understanding for the existence of these snRNA variants (10).

A indication to the function of variant snRNAs may lie in the fact that several studies have observed differentially expression of variants during development. For instance in Drosophila several variants are expressed during early embryogenesis, but eventually one variant gradually dominates expression (11,12). Similar expression patterns for snRNA variants were observed in *Xenopus* (13–15), mouse (12,16), sea urchin (17,18), and flatworm (19). Comparable findings have been reported for: snRNA U2 in silk moth (20) and *Dictyostelium dicoideum* (21); snRNA U4 in chicken (22); and snRNA U5 in *Drosophila* (5) and in human (6). snRNAs also display a tissue-preferred expression, which implicate them in different disease pathologies such as several neurological diseases (7,10,23,24).

The fact that there are snRNAs which variants are differentially expressed during embryogenesis relates to our previous findings where we discovered distinct maternal-types of rRNAs and snoRNAs specifically expressed during early zebrafish embryogenesis (25). These maternal-type RNAs seem to be part of a distinct early embryogenesis-specific translation machinery, which is gradually replaced during embryogenesis by a somatic type. Combining the snRNA and our rRNA plus snoRNA findings, lead us to hypothesize that there might also be distinct maternal-type snRNAs contributing to the embryogenesis-specific translation machinery.

As in our hands, snRNAs cannot consistently be sequenced by standard next-generation sequencing protocols, probably due to strong secondary structures and ample RNA modifications, we employed a wet-lab approach combining RT-PCR, for amplification, and DNA sequencing, for identification, of snRNAs. Using this approach on egg and mature adult-male samples, we were able to show that for each snRNA, similar to rRNA and snoRNA, there are maternal-type snRNAs that are uniquely present in zebrafish eggs. This means that snRNAs are a part of the zebrafish unique embryogenesis-specific translation machinery.

## RESULTS AND DISCUSSION

### Cataloguing the zebrafish snRNAs

In earlier studies we discovered cellular components related to the transcription machinery that have distinct embryogenesis expression or somatic expression during zebrafish development. We reported on 5S (26), 5.8S, 18S, and 28S (27), plus small-derived components from those molecules (28), as well as snoRNAs (25). As it is clear that in zebrafish oogenesis and early embryogenesis a different transcription machinery is employed, it is obvious to investigate other components of this system. Hence, we here focused on the small nuclear RNAs (snRNAs) of the spliceosome to determine whether their variants also display distinct expression profiles. We started by making an inventory of all snRNA elements present in the zebrafish genome based on the annotated snRNA sequences from the Ensembl database (29). In total, 541 snRNA loci were retrieved for the major spliceosome and seven snRNA loci for the minor spliceosome from the database. Given the many snRNA loci for the major spliceosome, we compared them by sequence alignment per snRNA (Supplemental Figure SFig1). Several snRNA sequences appeared aberrant and it turned out that these sequences partly existed of retrotransposon sequences (30), therefore they were excluded (Supplemental Table ST1). To complete our reference set of snRNAs, the database-derived snRNA sequences were used to explore the zebrafish genome for yet unannotated major and minor spliceosome snRNA loci. This resulted in a total of 958 snRNA loci (Table 1, Supplemental Table ST1 and Supplemental File SF1).

**Table 1:**
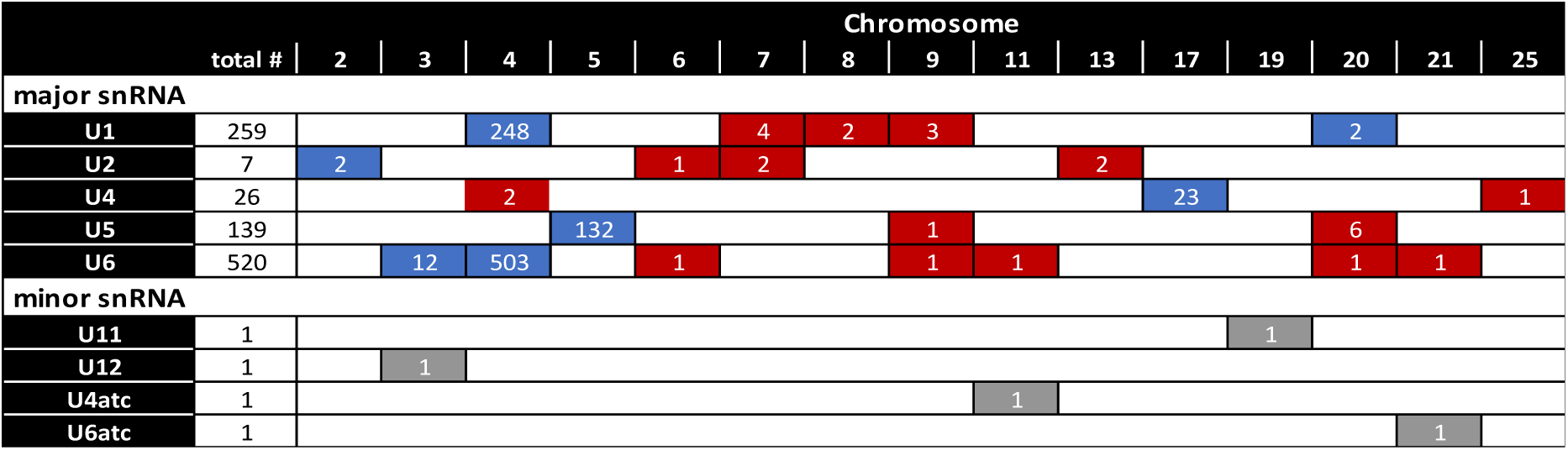
Genomic distribution of maternal and somatic zebrafish snRNA genes.

|  |  |  |  |  |  |  |  |  |  |  |  |  |  |  |  |  |
| --- | --- | --- | --- | --- | --- | --- | --- | --- | --- | --- | --- | --- | --- | --- | --- | --- |
|  | total # | 2 | 3 | 4 | 5 | 6 | 7 | 8 | 9 | 11 | 13 | 17 | 19 | 20 | 21 | 25 |
| major snRNA |  |  |  |  |  |  |  |  |  |  |  |  |  |  |  |  |
| U1 | 259 |  |  | 248 |  |  | 4 | 2 | 3 |  |  |  |  | 2 |  |  |
| U2 | 7 | 2 |  |  |  | 1 | 2 |  |  |  | 2 |  |  |  |  |  |
| U4 | 26 |  |  | 2 |  |  |  |  |  |  |  | 23 |  |  |  | 1 |
| U5 | 139 |  |  |  | 132 |  |  |  | 1 |  |  |  |  | 6 |  |  |
| U6 | 520 |  | 12 | 503 |  | 1 |  |  | 1 | 1 |  |  |  | 1 | 1 |  |
| minor snRNA |  |  |  |  |  |  |  |  |  |  |  |  |  |  |  |  |
| U11 | 1 |  |  |  |  |  |  |  |  |  |  |  | 1 |  |  |  |
| U12 | 1 |  | 1 |  |  |  |  |  |  |  |  |  |  |  |  |  |
| U4atc | 1 |  |  |  |  |  |  |  |  | 1 |  |  |  |  |  |  |
| U6atc | 1 |  |  |  |  |  |  |  |  |  |  |  |  |  | 1 |  |

The alignment of the sequences within each major spliceosome snRNA showed two clusters for each snRNA (Supplemental File SF1). The distribution of these two snRNA clusters coincides with a clear genomic organizational preference: either in condense genomic repeats (Table 1, blue), or scattered throughout the genome (Table 1, red). Obviously, the discovery of two sequence clusters for each major spliceosome snRNA is in line with the possible existence of maternal-type and somatic-type snRNA spliceosomes.

In contrast; for the minor spliceosome, U12 snRNA is present only once in the genome, which means that for U12 snRNA no maternal-type variant can exist. For all three other minor spliceosome-specific snRNAs, just two loci appeared to be present in the genome (Supplemental Table ST1). Because the observed sequences within each of these minor spliceosome snRNAs are so different, we questioned whether they are legitimate snRNA loci. By analyzing the expression of all these snRNA loci, it could be determined that for each minor spliceosome snRNAs, only one locus is expressed (Table 1, grey, Supplemental Table ST2). This precludes the existence of an alternative, completely embryogenesis-specific minor spliceosome and we therefore excluded these minor spliceosome snRNAs from the subsequent analyses.

### Embryogenesis-specific snRNA variants

After cataloguing all snRNAs in the zebrafish genome, we investigated whether embryogenesis-specific snRNAs exist by examining their expression in eggs and somatic tissue. However, it became quickly evident that (complete) snRNA molecules, similarly to tRNA, 5S, and snoRNA molecules, cannot readily be sequenced by standard smallRNA-seq procedures, probably due to their robust secondary configurations, as well as possible 3’ modifications. Only small numbers of partial snRNA sequences were observed, which did show a hint of differential snRNA expression between egg and adult tissue. To tackle this technical problem, we performed a RT-PCR-qSeq analysis effectively circumventing the snRNA 3’ issues. For this, we selected the sequence most prominently present in the genome for each cluster of every snRNAs of the major spliceosome (Figure 1A). The differences between these pairs of snRNA sequences ranges from 11 nucleotides (U1 and U6) to 48 nucleotides (U5) (Figure 1A, Supplemental Figure SFig1). We designed (degenerated) RT-PCR-primers (Figure 1A) that amplified virtual all snRNAs, after which the PCR products were sequenced and identified by mapping to the snRNA sequences. Using this procedure at least 0.5 M reads were obtained for each snRNA (Supplemental Table ST3). These results revealed that for each snRNA one variant accounts for almost all snRNA in the egg samples, whereas the other variant makes up the snRNAs in adult zebrafish (Figure 1B). Similar to rRNA and snoRNA, the snRNA variant present in egg is called maternal-type versus the somatic-type variant in adult tissue (Figure 1A).

**Figure 1.**
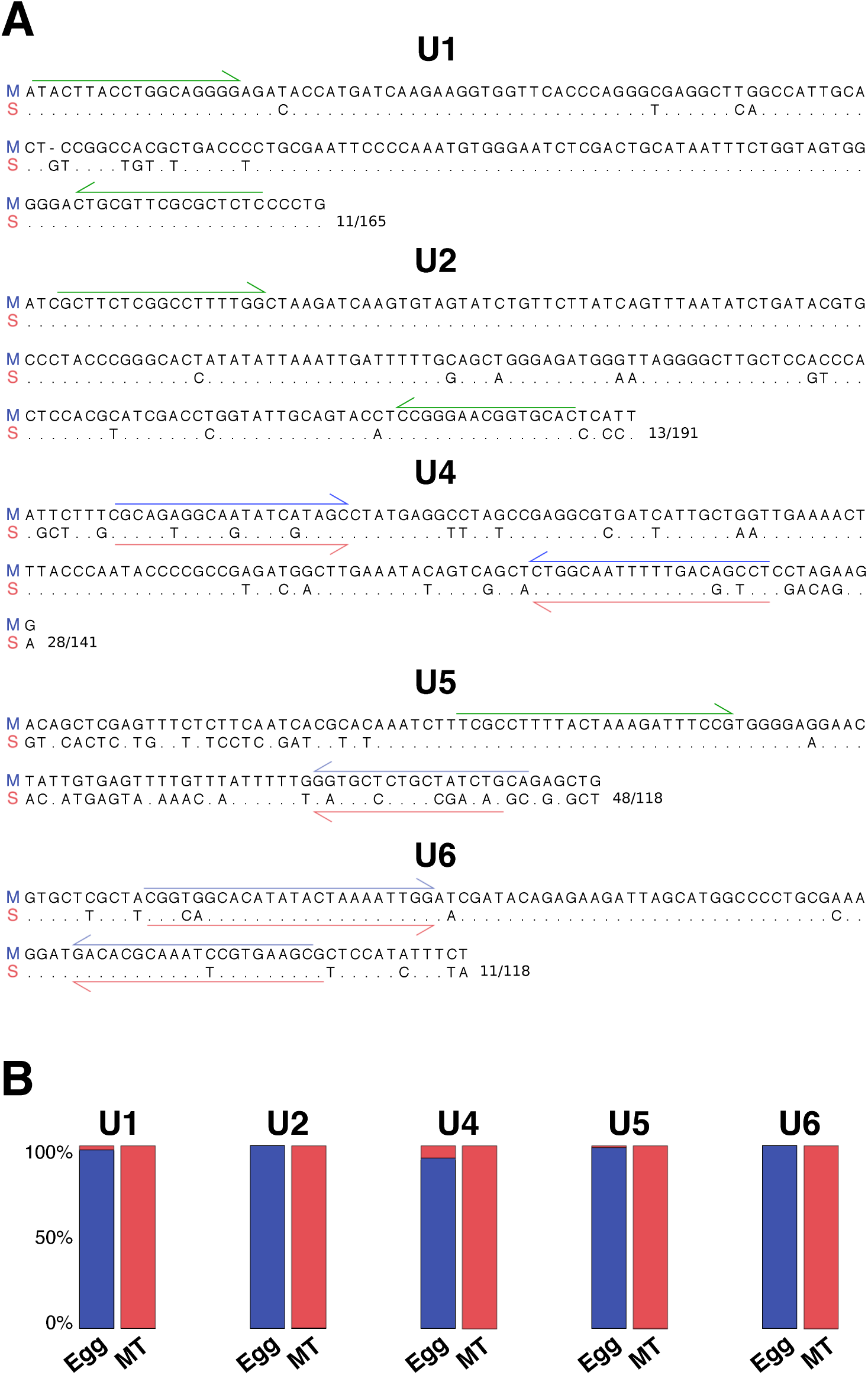
Maternal-type and somatic-type snRNA sequences and expression. **A**: Sequence comparison of selected maternal-type (M) and somatic-type (S) snRNA variants; identical nucleotides are indicated as dots, while gaps as dashes (For sequence alignment of all snRNAs cf. Supplemental File SF2). The RT-PCR primers are indicated with half arrows: maternal-type specific (blue); somatic-type specific (red); and non-distinctive (green). **B**: Relative expression of maternal-type (blue) and somatic-type (red) variants for each snRNA indicated by comparative percentage calculated using the RT-PCR-qSeq approach on RNA from egg and male tail tissue (MT).

To check the RT-PCR-qSeq results we developed a qRT-PCR analyses for snRNA U1 and snRNA U5. This qRT-PCR analysis confirmed the exclusive presence of embryogenesis-specific snRNA variants in egg samples (Supplemental Figure SFig2).

### Differences between maternal- and somatic-types snRNAs

In order to assign any functional relevance to the embryogenesis-specific snRNAs, nucleotide difference as compared to the somatic-type snRNAs were investigated (Figure 1A). One of the distinctions between the major and minor spliceosome snRNAs lies in their nucleotide sequences that bind to the mRNA, thus allowing each system to splice distinct introns. However, even though there are many sequence differences between the maternal-type and the somatic-type snRNAs, none of them involve the mRNA binding sites in these snRNAs (Figure 2 and Supplemental Figure SFig1). It turned out that the snRNA sequence differences are often located in specific parts of the secondary structure (Figure 2). For instance, for U1, all but one differences are located in one stem-loop and in U2 they are confined to one side of the structure (Figure 2). In general, many of the differences are found in the loops, which are thought to be specific binding locations for spliceosomal proteins. Hence, this would indicate that the embryogenesis spliceosome, besides specific snRNAs also may comprise (embryogenesis-)specific proteins.

**Figure 2.**
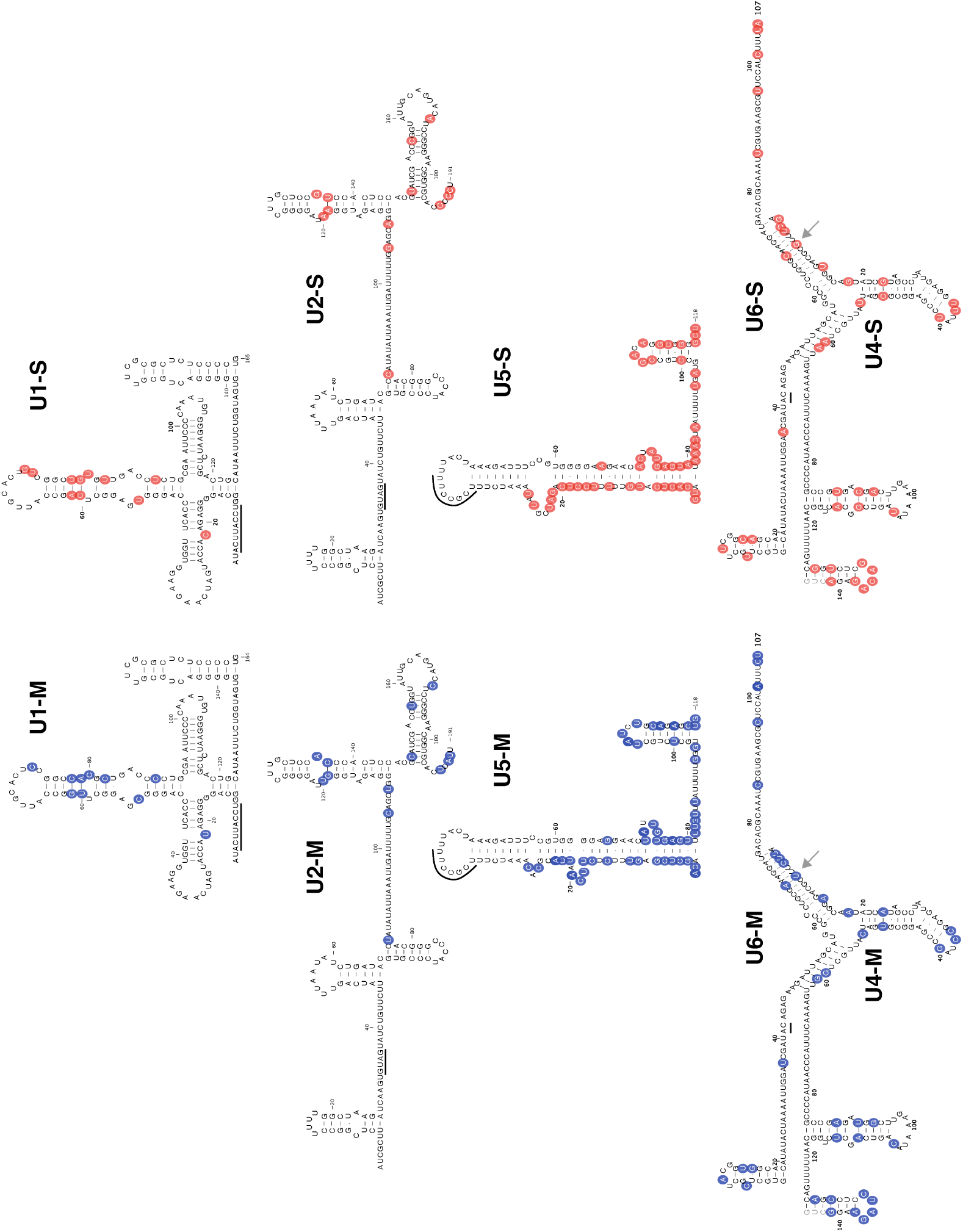
Sequence differences between maternal-type and somatic-type snRNAs. The secondary structure is presented for each maternal-type and somatic-type zebrafish snRNA. These structures are an adaptation from Figure 4 in reference (4) and also indicate the specific binding sites between U4 and U6. The arrows indicate the base pair that coevolved within one of the U4 – U6 interaction sites. The circles indicate nucleotides that are different between the maternal-type (blue) and somatic-type (red) sequences. The sites interacting with the pre-mRNA are underlined.

Despite many apparently co-evolved nucleotide pairs in stems of the snRNAs (Figure 2), there seems to be only one co-evolved nucleotide pair in the interaction site between U4 and U6. However, as this nucleotide is right in the middle of the largest interaction site, it might actually prevent the binding of somatic-type U4 to maternal-type U6 and vice versa.

## CONCLUDING REMARKS

In this study we reveal the existence of an embryogenesis-specific major spliceosome in zebrafish, consisting of at least a distinct set of U1, U2, U4, U5 and U6 snRNAs. We do not know if this embryogenesis-specific spliceosome also contains embryogenesis-specific proteins, yet the position of the distinguishing nucleotides in the loops of the maternal-type snRNA structure would suggest this. However, a quick scan of some of the major spliceosomal protein genes revealed that they are present with just one copy in the zebrafish genome, which effectively rules out any embryogenesis-specific spliceosomal protein gene variants. Yet, the genes for these proteins do contain introns, which have in general an important function with respect to production of alternative transcripts plus alternative proteins. This leads to the intriguing question, whether the maternal-type snRNAs will produce embryogenesis-specific alternative-spliced transcripts and thus embryogenesis-specific spliceosome proteins for the embryogenesis-specific variant of the major spliceosome.

With respect to the minor spliceosome, which acts on different splicing sites; we did not find any maternal-type snRNA variants for U11, U12, U4atc, and U6atc. This however does not lead to the conclusion that no embryogenesis-specific minor spliceosome exists. On the contrary, the minor spliceosome is always completed by the U5 snRNA from the major spliceosome (4) and because in eggs virtually only the maternal-type U5 snRNA is present, this U5 variant is used in combination with the other four ubiquitously-expressed minor spliceosome snRNAs. Even though only one snRNA in this splicing system is different, we would argue that this makes the whole minor spliceosome in eggs embryogenesis-specific. This might also explain the observation that the maternal-type U5 snRNA sequence differs a noteworthy 41% compared to the somatic-type, whereas in all other snRNAs this difference is just 7% to 20%. Yet, the mRNA binding loop that holds the exons together (32) and the associated stem are identical between maternal-type, somatic-type and human U5 snRNA. Besides this core the zebrafish U5 snRNA variants are almost completely different (53%). The functional implications remain elusive for now.

Although the differences between the variants of each snRNA are quite substantial, their secondary structure appears to be relatively unaltered. This is also due to many co-evolved nucleotides in the stem sequences. Similar to what we observed previously for the 45S rRNAs, the sequence homology between the somatic-type and maternal-type snRNA variants is often lower than snRNAs in distantly-related species. For instance, apart from the U1 snRNA, the sequence homology between maternal-type and somatic-type zebrafish snRNA variants is lower than between the somatic-type zebrafish variant and the human snRNA sequences (Supplemental Figure SFig1). The considerable sequence difference of the snRNA variants within the zebrafish hints on an intriguing need for two very different spliceosomal systems in early embryogenesis and adult zebrafish.

The genomic distribution of the maternal-type and somatic-type snRNA variants is also similar to that of the small snoRNAs and 5S rRNAs in that U1, U5 and U6 have a relative high number of maternal-type snRNA genes, whereas there are only a few somatic-type genes. This correlates with the fact that the U1, U5 and U6 are the snRNA that directly interact with the mRNA. However, it is still somewhat surprising that the maternal-type U2 variant is present only in two loci. Also similar to 5S rRNA, the majority of multiple U1, U5 and U6 snRNA genes are organized in a strictly repeated manner, but with several interruptions caused by retrotransposons (26). It was also clear that some snRNA genes were only partially present in the genome (Supplemental Table ST1). Together this would be in line with the reported presence of retrotransposons in snRNA gene repeats (33,34). A quick scan of these snRNA interrupting sequences, revealed the presence of retrotransposons with and without intact open reading frames. Alike 5S loci, we expect that retrotransposons might play a role in maintaining the needed number of gene copies in the snRNA repeats.

The strict differential expression of maternal-type and somatic-type snRNAs obviously is regulated via the promotor and auxiliary sequences related to the snRNA genes and we expect that an extensive analysis of all maternal-type and somatic-type snRNA promoter regions will eventually discover the relevant differences. This holds also for mechanisms that are involved in snRNA processing and degradation. For instance, a superficial comparison suggests that the core sequences of the 3’ box, which is involved in the post transcriptional processing of snRNAs (35), are different between the maternal-type and somatic-type snRNA variants (results not shown).

In many other species different snRNA variants have been found, often with differential expression during embryogenesis, such as in *Xenopus* (13), Human (36), mouse (37), sea urchin (17), *Drosophila* (12). The observed zebrafish snRNA variant system is unique in that the maternal-type snRNA is exclusively expressed in oocytes, eggs, and early embryogenesis. All other reports just mention up to 40% higher expression of some variants. This implies that by means of maternal-type snRNAs, together with the maternal-type rRNAs and maternal-type snoRNAs we previously reported, zebrafish employ a distinct translation system specifically for early embryogenesis

## MATERIAL AND METHODS

### Biological materials

Adult zebrafish (strain ABTL) were handled in compliance with local animal welfare regulations and maintained according to standard protocols (http://zfin.org). The breeding of adult fish was approved by the local animal welfare committee (DEC) of the University of Leiden, the Netherlands. All protocols adhered to the international guidelines specified by the EU Animal Protection Directive 86/609/EEC.

For this study samples were used of two pools of unfertilized eggs (oocyte clutches) and two whole male adult zebrafish. The harvesting of the biological materials and RNA-isolation have been described previously (26,27).

### Source data

In this study we use the zebrafish genome version GRCz11 and next-generation sequencing data previously generated by our group (26) and available through the BioProject database with accession number PRJNA347637.

### qRT-PCR analysis

For snRNA U1 and U5, forward and reverse PCR primers were used from the RT-PCR-qSeq analysis, and quantitative real-time PCR (qRT-PCR) probes were designed (Supplemental Table ST4). Reverse transcription was done in two independent reactions for zebrafish clutch (= egg pool) and adult male tail total RNA. SuperScript IV Reverse Transcriptase (Thermo Fisher Scientific) was used according to the manufacturer’s instructions. Separate qRT-PCR analyses were performed on 10-fold dilutions of the cDNAs with the snRNA U1 and U5 primer/probe combinations using a QuantStudio 3 Real-Time PCR System (Thermo Fisher Scientific).

### RT-PCR-qSeq analysis

Forward and reverse PCR primers were designed for maternal-type and somatic-type snRNA variants, in such a way that: 1) as much as possible of the 5’-end of the full-length variants is included in the final amplicon, and 2) generic primers will bind to the maternal-type, as well as the somatic-type variants (Supplemental Table ST4 and Figure 1A). To avoid positive results due to genomic DNA background, small RNA-enriched total RNA was treated twice with 5 μl of RNase-free DNase (Qiagen) for 45 minutes at 37°C. Next, cDNA was prepared from 50 ng of RNA as described in (26). For each sample, reverse transcription was primed using a mixture of all reverse PCR primers. Controls without reverse transcriptase were used to exclude genomic DNA contamination of the RNA in the downstream PCR. Subsequently, standard PCR reactions were performed on 1 ul of cDNA using Q5 High-Fidelity DNA Polymerase (New England Biolabs) and each of the variant PCR primer pairs independently. The resulting amplicons were purified using the QIAquick PCR Purification Kit (Qiagen) according to the manufacturer’s instructions, with the exception that a total of seven volumes of solution PB was added to allow for more efficient binding of fragments <100 bp. Next, the PCR products were phosphorylated using T4 PNK (New England Biolabs) and again purified as described above. Afterwards, the size of the PCR product was verified on a 2200 TapeStation System (Agilent). From 44 ng purified phosphorylated PCR product, barcoded sequencing libraries were prepared using a modified version of the Ion Xpress Plus Fragment Library Kit (Thermo Fisher Scientific) as described previously (26). Massive-parallel sequencing was performed on an Ion Proton System (Thermo Fisher Scientific) using an Ion PI Chip Kit v3.

### Bioinformatics analyses

#### Known snRNA sequences

The initial set of snRNA sequences of *D. rerio* (GRCz11) were downloaded from Ensemble 95 (29) in October 2018 using Biomart by selecting snRNA as *Gene type* (See the RF annotated tabs in Supplemental Table ST1).

#### Discarding “contaminated” snRNA sequences

For each of the five snRNAs a multiple alignment of the downloaded sequences was made. This was done using CLC Genomic Workbench with default settings (gap open cost 20, gap extension 20 end gap cost free, very accurate). Via visual inspection of the alignments (Supplemental Figure SFig1) the snRNA sequences that contained obvious contaminating, non-snRNA sequences, such as retrotransposon sequences, were discarded or truncated.

#### Discovery of new snRNA sequences and loci

The known snRNA sequences were then aligned with the zebrafish genome using BLASTN and all filters and masking unselected. The hits that were at least 95% of the query length were selected. Names were then assigned to the unique sequences.

#### Mapping of the smallRNA-seq reads

The reads from the source data were mapped to the major and minor snRNA sequences using default settings of Bowtie-2 (31).

#### Analysis of the RT-PCR-qSeq

To count the number of maternal-type and somatic-type reads in egg and adult male zebrafish, an exact string search of the maternal-type and somatic-type snRNA sequences was performed for each snRNA. Data normalization was done using the total count of mapped smallRNA-seq reads per sample.

## Supporting information

Supplemental Figure SFig1

Supplemental Figure SFig2

## ACKNOWLEDGMENTS

We acknowledge the support of The Netherlands Organization for Scientific Research (NWO) grant number 834.12.003.

## SUPPLEMENTAL FILES

SF1: Zebrafish snRNA sequence alignments

SF2: Alignment of snRNA sequences to selected maternal-type and somatic-type variant sequences

## SUPPLEMENTAL TABLES

ST1: Genome distribution of zebrafish snRNAs

ST2: Minor spliceosome snRNA read counts

ST3: Read counts from the RT-PCR-qSeq experiment

ST4: Primer sequences

## SUPPLEMENTAL FIGURES

SFig1: Sequence alignments of human snRNAs with maternal-type and somatic-type zebrafish snRNA

SFig2: qRT-PCR results for snRNA U1 and snRNA U5

