## Supplemental Figure SFig1 for "An alternative spliceosome defined by distinct snRNAs in early zebrafish embryogenesis"

## U1

```
H: ATACTTACCTGGCAGGGGAGATACCATGATCACGAAGGTGGTTTTCCCAGGGCGAGGCTTATCCATTGCA
S: .....................C..........A..........CA.......T......C.G........
M: ................................A..........CA...............GG........

H: CT-CCGGATGTGCTGACCCCTGCGATTTCCCCAAATGTGGGAAACTCGACTGCATAATTTGTGGTAGTGG
S: ..GT...C....T.....T......A.................T................C.........
M: ..-....CCAC..............A.................T................C.........

H: GGGACTGCGTTCGCGCTTTCCCCTG
S: .................C.......  16/165
M: .................C.......  12/164
```

## U2

```
H: ATCGCTTCTCGGCCTTTTGGCTAAGATCAAGTGTAGTATCTGTTCTTATCAGTTTAATATCTGATACGTC
S: .....................................................................G
M: .....................................................................G

H: CTCTATCCGAGGACAATATATTAAATGGATTTTTGGAGCAGGGAGATGGAATAGGAGCTTGCTCCGTCCA
S: .C...C...G.C..C...........T............................G..............
M: .C...C...G.C..T...........T........C...T.........GT....G.........AC...

H: CTCCACGCATCGACCTGGTATTGCAGTACCTCCAGGAACGGTGCACCCCCT
S: ...............C.............T...G.............A...  12/191
M: .................................G.............A.T.  17/191
```

## U4

```
H: AGCTTTGCGCAGTGGCAGTATCGTAGCCAATGAGGTTTATCCGAGGCGCGATTATTGCTAATTGAAAACT
S: ............................T.........................................
M: .TTC..T.....A....A....A.....T......CC..G........T...C......GG.........

H: TTTCCCAATACCCCGCCATGACGACTTGAAATATAGTCGGCATTGGCAATTTTTGACAGTCTCTACGGAG
S: ..A..............G........................C..............G.....G..A...
M: ..A..............GA..T.G.........C....A..TC................C...CTA.A..

H: A
S: .   7/141
M: G  30/141
```

## U5

```
H: GCATACTCTGGTTTCTCTTCAGATCGTATAAATCTTTCGCCTTTTACTAAAGATTTCCGTGGAGAGGAAC
S: .T.C....AT....T..C............................................G..A....
M: A..GCTCGA.T..CTCT.CA.TCA..C.C.................................G.......

H: AACTCTGAGTCTTAAGCTAATTTTTTGAGGCCTTGTTCCGACAAGGCTATAT
S: -...A.....A.A-.A............T...C.C-.-.....G.CGGGGC.  22/118
M: T.T.G.....T..--.-.TTA......G.TG..C.G-.TAT.TGCAGAGCTG  45/118
```

## U6

```
H: GTGCTCGCTTCGGCAGCACATATACTAAAATTGGAACGATACAGAGAAGATTAGCATGGCCCCTGCGCAA
S: .....T................................................................
M: .........A...TG....................T...............................A..

H: GGATGACACGCAAATTCGTGAAGCGTTCCATATTTTT
S: ...............................C....A  3/107
M: ...............C.........C.........C.  8/107
```

**Supplemental Figure 1. Sequence alignments of Human snRNAs with maternal and somatic zebrafish snRNA.** Human snRNAs are labeled with an H and maternal and somatic zebrafish snRNAs with an M and an S, respectively. The sequences interacting with the pre-mRNA are boxed with a black line. The U2/U6 interactions are highlighted with yellow, green and pink.
