## Supplemental Figure SFig2 for "An alternative spliceosome defined by distinct snRNAs in early zebrafish embryogenesis"

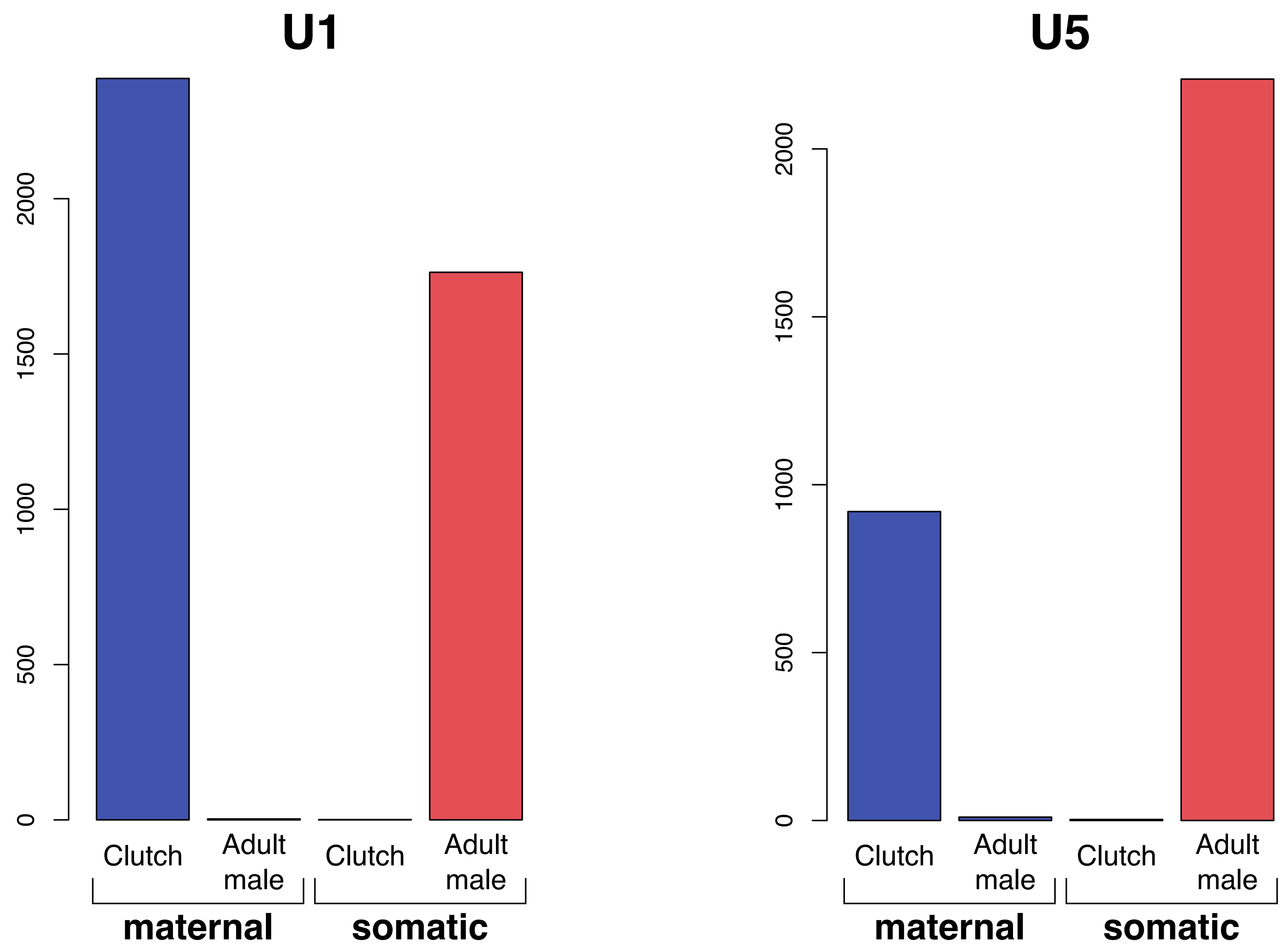


**Supplemental Figure 2. qRT-PCR results for snRNA U1 and snRNA U5.**
Barplot of the qRT-PCR analysis results for snRNA U1 an snRNA U5 using clutch and adult male samples. This analysis was done for the maternal variants (blue) and somatic variants (red) of each snRNA. The bars are an average of two technical replicates.
