## Supplemental File SF1 for "An alternative spliceosome defined by distinct snRNAs in early zebrafish embryogenesis"

```
                                            20         40         60         80         100        120        140        160
U1consensus  ATACTTACCT GGCAGGGGAG ATACCATGAT CAAGAAGGTG GTTCACCCAG GGCGAGGCTT -GGCCATTGC ACT-CCGGCC ACGCTGACCC CTGCGAATT- CCCCAAATGT GGGAATCTCG ACTGCATAAT TTCTGGTAGT GGGGG-ACTG CGTTCGCGCT CTCCCCTG 164
U1_80_4      ...A...... .......... .......... .......... .......... .......... -......... ...-...... .......... .........- .......... .......... .......... .......... A....-.... ......T... ........ 164
U1_96_4      ...A...... .......... .......... .......... .......... .......... -......... ...-...... .......... .........- .......... .......... .......... .......... A....-.... ......T... ........ 164
U1_91_4      .......... .......... .......... .......... .......... .......... -......... ...-...... .......... .........- .......... .......... .......... .......... A....-.... ......T... ........ 164
U1_85_4      ...A...... .......... .......... .......... .......... .......... -......... ...-...... .......... .........- .......... ........T. .......... .......... A....-.... .......... ........ 164
U1_69_4      ........T. .......... .....G.... .......... .......... .......... -......... ...-...... .......... .........- .......... .......... .......... .......... .....-.... .......... ........ 164
U1_70_4      ........T. .......... .....G.... .......... .......... .......... -......... ...-...... .......... .........- .......... .......... .......... .......... .....-.... .......... ........ 164
U1_124_4     ........T. .......... .......... .....G.... .......... .......... -......... ...-...... .......... .........- .......... .......... .......... .......... .....-.... .......... ........ 164
U1_103_4     .......... .......... .......... .......... .......... .......... -......... ...-...... .......... .........- .......... .......... .......... ....A..... .....-.... .......... ........ 164
U1_119_4     .......... .......... .......... .......... .......... .......... -......... ...-...... .......... .........- .......... .......... .......... ....A.A... .....-.... .......... ........ 164
U1_137_4     .......... .......... .......... .......... .......... .......... -......... ...-...... .......... .........- .......... .......... ..G....... .......... .....-.... .......... ........ 164
U1_145_4     .......... .......... .......... .......... .......... .......... -......... G..-...... .......... .........- .......... .......... ..G....... .......... .....-.... .......... ........ 164
U1_5_4       .......... T......... .......... .......... .......... .......... -......... ...-...... .......... .........- .......... .......... .......... .......... .....-.... .......... ........ 164
U1_22_4      .......... T......... .......... .......... .......... .......... -......... ...-...... .......... .........- .......... .......... .......... .......... .....-.... .......... ........ 164
U1_29_4      .......... T......... .......... .......... .......... .......... -......... ...-...... .......... .........- .......... .......... .......... .......... .....-.... .......... ........ 164
U1_9_4       .......... T......... .......... .......C.. .......... .......... -......... ...-...... .......... .........- .......... .......... .......... .......... .....-.... .......... ........ 164
U1_27_4      .......... T......... .......... .......C.. .......... .......... -......... ...-...... .......... .........- .......... .......... .......... .......... .....-.... .......... ........ 164
U1_35_4      .......... T......... .......... .......C.. .......... .......... -......... ...-...... .......... .........- .......... .......... .......... .......... .....-.... .......... ........ 164
U1_84_4      .......... .......... .......... .......... .......... .......... -......... ...-...... .......... .........- .......... .......... .......... .......... A....-.... .......... ........ 164
U1_105_4     .......... .......... .......... .......... .......... .......... -......... ...-...... .......... .........- .......... .......... .......... .......... A....-.... .......... ........ 164
U1_150_4     .......... T......... .......... .......... .......... .......... -......... ...-...... .......... .........- .......... .......... .......... .......... A....-.... .......... ........ 164
U1_79_4      .......... ....A..... .......... .......... .......... .......... -......... ...-...... .......... .........- .......... .......... .......... .......... .....-.... .......... ........ 164
U1_86_4      .......... ....A..... .......... .......... .......... .......... -......... ...-...... .......... .........- .......... .......... .......... .......... .....-.... .......... ........ 164
U1_94_4      .......... ....A..... .......... .......... .......... .......... -......... ...-...... .......... .........- .......... .......... .......... .......... .....-.... .......... ........ 164
U1_138_4     .......... ....A..... .......... .......... .......... .......... -......... ...-...A.. .......... .........- .......... .......... .......... .......... .....-.... .......... ........ 164
U1_18_4      .......... .......... .......... .......... .......... .......... -......... ...-...... ......G... .........- .......... .......... .......... .......... .....-.... .......... ........ 164
U1_19_4      .......... .......... .......... .......... .......... .......... -......... ...-...... ......G... .........- .......... .......... .......... .......... .....-.... .......... ........ 164
U1_4_4       .......... .......... .......... .......... .......... .......... -......... ...-...... ......G... .........- .......... .......... .......... .......... .....-.... .......... ........ 164
U1_7_4       .......... .......... .......... .......... .......... .......... -......... ...-...... ......G... .........- .......... .......... .......... .......... .....-.... .......... ........ 164
U1_11_4      .......... .......... .......... .......... .......... .......... -......... ...-...... ......G... .........- .......... .......... .......... .......... .....-.... .......... ........ 164
U1_12_4      .......... .......... .......... .......... .......... .......... -......... ...-...... ......G... .........- .......... .......... .......... .......... .....-.... .......... ........ 164
U1_31_4      .......... .......... .......... .......... .......... .......... -......... ...-...... ......G... .........- .......... .......... .......... .......... .....-.... .......... ........ 164
U1_37_4      .......... .......... .......... .......... .......... .......... -......... ...-...... ......G... .........- .......... .......... .......... .......... .....-.... .......... ........ 164
U1_23_4      .......... .......... .......... .......... .......... .......... -......... ...-...... ......G... .........- .......... .......... .......... .......... .....-.... .......... ........ 164
U1_26_4      .......... .......... .......... .......... .......... .......... -......... ...-...... ......G... .........- .......... .......... .......... .......... .....-.... .......... ........ 164
U1_28_4      .......... .......... .......... .......... .......... .......... -......... ...-...... ......G... .........- .......... .......... .......... .......... .....-.... .......... ........ 164
U1_21_4      .......... .......... .......... .......... .......... .......... -......... ...-...... ......G..G .........- .......... .......... .......... .......... .....-.... .......... ........ 164
U1_24_4      .......... .......... .......... .......... .......... .......... -......... ...-...... ......G... .........- .......... .......... .A........ .......... .....-.... .......... ........ 164
U1_30_4      .......... .......... .......... .......... .......... .......... -......... ...-...... ......G... .........- .......... .......... .A........ .......... .....-.... .......... ........ 164
U1_3_4       .......... .......... .......... .......... .......... .......... -......... ...-...... .......... .........- .......... .......... .......... .......... .....-.... .......... .......T 164
U1_6_4       .......... .......... .......... .......... .......... .......... -......... ...-...... .......... .........- .......... .......... .......... .......... .....-.... .......... .......T 164
U1_34_4      .......... .......... .......... .......... .......... .......... -......... ...-...... .......... .........- .......... .......... .......T.. .......... .....-.... .......... ........ 164
U1_92_4      ...A...... .......... .......... .......... .......... .......... -......... ...-...... .......... .........- .......... .......... .......... .......... .....-.... .......... ........ 164
U1_17_4      .......... .......... .......... .......... .......... .......... -......... ...-...... .......... .........- .......... .......... .......... .......... .....-.... .......... ........ 164
U1_20_4      .......... .......... .......... .......... .......... .......... -......... ...-...... .......... .........- .......... .......... .......... .......... .....-.... .......... ........ 164
U1_25_4      .......... .......... .......... .......... .......... .......... -......... ...-...... .......... .........- .......... .......... .......... .......... .....-.... .......... ........ 164
U1_32_4      .......... .......... .......... .......... .......... .......... -......... ...-...... .......... .........- .......... .......... .......... .......... .....-.... .......... ........ 164
U1_33_4      .......... .......... .......... .......... .......... .......... -......... ...-...... .......... .........- .......... .......... .......... .......... .....-.... .......... ........ 164
U1_36_4      .......... .......... .......... .......... .......... .......... -......... ...-...... .......... .........- .......... .......... .......... .......... .....-.... .......... ........ 164
U1_38_4      .......... .......... .......... .......... .......... .......... -......... ...-...... .......... .........- .......... .......... .......... .......... .....-.... .......... ........ 164
U1_40_4      .......... .......... .......... .......... .......... .......... -......... ...-...... .......... .........- .......... .......... .......... .......... .....-.... .......... ........ 164
U1_41_4      .......... .......... .......... .......... .......... .......... -......... ...-...... .......... .........- .......... .......... .......... .......... .....-.... .......... ........ 164
U1_42_4      .......... .......... .......... .......... .......... .......... -......... ...-...... .......... .........- .......... .......... .......... .......... .....-.... .......... ........ 164
U1_43_4      .......... .......... .......... .......... .......... .......... -......... ...-...... .......... .........- .......... .......... .......... .......... .....-.... .......... ........ 164
U1_44_4      .......... .......... .......... .......... .......... .......... -......... ...-...... .......... .........- .......... .......... .......... .......... .....-.... .......... ........ 164
U1_45_4      .......... .......... .......... .......... .......... .......... -......... ...-...... .......... .........- .......... .......... .......... .......... .....-.... .......... ........ 164
U1_46_4      .......... .......... .......... .......... .......... .......... -......... ...-...... .......... .........- .......... .......... .......... .......... .....-.... .......... ........ 164
U1_47_4      .......... .......... .......... .......... .......... .......... -......... ...-...... .......... .........- .......... .......... .......... .......... .....-.... .......... ........ 164
U1_48_4      .......... .......... .......... .......... .......... .......... -......... ...-...... .......... .........- .......... .......... .......... .......... .....-.... .......... ........ 164
U1_49_4      .......... .......... .......... .......... .......... .......... -......... ...-...... .......... .........- .......... .......... .......... .......... .....-.... .......... ........ 164
U1_50_4      .......... .......... .......... .......... .......... .......... -......... ...-...... .......... .........- .......... .......... .......... .......... .....-.... .......... ........ 164
U1_51_4      .......... .......... .......... .......... .......... .......... -......... ...-...... .......... .........- .......... .......... .......... .......... .....-.... .......... ........ 164
U1_52_4      .......... .......... .......... .......... .......... .......... -......... ...-...... .......... .........- .......... .......... .......... .......... .....-.... .......... ........ 164
U1_53_4      .......... .......... .......... .......... .......... .......... -......... ...-...... .......... .........- .......... .......... .......... .......... .....-.... .......... ........ 164
U1_54_4      .......... .......... .......... .......... .......... .......... -......... ...-...... .......... .........- .......... .......... .......... .......... .....-.... .......... ........ 164
U1_55_4      .......... .......... .......... .......... .......... .......... -......... ...-...... .......... .........- .......... .......... .......... .......... .....-.... .......... ........ 164
U1_56_4      .......... .......... .......... .......... .......... .......... -......... ...-...... .......... .........- .......... .......... .......... .......... .....-.... .......... ........ 164
U1_57_4      .......... .......... .......... .......... .......... .......... -......... ...-...... .......... .........- .......... .......... .......... .......... .....-.... .......... ........ 164
U1_58_4      .......... .......... .......... .......... .......... .......... -......... ...-...... .......... .........- .......... .......... .......... .......... .....-.... .......... ........ 164
U1_59_4      .......... .......... .......... .......... .......... .......... -......... ...-...... .......... .........- .......... .......... .......... .......... .....-.... .......... ........ 164
U1_60_4      .......... .......... .......... .......... .......... .......... -......... ...-...... .......... .........- .......... .......... .......... .......... .....-.... .......... ........ 164
U1_61_4      .......... .......... .......... .......... .......... .......... -......... ...-...... .......... .........- .......... .......... .......... .......... .....-.... .......... ........ 164
U1_65_4      .......... .......... .......... .......... .......... .......... -......... ...-...... .......... .........- .......... .......... .......... .......... .....-.... .......... ........ 164
U1_68_4      .......... .......... .......... .......... .......... .......... -......... ...-...... .......... .........T .......... .......... .......... .......... .....-.... .......... ........ 165
U1_72_4      .......... .......... .......... .......... .......... .......... -......... ...-...... .......... .........- .......... .......... .......... .......... .....-.... .......... ........ 164
U1_73_4      .......... .......... .......... .......... .......... .......... -......... ...-...... .......... .........- .......... .......... .......... .......... .....-.... .......... ........ 164
U1_74_4      .......... .......... .......... .......... .......... .......... -......... ...-...... .......... .........- .......... .......... .......... .......... .....-.... .......... ........ 164
U1_75_4      .......... .......... .......... .......... .......... .......... -......... ...-...... .......... .........- .......... .......... .......... .......... .....-.... .......... ........ 164
U1_76_4      .......... .......... .......... .......... .......... .......... -......... ...-...... .......... .........- .......... .......... .......... .......... .....-.... .......... ........ 164
U1_81_4      .......... .......... .......... .......... .......... .......... -......... ...-...... .......... .........- .......... .......... .......... .......... .....-.... .......... ........ 164
U1_87_4      .......... .......... .......... .......... .......... .......... -......... ...-...... .......... .........- .......... .......... .......... .......... .....-.... .......... ........ 164
U1_88_4      .......... .......... .......... .......... .......... .......... -......... ...-...... .......... .........- .......... .......... .......... .......... .....-.... .......... ........ 164
U1_89_4      .......... .......... .......... .......... .......... .......... -......... ...-...... .......... .........- .......... .......... .......... .......... .....-.... .......... ........ 164
U1_97_4      .......... .......... .......... .......... .......... .......... -......... ...-...... .......... .........- .......... .......... .......... .......... .....-.... .......... ........ 164
U1_98_4      .......... .......... .......... .......... .......... .......... -......... ...-...... .......... .........- .......... .......... .......... .......... .....-.... .......... ........ 164
U1_101_4     .......... .......... .......... .......... .......... .......... -......... ...-...... .......... .........- .......... .......... .......... .......... .....-.... .......... ........ 164
U1_106_4     .......... .......... .......... .......... .......... .......... -......... ...-...... .......... .........- .......... .......... .......... .......... .....-.... .......... ........ 164
U1_125_4     .......... .......... .......... .......... .......... .......... -......... ...-...... .......... .........- .......... .......... .......... .......... .....-.... .......... ........ 164
U1_131_4     .......... .......... .......... .......... .......... .......... -......... ...-...... .......... .........- .......... .......... .......... .......... .....-.... .......... ........ 164
U1_132_4     .......... .......... .......... .......... .......... .......... -......... ...-...... .......... .........- .......... .......... .......... .......... .....-.... .......... ........ 164
U1_134_4     .......... .......... .......... .......... .......... .......... -......... ...-...... .......... .........- .......... .......... .......... .......... .....-.... .......... ........ 164
```

```
                    20         40         60         80        100        120        140        160
U1_135_4   .......... .......... .......... .......... .......... .......... -......... ...-...... .......... .........- .......... .......... .......... .......... .....-.... .......... ........ 164
U1_136_4   .......... .......... .......... .......... .......... .......... -......... ...-...... .......... .........- .......... .......... .......... .......... .....-.... .......... ........ 164
U1_139_4   .......... .......... .......... .......... .......... .......... -......... ...-...... .......... .........- .......... .......... .......... .......... .....-.... .......... ........ 164
U1_140_4   .......... .......... .......... .......... .......... .......... -......... ...-...... .......... .........- .......... .......... .......... .......... .....-.... .......... ........ 164
U1_142_4   .......... .......... .......... .......... .......... .......... -......... ...-...... .......... .........- .......... .......... .......... .......... .....-.... .......... ........ 164
U1_143_4   .......... .......... .......... .......... .......... .......... -......... ...-...... .......... .........- .......... .......... .......... .......... .....-.... .......... ........ 164
U1_144_4   .......... .......... .......... .......... .......... .......... -......... ...-...... .......... .........- .......... .......... .......... .......... .....-.... .......... ........ 164
U1_147_4   .......... .......... .......... .......... .......... .......... -......... ...-...... .......... .........- .......... .......... .......... .......... .....-.... .......... ........ 164
U1_148_4   .......... .......... .......... .......... .......... .......... -......... ...-...... .......... .........- .......... .......... .......... .......... .....-.... .......... ........ 164
U1_149_4   .......... .......... .......... .......... .......... .......... -......... ...-...... .......... .........- .......... .......... .......... .......... .....-.... .......... ........ 164
U1_154_4   .......... .......... .......... .......... .......... .......... -......... ...-...... .......... .........- .......... .......... .......... .......... .....-.... .......... ........ 164
U1_158_4   .......... .......... .......... .......... .......... .......... -......... ...-...... .......... .........- .......... .......... .......... .......... .....-.... .......... ........ 164
U1_179_4   .......... .......... .......... .......... .......... .......... -......... ...-...... .......... .........- .......... .......... .......... .......... .....-.... .......... ........ 164
U1_8_4     .......... .......... .......... .......... .......... .......... -......... ...-...... .......... .........- .......... .......... .......... .......... .....-.... .......... ........ 164
U1_10_4    .......... .......... .......... .......... .......... .......... -......... ...-...... .......... .........- .......... .......... .......... .......... .....-.... .......... ........ 164
U1_13_4    .......... .......... .......... .......... .......... .......... -......... ...-...... .......... .........- .......... .......... .......... .......... .....-.... .......... ........ 164
U1_14_4    .......... .......... .......... .......... .......... .......... -......... ...-...... .......... .........- .......... .......... .......... .......... .....-.... .......... ........ 164
U1_15_4    .......... .......... .......... .......... .......... .......... -......... ...-...... .......... .........- .......... .......... .......... .......... .....-.... .......... ........ 164
U1_16_4    .......... .......... .......... .......... .......... .......... -......... ...-...... .......... .........- .......... .......... .......... .......... .....-.... .......... ........ 164
U1_198_4   .......... .......... .......... .......... .......... .......... -......... ...-...... .......... .........- .......... .......... .......... .......... .....-.... .......... ........ 164
U1_199_4   .......... .......... .......... .......... .......... .......... -......... ...-...... .......... .........- .......... .......... .......... .......... .....-.... .......... ........ 164
U1_194_4   .......... .......... .......... .......... .......... .......... -......... ...-...... .......... .........- .......... .......... .......... .......... .....-.... .......... ........ 164
U1_195_4   .......... .......... .......... .......... .......... .......... -......... ...-...... .......... .........- .......... .......... .......... .......... .....-.... .......... ........ 164
U1_196_4   .......... .......... .......... .......... .......... .......... -......... ...-...... .......... .........- .......... .......... .......... .......... .....-.... .......... ........ 164
U1_197_4   .......... .......... .......... .......... .......... .......... -......... ...-...... .......... .........- .......... .......... .......... .......... .....-.... .......... ........ 164
U1_182_4   .......... .......... .......... .......... .......... .......... -......... ...-...... .......... .........- .......... .......... .......... .......... .....-.... .......... ........ 164
U1_183_4   .......... .......... .......... .......... .......... .......... -......... ...-...... .......... .........- .......... .......... .......... .......... .....-.... .......... ........ 164
U1_184_4   .......... .......... .......... .......... .......... .......... -......... ...-...... .......... .........- .......... .......... .......... .......... .....-.... .......... ........ 164
U1_185_4   .......... .......... .......... .......... .......... .......... -......... ...-...... .......... .........- .......... .......... .......... .......... .....-.... .......... ........ 164
U1_186_4   .......... .......... .......... .......... .......... .......... -......... ...-...... .......... .........- .......... .......... .......... .......... .....-.... .......... ........ 164
U1_189_4   .......... .......... .......... .......... .......... .......... -......... ...-...... .......... .........- .......... .......... .......... .......... .....-.... .......... ........ 164
U1_190_4   .......... .......... .......... .......... .......... .......... -......... ...-...... .......... .........- .......... .......... .......... .......... .....-.... .......... ........ 164
U1_193_4   .......... .......... .......... .......... .......... .......... -......... ...-...... .......... .........- .......... .......... .......... .......... .....-.... .......... ........ 164
U1_212_4   .......... .......... .......... .......... .......... .......... -......... ...-...... .......... .........- .......... .......... .......... .......... .....-.... .......... ........ 164
U1_215_4   .......... .......... .......... .......... .......... .......... -......... ...-...... .......... .........- .......... .......... .......... .......... .....-.... .......... ........ 164
U1_208_4   .......... .......... .......... .......... .......... .......... -......... ...-...... .......... .........- .......... .......... .......... .......... .....-.... .......... ........ 164
U1_209_4   .......... .......... .......... .......... .......... .......... -......... ...-...... .......... .........- .......... .......... .......... .......... .....-.... .......... ........ 164
U1_210_4   .......... .......... .......... .......... .......... .......... -......... ...-...... .......... .........- .......... .......... .......... .......... .....-.... .......... ........ 164
U1_211_4   .......... .......... .......... .......... .......... .......... -......... ...-...... .......... .........- .......... .......... .......... .......... .....-.... .......... ........ 164
U1_200_4   .......... .......... .......... .......... .......... .......... -......... ...-...... .......... .........- .......... .......... .......... .......... .....-.... .......... ........ 164
U1_201_4   .......... .......... .......... .......... .......... .......... -......... ...-...... .......... .........- .......... .......... .......... .......... .....-.... .......... ........ 164
U1_202_4   .......... .......... .......... .......... .......... .......... -......... ...-...... .......... .........- .......... .......... .......... .......... .....-.... .......... ........ 164
U1_203_4   .......... .......... .......... .......... .......... .......... -......... ...-...... .......... .........- .......... .......... .......... .......... .....-.... .......... ........ 164
U1_204_4   .......... .......... .......... .......... .......... .......... -......... ...-...... .......... .........- .......... .......... .......... .......... .....-.... .......... ........ 164
U1_205_4   .......... .......... .......... .......... .......... .......... -......... ...-...... .......... .........- .......... .......... .......... .......... .....-.... .......... ........ 164
U1_206_4   .......... .......... .......... .......... .......... .......... -......... ...-...... .......... .........- .......... .......... .......... .......... .....-.... .......... ........ 164
U1_207_4   .......... .......... .......... .......... .......... .......... -......... ...-...... .......... .........- .......... .......... .......... .......... .....-.... .......... ........ 164
U1_180_4   .......... .......... .......... .......... .......... .......... -......... ...-...... .......... .........- .......... .......... .......... .......... .....-.... .......... ........ 164
U1_181_4   .......... .......... .......... .......... .......... .......... -......... ...-...... .......... .........- .......... .......... .......... .......... .....-.... .......... ........ 164
U1_133_4   .......... .......... .......... .......... .......... .......... -......... ...-...... .......... .........- .......... .......... .......... .......... ...T.-.... .......... ........ 164
U1_259_20  T......... .......... .......... .......... .......... .......... -......... ...-...... .......... .........- .......... .......... .......... .......... .....-.... .......... ........ 164
U1_99_4    .......... .......... .......... .......... .......... .......... -......... ...-..A... .......... .........- .......... .......... .......... .......... .....-.... .......... ........ 164
U1_121_4   .......... .......... .......... .......... .......... .......... -......... ...-...... .......... .........- .......... .......... .......... .......... .....-.T.. .......... ........ 164
U1_213_4   .......... .......... .......... .......... .......... .......... -......... ...-...... .......... .....T...- .......... .......... .......... .......... .....-.... .......... ........ 164
U1_130_4   .......... .......... .......... .......... .......... ....A..... -......... ...-...... .......... .........- .......... .......... .......... .......... .....-.... .......... ........ 164
U1_258_20  .......... .......... .......... .......... .......... ....A..... -......... ...-...... .......... .........- .......... .......... .......... .......... .....-.... .......... ........ 164
U1_187_4   .......... .......... .......... .......... .....C.... .......... -......... ...-...... .......... .........- .......... .......... .......... .......... .....G.... .......... ........ 165
U1_188_4   .......... .......... .......... .......... .....C.... .......... -......... ...-...... .......... .........- .......... .......... .......... .......... .....-.... .......... ........ 164
U1_122_4   .......... .......... .......... .......... .......... .......... -......... ...-...... .......... .........- .......... ........A. .......... .......... .....-.... .......... ........ 164
U1_126_4   .......... .......... .......... .......... .......... .......... -......... ...-...... .......... .........- .......... ........A. .......... .......... .....-.... .......... ........ 164
U1_127_4   .......... .......... .......... .......... .......... .......... -......... ...-...... .......... .........- .......... ........A. .......... .......... .....-.... .......... ........ 164
U1_151_4   .......... .......... .......... .......... .......T.. .......... -......... ...-...... .......... .........- .......... .......... .......... .......... .....-.... .......... ........ 164
U1_156_4   .......... .......... .......... .......... .......T.. .......... -......... ...-...... .......... .........- .......... .......... .......... .......... .....-.... .......... ........ 164
U1_157_4   .......... .......... .......... .......... .......T.. .......... -......... ...-...... .......... .........- .......... .......... .......... .......... .....-.... .......... ........ 164
U1_164_4   .......... .......... .......... .......... .......T.. .......... -......... ...-...... .......... .........- .......... .......... .......... .......... .....-.... .......... ........ 164
U1_166_4   .......... .......... .......... .......... .......T.. .......... -......... ...-...... .......... .........- .......... .......... .......... .......... .....-.... .......... ........ 164
U1_169_4   .......... .......... .......... .......... .......T.. .......... -......... ...-...... .......... .........- .......... .......... .......... .......... .....-.... .......... ........ 164
U1_175_4   .......... .......... .......... .......... .......T.. .......... -......... ...-...... .......... .........- .......... .......... .......... .......... .....-.... .......... ........ 164
U1_176_4   .......... .......... .......... .......... .......T.. .......... -......... ...-...... .......... .........- .......... .......... .......... .......... .....-.... .......... ........ 164
U1_223_4   .......... .......... .......... .......... .......... .......... -......... ...-...... .......... .........- .......... .......... .......... .......... .....-.... ....T..... ........ 164
U1_224_4   .......... .......... .......... .......... .......... .......... -......... ...-...... .......... .........- .......... .......... .......... .......... .....-.... ....T..... ........ 164
U1_225_4   .......... .......... .......... .......... .......... .......... -......... ...-...... .......... .........- .......... .......... .......... .......... .....-.... ....T..... ........ 164
U1_226_4   .......... .......... .......... .......... .......... .......... -......... ...-...... .......... .........- .......... .......... .......... .......... .....-.... ....T..... ........ 164
U1_227_4   .......... .......... .......... .......... .......... .......... -......... ...-...... .......... .........- .......... .......... .......... .......... .....-.... ....T..... ........ 164
U1_228_4   .......... .......... .......... .......... .......... .......... -......... ...-...... .......... .........- .......... .......... .......... .......... .....-.... ....T..... ........ 164
U1_229_4   .......... .......... .......... .......... .......... .......... -......... ...-...... .......... .........- .......... .......... .......... .......... .....-.... ....T..... ........ 164
U1_230_4   .......... .......... .......... .......... .......... .......... -......... ...-...... .......... .........- .......... .......... .......... .......... .....-.... ....T..... ........ 164
U1_231_4   .......... .......... .......... .......... .......... .......... -......... ...-...... .......... .........- .......... .......... .......... .......... .....-.... ....T..... ........ 164
U1_232_4   .......... .......... .......... .......... .......... .......... -......... ...-...... .......... .........- .......... .......... .......... .......... .....-.... ....T..... ........ 164
U1_233_4   .......... .......... .......... .......... .......... .......... -......... ...-...... .......... .........- .......... .......... .......... .......... .....-.... ....T..... ........ 164
U1_234_4   .......... .......... .......... .......... .......... .......... -......... ...-...... .......... .........- .......... .......... .......... .......... .....-.... ....T..... ........ 164
U1_214_4   .......... .......... .......... .......... .......... .......... -......... ...-...... .......... .........- .......... .......... .......... .......... .....-.... ....T..... ........ 164
U1_216_4   .......... .......... .......... .......... .......... .......... -......... ...-...... .......... .........- .......... .......... .......... .......... .....-.... ....T..... ........ 164
U1_217_4   .......... .......... .......... .......... .......... .......... -......... ...-...... .......... .........- .......... .......... .......... .......... .....-.... ....T..... ........ 164
U1_218_4   .......... .......... .......... .......... .......... .......... -......... ...-...... .......... .........- .......... .......... .......... .......... .....-.... ....T..... ........ 164
U1_219_4   .......... .......... .......... .......... .......... .......... -......... ...-...... .......... .........- .......... .......... .......... .......... .....-.... ....T..... ........ 164
U1_220_4   .......... .......... .......... .......... .......... .......... -......... ...-...... .......... .........- .......... .......... .......... .......... .....-.... ....T..... ........ 164
U1_221_4   .......... .......... .......... .......... .......... .......... -......... ...-...... .......... .........- .......... .......... .......... .......... .....-.... ....T..... ........ 164
U1_222_4   .......... .......... .......... .......... .......... .......... -......... ...-...... .......... .........- .......... .......... .......... .......... .....-.... ....T..... ........ 164
U1_241_4   .......... .......... .......... .......... .......... .......... -......... ...-...... .......... .........- .......... .......... .......... .......... .....-.... ....T..... ........ 164
U1_242_4   .......... .......... .......... .......... .......... .......... -......... ...-...... .......... .........- .......... .......... .......... .......... .....-.... ....T..... ........ 164
```

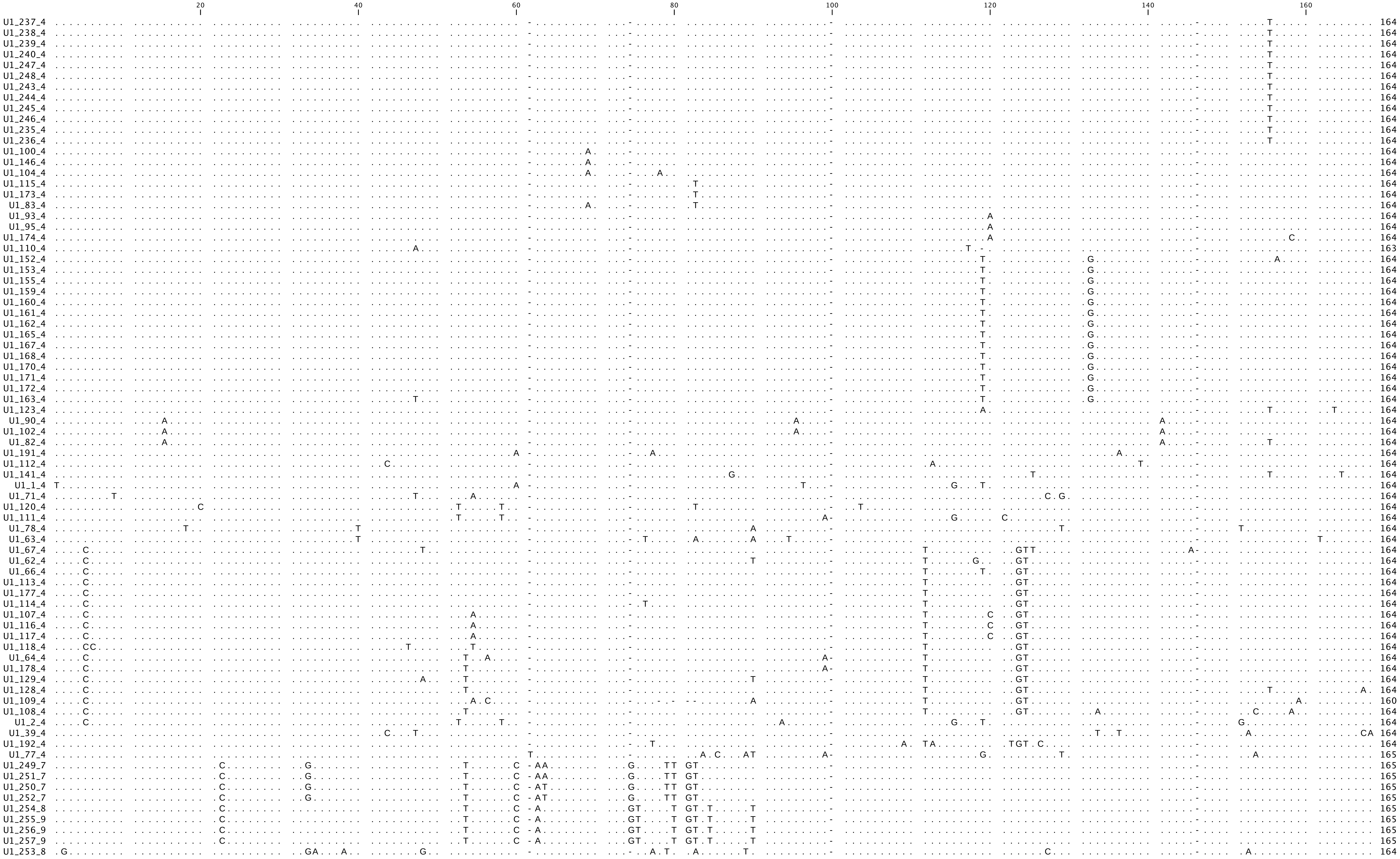
20
40
60
80
100
120
140
160
U1_237_4 164
U1_238_4 164
U1_239_4 164
U1_240_4 164
U1_247_4 164
U1_248_4 164
U1_243_4 164
U1_244_4 164
U1_245_4 164
U1_246_4 164
U1_235_4 164
U1_236_4 164
U1_100_4 164
U1_146_4 164
U1_104_4 164
U1_115_4 164
U1_173_4 164
U1_83_4 164
U1_93_4 164
U1_95_4 164
U1_174_4 164
U1_110_4 163
U1_152_4 164
U1_153_4 164
U1_155_4 164
U1_159_4 164
U1_160_4 164
U1_161_4 164
U1_162_4 164
U1_165_4 164
U1_167_4 164
U1_168_4 164
U1_170_4 164
U1_171_4 164
U1_172_4 164
U1_163_4 164
U1_123_4 164
U1_90_4 164
U1_102_4 164
U1_82_4 164
U1_191_4 164
U1_112_4 164
U1_141_4 164
U1_1_4 164
U1_71_4 164
U1_120_4 164
U1_111_4 164
U1_78_4 164
U1_63_4 164
U1_67_4 164
U1_62_4 164
U1_66_4 164
U1_113_4 164
U1_177_4 164
U1_114_4 164
U1_107_4 164
U1_116_4 164
U1_117_4 164
U1_118_4 164
U1_64_4 164
U1_178_4 164
U1_129_4 164
U1_128_4 164
U1_109_4 160
U1_108_4 164
U1_2_4 164
U1_39_4 164
U1_192_4 164
U1_77_4 165
U1_249_7 165
U1_251_7 165
U1_250_7 165
U1_252_7 165
U1_254_8 165
U1_255_9 165
U1_256_9 165
U1_257_9 165
U1_253_8 164

```
                                              20                   40                   60                   80                   100                  120                  140                  160                  180
                                               |                    |                    |                    |                    |                    |                    |                    |                    |
U2consensus ATCGCTTCTC GGCCTTTTGG CTAAGATCAA GTGTAGTATC TGTTCTTATC AGTTTAATAT CTGATACGTG CCCTACCCGG GCACCATATA TTAAATTGAT TTTTGGAACA GGGAGATGGA ATAGGGGCTT GCTCCGTCCA CTCCACGCAT CGACCCGGTA TTGCAGTACC TCCGGGAACG GTGCACCCCC T 191
   U2_6_13  .......... .......... .......... .......... .......... .......... .......... .......... .......... .......... .......... .......... .......... .......... .......... .......... .......... .......... .......... . 191
   U2_7_13  .......... .......... .......... .......... .......... .......... .......... .......... .......... .......... .......... .......... .......... .......... .......... .......... .......... .......... .......... . 191
    U2_5_7  .......... .......... .......... .......... .......... .......... .......... .......... .......... .......... .......... .......... .......... .......... .......T.. .......... .........A .......... .......... . 191
    U2_3_6  .......... .......... .......... .......... .......... .......... .......... .......... .......... .......... .......G.. .......... .......... .......... .......... .......... .........T .......... .......... . 191
    U2_4_7  .......... .......... .......... .......... .......... .......... .......... .......... .......... .......... ........T. .......... .......... .......... .......... .......... .........T .......... .......... . 191
    U2_1_2  .......... .......... .......... .......... .......... .......... .......... .......... ....T..... .......... .....C.G.T .........G T......... .....AC... .......... .....T.... .......... .......... ......T.AT . 191
    U2_2_2  .......... .......... .......... .......... .......... .......... .......... .......... ....T..... .......... .....C.G.T .........G T......... .....AC... .......... .....T.... .......... .......... ......T.AT . 191
```

```
                                     20         40         60         80         100        120        140
U4consensus ATTCTTTCGC AGAGGCAATA TCATAGCCTA TGAGGCCTAG CCGAGGCGTG ATCATTGCTG GTTGAAAACT TTACCCAATA CCCCGCCGAG ATGGCTTGAA ATACAGTCAG CTCTGGCAAT TTTTGACAGC CTCCTAGAAG G 141
U4_7_17     .......... .......... .......... .......... ......T... .......... .......C.. .......... .......... .......... .......... ........G. .......... .......... . 141
U4_3_17     .......... .......... .......... .......... ......T... .......... .......... .......... .......... .......... .......... .......... .......... .......... . 141
U4_24_17    .......... .......... .......... .......... ......T... .......... .......... .......... .......... .......... .......... ........G. .......... .......... . 141
U4_6_17     .......... .......... ......T... .......... .......... .......... .......... .......... .......... .......... .......... .......... .......... .......... . 141
U4_4_17     .......... .......... .......... .......... .......... .......... .......... .......... .......... .......... .......... .......... .......... .......... . 141
U4_5_17     .......... .......... .......... .......... .......... .......... .......... .......... .......... .......... .......... .......... .......... .......... . 141
U4_8_17     .......... .......... .......... .......... .......... .......... .......... .......... .......... .......... .......... .......... .......... .......... . 141
U4_10_17    .......... .......... .......... .......... .......... .......... .......... .......... .......... .......... .......... .......... .......... .......... . 141
U4_14_17    .......... .......... .......... .......... .......... .......... .......... .......... .......... .......... .......... .......... .......... .......... . 141
U4_17_17    .......... .......... .......... .......... .......... .......... .......... .......... .......... .......... .......... .......... .......... .......... . 141
U4_18_17    .......... .......... .......... .......... .......... .......... .......... .......... .......... .......... .......... .......... .......... .......... . 141
U4_19_17    .......... .......... .......... .......... .......... .......... .......... .......... .......... .......... .......... .......... .......... .......... . 141
U4_20_17    .......... .......... .......... .......... .......... .......... .......... .......... .......... .......... .......... .......... .......... .......... . 141
U4_21_17    .......... .......... .......... .......... .......... .......... .......... .......... .......... .......... .......... .......... .......... .......... . 141
U4_22_17    .......... .......... .......... .......... .......... .......... .......... .......... .......... .......... .......... .......... .......... .......... . 141
U4_12_17    .......... .......... .......... .......... .......... .......... T......... .......... .......... .......... .......... .......... .......... .......... . 141
U4_16_17    .......... .......... .......T.. .......... .......... .......... .......... .......... .......... .......... .......... .......... .......... .......... . 141
U4_23_17    .......... .......... .......T.. .......... .......... .......... .......... .......... .......... .......... .......... .......... .......... .......... . 141
U4_9_17     .......... .......... .......... .......... .......... .......... .......... ..G....... ......T... .......... .......... .......... .......... .......... . 141
U4_13_17    .......... .........G .......... .......... .......... .......... .......... ..G....... ......T... .......... .......... .......... .......... .......... . 141
U4_11_17    .......... .A........ .......... .......... .......... .......... .......... .......... ...T...... ...T...... .......... .......... .......... .......... . 141
U4_15_17    .......... .A........ .......... .......... .......... .......... .......... .......... ...T...... ...T...... .......... .......... .......... .......... . 141
U4_25_17    ...T...... .......... .......... .......... .T...T.... .......... .......... .......... .......... .........G .......... .......... .......... .......... . 141
U4_26_25    .GCT..G... ..T....G.. ..G....... .....TT..T ........C. ..T......A A......... .......... ........T. .C.A...... ...T....G. .A........ .........T ..T..C.... A 141
U4_1_4      .GCT..G... ..T....G.. ..G....... .....TT..T ........C. ..T......A A......... .......... ........T. .C.A...... ...T....G. .A........ .......G.T ...GACAG.. A 141
U4_2_4      ..C...G... ..T....G.. ..G.....C. .....TT..T ........C. ..T....... .......... .......... ........T. .C.A...... ...T....G. .......... .......G.T ...AA.AG.. A 141
```

```
                           20         40         60         80         100
                            |          |          |          |          |
U5consensus -CAGCTCGAG TTTCTCTTCA ATCACGCACA AATCTTTCGC CTTTTACTAA AGATTTCCGT GGGGAGGAAC TATTGTGAGT TTTGTTTATT TTTGGGTGCT CTGCTATCTG CAGAGCT- 116
U5_11_5     -......... .......... .......... .......... .......... .......... .......... .......... .......... .......... .......... .......- 116
U5_12_5     -......... .......... .......... .......... .......... .......... .......... .......... .......... .......... .......... .......- 116
U5_13_5     -......... .......... .......... .......... .......... .......... .......... .......... .......... .......... .......... .......- 116
U5_14_5     -......... .......... .......... .......... .......... .......... .......... .......... .......... .......... .......... .......- 116
U5_15_5     -......... .......... .......... .......... .......... .......... .......... .......... .......... .......... .......... .......- 116
U5_16_5     -......... .......... .......... .......... .......... .......... .......... .......... .......... .......... .......... .......- 116
U5_17_5     -......... .......... .......... .......... .......... .......... .......... .......... .......... .......... .......... .......- 116
U5_18_5     -......... .......... .......... .......... .......... .......... .......... .......... .......... .......... .......... .......- 116
U5_19_5     -......... .......... .......... .......... .......... .......... .......... .......... .......... .......... .......... .......- 116
U5_20_5     -......... .......... .......... .......... .......... .......... .......... .......... .......... .......... .......... .......- 116
U5_21_5     -......... .......... .......... .......... .......... .......... .......... .......... .......... .......... .......... .......- 116
U5_22_5     -......... .......... .......... .......... .......... .......... .......... .......... .......... .......... .......... .......- 116
U5_23_5     -......... .......... .......... .......... .......... .......... .......... .......... .......... .......... .......... .......- 116
U5_24_5     -......... .......... .......... .......... .......... .......... .......... .......... .......... .......... .......... .......- 116
U5_25_5     -......... .......... .......... .......... .......... .......... .......... .......... .......... .......... .......... .......- 116
U5_26_5     -......... .......... .......... .......... .......... .......... .......... .......... .......... .......... .......... .......- 116
U5_5_5      -......... .......... .......... .......... .......... .......... .......... .......... .......... .......... .......... .......- 116
U5_6_5      -......... .......... .......... .......... .......... .......... .......... .......... .......... .......... .......... .......- 116
U5_7_5      -......... .......... .......... .......... .......... .......... .......... .......... .......... .......... .......... .......- 116
U5_8_5      -......... .......... .......... .......... .......... .......... .......... .......... .......... .......... .......... .......- 116
U5_131_5    -......... .......... .......... .......... .......... .......... .......... .......... .......... .......... .......... .......- 116
U5_126_5    -......... .......... .......... .......... .......... .......... .......... .......... .......... .......... .......... .......- 116
U5_127_5    -......... .......... .......... .......... .......... .......... .......... .......... .......... .......... .......... .......- 116
U5_128_5    -......... .......... .......... .......... .......... .......... .......... .......... .......... .......... .......... .......- 116
U5_129_5    -......... .......... .......... .......... .......... .......... .......... .......... .......... .......... .......... .......- 116
U5_27_5     -......... .......... .......... .......... .......... .......... .......... .......... .......... .......... .......... .......- 116
U5_28_5     -......... .......... .......... .......... .......... .......... .......... .......... .......... .......... .......... .......- 116
U5_29_5     -......... .......... .......... .......... .......... .......... .......... .......... .......... .......... .......... .......- 116
U5_30_5     -......... .......... .......... .......... .......... .......... .......... .......... .......... .......... .......... .......- 116
U5_31_5     -......... .......... .......... .......... .......... .......... .......... .......... .......... .......... .......... .......- 116
U5_32_5     -......... .......... .......... .......... .......... .......... .......... .......... .......... .......... .......... .......- 116
U5_33_5     -......... .......... .......... .......... .......... .......... .......... .......... .......... .......... .......... .......- 116
U5_34_5     -......... .......... .......... .......... .......... .......... .......... .......... .......... .......... .......... .......- 116
U5_35_5     -......... .......... .......... .......... .......... .......... .......... .......... .......... .......... .......... .......- 116
U5_36_5     -......... .......... .......... .......... .......... .......... .......... .......... .......... .......... .......... .......- 116
U5_37_5     -......... .......... .......... .......... .......... .......... .......... .......... .......... .......... .......... .......- 116
U5_38_5     -......... .......... .......... .......... .......... .......... .......... .......... .......... .......... .......... .......- 116
U5_39_5     -......... .......... .......... .......... .......... .......... .......... .......... .......... .......... .......... .......- 116
U5_40_5     -......... .......... .......... .......... .......... .......... .......... .......... .......... .......... .......... .......- 116
U5_41_5     -......... .......... .......... .......... .......... .......... .......... .......... .......... .......... .......... .......- 116
U5_42_5     -......... .......... .......... .......... .......... .......... .......... .......... .......... .......... .......... .......- 116
U5_43_5     -......... .......... .......... .......... .......... .......... .......... .......... .......... .......... .......... .......- 116
U5_44_5     -......... .......... .......... .......... .......... .......... .......... .......... .......... .......... .......... .......- 116
U5_45_5     -......... .......... .......... .......... .......... .......... .......... .......... .......... .......... .......... .......- 116
U5_46_5     -......... .......... .......... .......... .......... .......... .......... .......... .......... .......... .......... .......- 116
U5_47_5     -......... .......... .......... .......... .......... .......... .......... .......... .......... .......... .......... .......- 116
U5_48_5     -......... .......... .......... .......... .......... .......... .......... .......... .......... .......... .......... .......- 116
U5_49_5     -......... .......... .......... .......... .......... .......... .......... .......... .......... .......... .......... .......- 116
U5_50_5     -......... .......... .......... .......... .......... .......... .......... .......... .......... .......... .......... .......- 116
U5_51_5     -......... .......... .......... .......... .......... .......... .......... .......... .......... .......... .......... .......- 116
U5_52_5     -......... .......... .......... .......... .......... .......... .......... .......... .......... .......... .......... .......- 116
U5_53_5     -......... .......... .......... .......... .......... .......... .......... .......... .......... .......... .......... .......- 116
U5_54_5     -......... .......... .......... .......... .......... .......... .......... .......... .......... .......... .......... .......- 116
U5_55_5     -......... .......... .......... .......... .......... .......... .......... .......... .......... .......... .......... .......- 116
U5_56_5     -......... .......... .......... .......... .......... .......... .......... .......... .......... .......... .......... .......- 116
U5_57_5     -......... .......... .......... .......... .......... .......... .......... .......... .......... .......... .......... .......- 116
U5_58_5     -......... .......... .......... .......... .......... .......... .......... .......... .......... .......... .......... .......- 116
U5_59_5     -......... .......... .......... .......... .......... .......... .......... .......... .......... .......... .......... .......- 116
U5_60_5     -......... .......... .......... .......... .......... .......... .......... .......... .......... .......... .......... .......- 116
U5_61_5     -......... .......... .......... .......... .......... .......... .......... .......... .......... .......... .......... .......- 116
U5_62_5     -......... .......... .......... .......... .......... .......... .......... .......... .......... .......... .......... .......- 116
U5_63_5     -......... .......... .......... .......... .......... .......... .......... .......... .......... .......... .......... .......- 116
U5_64_5     -......... .......... .......... .......... .......... .......... .......... .......... .......... .......... .......... .......- 116
U5_65_5     -......... .......... .......... .......... .......... .......... .......... .......... .......... .......... .......... .......- 116
U5_66_5     -......... .......... .......... .......... .......... .......... .......... .......... .......... .......... .......... .......- 116
U5_67_5     -......... .......... .......... .......... .......... .......... .......... .......... .......... .......... .......... .......- 116
U5_68_5     -......... .......... .......... .......... .......... .......... .......... .......... .......... .......... .......... .......- 116
U5_69_5     -......... .......... .......... .......... .......... .......... .......... .......... .......... .......... .......... .......- 116
U5_70_5     -......... .......... .......... .......... .......... .......... .......... .......... .......... .......... .......... .......- 116
U5_71_5     -......... .......... .......... .......... .......... .......... .......... .......... .......... .......... .......... .......- 116
U5_72_5     -......... .......... .......... .......... .......... .......... .......... .......... .......... .......... .......... .......- 116
U5_73_5     -......... .......... .......... .......... .......... .......... .......... .......... .......... .......... .......... .......- 116
U5_74_5     -......... .......... .......... .......... .......... .......... .......... .......... .......... .......... .......... .......- 116
U5_75_5     -......... .......... .......... .......... .......... .......... .......... .......... .......... .......... .......... .......- 116
U5_77_5     -......... .......... .......... .......... .......... .......... .......... .......... .......... .......... .......... .......- 116
U5_78_5     -......... .......... .......... .......... .......... .......... .......... .......... .......... .......... .......... .......- 116
U5_80_5     -......... .......... .......... .......... .......... .......... .......... .......... .......... .......... .......... .......- 116
U5_81_5     -......... .......... .......... .......... .......... .......... .......... .......... .......... .......... .......... .......- 116
U5_83_5     -......... .......... .......... .......... .......... .......... .......... .......... .......... .......... .......... .......- 116
U5_84_5     -......... .......... .......... .......... .......... .......... .......... .......... .......... .......... .......... .......- 116
U5_85_5     -......... .......... .......... .......... .......... .......... .......... .......... .......... .......... .......... .......- 116
U5_86_5     -......... .......... .......... .......... .......... .......... .......... .......... .......... .......... .......... .......- 116
U5_87_5     -......... .......... .......... .......... .......... .......... .......... .......... .......... .......... .......... .......- 116
U5_88_5     -......... .......... .......... .......... .......... .......... .......... .......... .......... .......... .......... .......- 116
U5_89_5     -......... .......... .......... .......... .......... .......... .......... .......... .......... .......... .......... .......- 116
U5_90_5     -......... .......... .......... .......... .......... .......... .......... .......... .......... .......... .......... .......- 116
U5_91_5     -......... .......... .......... .......... .......... .......... .......... .......... .......... .......... .......... .......- 116
U5_92_5     -......... .......... .......... .......... .......... .......... .......... .......... .......... .......... .......... .......- 116
U5_93_5     -......... .......... .......... .......... .......... .......... .......... .......... .......... .......... .......... .......- 116
U5_94_5     -......... .......... .......... .......... .......... .......... .......... .......... .......... .......... .......... .......- 116
```

```
                                  20         40         60         80         100
                                   |          |          |          |          |
  U5_95_5 -......... .......... .......... .......... .......... .......... .......... .......... .......... .......... .......... .......-  116
  U5_96_5 -......... .......... .......... .......... .......... .......... .......... .......... .......... .......... .......... .......-  116
  U5_97_5 -......... .......... .......... .......... .......... .......... .......... .......... .......... .......... .......... .......-  116
  U5_98_5 -......... .......... .......... .......... .......... .......... .......... .......... .......... .......... .......... .......-  116
  U5_99_5 -......... .......... .......... .......... .......... .......... .......... .......... .......... .......... .......... .......-  116
 U5_100_5 -......... .......... .......... .......... .......... .......... .......... .......... .......... .......... .......... .......-  116
 U5_101_5 -......... .......... .......... .......... .......... .......... .......... .......... .......... .......... .......... .......-  116
 U5_102_5 -......... .......... .......... .......... .......... .......... .......... .......... .......... .......... .......... .......-  116
 U5_103_5 -......... .......... .......... .......... .......... .......... .......... .......... .......... .......... .......... .......-  116
 U5_104_5 -......... .......... .......... .......... .......... .......... .......... .......... .......... .......... .......... .......-  116
 U5_105_5 -......... .......... .......... .......... .......... .......... .......... .......... .......... .......... .......... .......-  116
 U5_106_5 -......... .......... .......... .......... .......... .......... .......... .......... .......... .......... .......... .......-  116
 U5_107_5 -......... .......... .......... .......... .......... .......... .......... .......... .......... .......... .......... .......-  116
 U5_108_5 -......... .......... .......... .......... .......... .......... .......... .......... .......... .......... .......... .......-  116
 U5_109_5 -......... .......... .......... .......... .......... .......... .......... .......... .......... .......... .......... .......-  116
 U5_110_5 -......... .......... .......... .......... .......... .......... .......... .......... .......... .......... .......... .......-  116
 U5_111_5 -......... .......... .......... .......... .......... .......... .......... .......... .......... .......... .......... .......-  116
 U5_112_5 -......... .......... .......... .......... .......... .......... .......... .......... .......... .......... .......... .......-  116
 U5_113_5 -......... .......... .......... .......... .......... .......... .......... .......... .......... .......... .......... .......-  116
 U5_114_5 -......... .......... .......... .......... .......... .......... .......... .......... .......... .......... .......... .......-  116
 U5_115_5 -......... .......... .......... .......... .......... .......... .......... .......... .......... .......... .......... .......-  116
 U5_116_5 -......... .......... .......... .......... .......... .......... .......... .......... .......... .......... .......... .......-  116
 U5_117_5 -......... .......... .......... .......... .......... .......... .......... .......... .......... .......... .......... .......-  116
 U5_118_5 -......... .......... .......... .......... .......... .......... .......... .......... .......... .......... .......... .......-  116
 U5_119_5 -......... .......... .......... .......... .......... .......... .......... .......... .......... .......... .......... .......-  116
 U5_120_5 -......... .......... .......... .......... .......... .......... .......... .......... .......... .......... .......... .......-  116
 U5_121_5 -......... .......... .......... .......... .......... .......... .......... .......... .......... .......... .......... .......-  116
 U5_122_5 -......... .......... .......... .......... .......... .......... .......... .......... .......... .......... .......... .......-  116
 U5_123_5 -......... .......... .......... .......... .......... .......... .......... .......... .......... .......... .......... .......-  116
 U5_124_5 -......... .......... .......... .......... .......... .......... .......... .......... .......... .......... .......... .......-  116
 U5_125_5 -......... .......... .......... .......... .......... .......... .......... .......... .......... .......... .......... .......-  116
   U5_4_5 -......A.. .......... .......... .......... .......... .......... .......... .......... .......... .......... .......... .......-  116
  U5_82_5 -......... .......... .......... .......... .......... .......... .........T .......... .......... .......... .......... .......-  116
 U5_130_5 -......... .......... .......... .......... .......... .......... ........G. .......... .......... .......... .......... .......-  116
  U5_10_5 -......... .......A.. .......... .......... .......... .......... .......... .......... .......... .......... .......... .......-  116
   U5_9_5 -......... .......... .......... .......... .......... .T........ .......... .......... .......... .......... .......... .......-  116
  U5_79_5 -......... .......... .......... .......... .......... .T........ .......... .......... .......... .......... .......... .......-  116
   U5_1_5 -......... .......... .......... .......... .......... .......... .......... ........A. .......... .......... .......... .......-  116
   U5_2_5 -......... .......... .....T.... .......... .......... .......... .......... ........A. .......... .......... .......... .......-  116
   U5_3_5 -......... .......... .A........ .......... .......... .......... .......... .......... .A........ .......... .......... .......-  116
  U5_76_5 -.....T... .....G.... .......... .......... .......... .......... .......... .......... .......... .......... .......... .......-  116
 U5_132_5 -T.....A.. .......... .......T.. .........T T......... .......... .......... .......... .......... .......... .......... .......-  116
U5_136_20 -ACA...AT. .......... GAT-..T.T. .......... .......... .......... .......... ACAATGAGTA .AACC.A... ...T.A...C ....CTATC. GT.G.GCT 116
U5_139_20 -ACA...A.. .......... GAT-..T.T. .......... .......... .......... .......... .CAATGAGTA .AATC.A... ...T.A...C ....CTATC. GT.G.GCT 116
U5_134_20 -ACA...AT. ...T..C... GAT-..T.T. .......... .......... .......... .....A.... AC.ATGAGTA .AAAC.A... ...T.A...C ....CGATA. GC.G.GCT 116
U5_135_20 -ACA...AT. ....A.C... GAT-..T.T. .......... .......... .......... .......... ACAATGAGTA ..AAC.A... ...T.A...C ....A.CA.. .G.G..C- 115
 U5_133_9 -ACA...AT. .......... TAT-..A.T. .G........ .......... ........A. ..A....... ACC.TGAGTA ...AC.A... ...T.A..TC ...G.TCAAT GTAG.A.T 116
U5_138_20 -ACA...AT. .......... GAT-..T.T. .......... .......... .......... .......... ACAATGAGTA .AATC.A... ....ATGC.. TGCT...ACC GC.G.GCT 116
U5_137_20 AGCT..T..C ...T...... GAT-..T.T. .......... .......... .......... .......... ACAATGAGTA .AAAC.A... ....ATGC.C TGCT...ACC GC.G.GC- 116
```

```
                                     20                    40                    60                    80                    100
                                      |                     |                     |                     |                     |
U6consensus GTGCTCGCTA CGGTGGCACA TATACTAAAA TTGGATCGAT ACAGAGAAGA TTAGCATGGC CCCTGCGAAA GGATGACACG CAAATCCGTG AAGCGCTCCA TATTGCT 107
   U6_276_4 -GTG...... .......... .......... .......... .......... .........T .......... ........T. .......... .......... ....... 106
   U6_368_4 ......A... T...T..... .......... .......... .......... .......... ......A... .......... .......... .......... ....--- 104
     U6_2_3 .......... T......... .......... ...A...... G......... .......... .......... .......... .......... .......... ....--- 104
   U6_341_4 ..A....... .......... .......... .........A .......... .......... .......... .......... .......... .......... ....--- 104
   U6_343_4 ..A....... .......... .......... .........A .......... .......... .......... .......... .......... .......... ....--- 104
   U6_294_4 .......... .......... .......... .......... ........T. .......... .......... .......... .......... .......T.. ....... 107
   U6_302_4 .......... .......... .......... .......... ........T. .......... .......... .......... .......... .......T.. ....... 107
   U6_306_4 .......... .......... .......... .......... ........T. .......... .......... .......... .......... .......T.. ....... 107
   U6_310_4 .......... .......... .......... .......... ........T. .......... .......... .......... .......... .......T.. ....... 107
     U6_8_3 .......... .......... .......... .......... .....A.... .......... .......... .......... .......... .......... ....--- 104
   U6_281_4 .......... .......... .......... ...C...... .....A.... .......... .......... .......... .......... .......... ....... 107
    U6_85_4 .......... .......... .......... .......... .......... .......... .......... .G........ .......... .......... ....--- 104
   U6_101_4 .......... .......... .......... .......... .......... .......... .......... .G........ .......... .......... ....--- 104
   U6_121_4 .....T.... .......... .......... .......... .......... .......... .......... .G........ .......... .......... ....--- 104
   U6_172_4 .......... .......... .......... ...C...... .......... .......... .......... .......... .......... .......... ....T.. 107
   U6_503_4 .......... .......... .......... ...C...... .......... .......... .......... .......... .......... .......... ....... 107
   U6_507_4 .......... .......... .......... ...C...... .......... .......... .......... .......... .......... .......... ....... 107
    U6_87_4 .......... A......... .......... .......... .......... .......... .......... .......... .......... .......... ....--- 104
   U6_123_4 .......... T......... .......... .......... .......... .......... .......... .......... .......... .......... ....--- 104
   U6_261_4 .......... T......... .......... .......... .......... .......... .......... .......... .......... .......... ....... 107
   U6_285_4 .......... T.-....... .......... .......... .......... .......... .......... .......... .......... .......... ....... 106
   U6_342_4 .......... T......... .......... .......... .......... .......... .......... .......... .......... .......... ....--- 104
   U6_179_4 .......... .......... .......... .......... .......... ........A. .......... .......... .......... .......... ....--- 104
   U6_185_4 .......... .......... .......... .......... .......... ........A. .......... .......... .......... .......... ....--- 104
   U6_436_4 .......... .......... .......... .......... .......... ........T. .......... .......... .......... .......... ....... 107
    U6_25_4 .......... .......... .......... .......... .......... .......... .......... .........A .......... .......... ....... 107
   U6_245_4 .......... .......... .......... .......... .......... .......... .......... .........T .......... .......... ....... 107
   U6_146_4 .......... .......... .T........ .......... .......... .......... .......... .......... .......... .......... ....... 107
   U6_165_4 .......... .......... .T........ .......... .......... .......... .......... .......... .......... .......... ....... 107
    U6_98_4 .......... .......... .......... .......... .......... .......... .......... .......... .......... .......... ....--- 104
   U6_100_4 .......... .......... .......... .......... .......... .......... .......... .......... .......... .......... ....--- 104
   U6_103_4 .......... .......... .......... .......... .......... .......... .......... .......... .......... .......... ....--- 104
   U6_106_4 .......... .......... .......... .......... .......... .......... .......... .......... .......... .......... ....--- 104
   U6_107_4 .......... .......... .......... .......... .......... .......... .......... .......... .......... .......... ....--- 104
   U6_109_4 .......... .......... .......... .......... .......... .......... .......... .......... .......... .......... ....--- 104
   U6_110_4 .......... .......... .......... .......... .......... .......... .......... .......... .......... .......... ....--- 104
   U6_111_4 .......... .......... .......... .......... .......... .......... .......... .......... .......... .......... ....--- 104
   U6_112_4 .......... .......... .......... .......... .......... .......... .......... .......... .......... .......... ....--- 104
   U6_114_4 .......... .......... .......... .......... .......... .......... .......... .......... .......... .......... ....--- 104
   U6_116_4 .......... .......... .......... .......... .......... .......... .......... .......... .......... .......... ....--- 104
   U6_117_4 .......... .......... .......... .......... .......... .......... .......... .......... .......... .......... ....--- 104
   U6_119_4 .......... .......... .......... .......... .......... .......... .......... .......... .......... .......... ....--- 104
   U6_120_4 .......... .......... .......... .......... .......... .......... .......... .......... .......... .......... ....--- 104
   U6_122_4 .......... .......... .......... .......... .......... .......... .......... .......... .......... .......... ....--- 104
   U6_124_4 .......... .......... .......... .......... .......... .......... .......... .......... .......... .......... ....--- 104
   U6_126_4 .......... .......... .......... .......... .......... .......... .......... .......... .......... .......... ....--- 104
   U6_127_4 .......... .......... .......... .......... .......... .......... .......... .......... .......... .......... ....--- 104
   U6_129_4 .......... .......... .......... .......... .......... .......... .......... .......... .......... .......... ....--- 104
   U6_131_4 .......... .......... .......... .......... .......... .......... .......... .......... .......... .......... ....--- 104
   U6_133_4 .......... .......... .......... .......... .......... .......... .......... .......... .......... .......... ....... 107
   U6_135_4 .......... .......... .......... .......... .......... .......... .......... .......... .......... .......... ....... 107
   U6_139_4 .......... .......... .......... .......... .......... .......... .......... .......... .......... .......... ....... 107
   U6_141_4 .......... .......... .......... .......... .......... .......... .......... .......... .......... .......... ....... 107
   U6_142_4 .......... .......... .......... .......... .......... .......... .......... .......... .......... .......... ....... 107
   U6_147_4 .......... .......... .......... .......... .......... .......... .......... .......... .......... .......... ....... 107
   U6_155_4 .......... .......... .......... .......... .......... .......... .......... .......... .......... .......... ....... 107
   U6_156_4 .......... .......... .......... .......... .......... .......... .......... .......... .......... .......... ....... 107
   U6_158_4 .......... .......... .......... .......... .......... .......... .......... .......... .......... .......... ....... 107
   U6_159_4 .......... .......... .......... .......... .......... .......... .......... .......... .......... .......... ....... 107
   U6_160_4 .......... .......... .......... .......... .......... .......... .......... .......... .......... .......... ....... 107
   U6_161_4 .......... .......... .......... .......... .......... .......... .......... .......... .......... .......... ....... 107
   U6_163_4 .......... .......... .......... .......... .......... .......... .......... .......... .......... .......... ....... 107
   U6_164_4 .......... .......... .......... .......... .......... .......... .......... .......... .......... .......... ....... 107
   U6_173_4 .......... .......... .......... .......... .......... .......... .......... .......... .......... .......... ....... 107
   U6_177_4 .......... .......... .......... .......... .......... .......... .......... .......... .......... .......... ....--- 104
   U6_181_4 .......... .......... .......... .......... .......... .......... .......... .......... .......... .......... ....--- 104
   U6_182_4 .......... .......... .......... .......... .......... .......... .......... .......... .......... .......... ....--- 104
   U6_183_4 .......... .......... .......... .......... .......... .......... .......... .......... .......... .......... ....--- 104
   U6_186_4 .......... .......... .......... .......... .......... .......... .......... .......... .......... .......... ....--- 104
   U6_187_4 .......... .......... .......... .......... .......... .......... .......... .......... .......... .......... ....--- 104
   U6_188_4 .......... ..-....... .......... .......... .......... .......... .......... .......... .......... .......... ....--- 103
   U6_190_4 .......... .......... .......... .......... .......... .......... .......... .......... .......... .......... ....--- 104
   U6_191_4 .......... ..-....... .......... .......... .......... .......... .......... .......... .......... .......... ....--- 103
   U6_193_4 .......... .......... .......... .......... .......... .......... .......... .......... .......... .......... ....--- 104
   U6_195_4 .......... .......... .......... .......... .......... .......... .......... .......... .......... .......... ....--- 104
   U6_197_4 .......... .......... .......... .......... .......... .......... .......... .......... .......... .......... ....--- 104
   U6_199_4 .......... .......... .......... .......... .......... .......... .......... .......... .......... .......... ....--- 104
   U6_202_4 .......... .......... .......... .......... .......... .......... .......... .......... .......... .......... ....--- 104
   U6_203_4 .......... .......... .......... .......... .......... .......... .......... .......... .......... .......... ....--- 104
   U6_204_4 .......... .......... .......... .......... .......... .......... .......... .......... .......... .......... ....--- 104
   U6_205_4 .......... .......... .......... .......... .......... .......... .......... .......... .......... .......... ....--- 104
   U6_206_4 .......... .......... .......... .......... .......... .......... .......... .......... .......... .......... ....--- 104
   U6_209_4 .......... .......... .......... .......... .......... .......... .......... .......... .......... .......... ....--- 104
   U6_212_4 .......... .......... .......... .......... .......... .......... .......... .......... .......... .......... ....--- 104
   U6_214_4 .......... .......... .......... .......... .......... .......... .......... .......... .......... .......... ....--- 104
   U6_223_4 .......... .......... .......... .......... .......... .......... .......... .......... .......... .......... ....... 107
   U6_224_4 .......... .......... .......... .......... .......... .......... .......... .......... .......... .......... ....... 107
   U6_225_4 .......... .......... .......... .......... .......... .......... .......... .......... .......... .......... ....... 107
   U6_226_4 .......... .......... .......... .......... .......... .......... .......... .......... .......... .......... ....... 107
   U6_227_4 .......... .......... .......... .......... .......... .......... .......... .......... .......... .......... ....... 107
```

```
U6_229_4 .......... .......... .......... .......... .......... .......... .......... .......... .......... .......... ....... 107
U6_230_4 .......... .......... .......... .......... .......... .......... .......... .......... .......... .......... ....... 107
U6_231_4 .......... .......... .......... .......... .......... .......... .......... .......... .......... .......... ....... 107
U6_232_4 .......... .......... .......... .......... .......... .......... .......... .......... .......... .......... ....... 107
U6_233_4 .......... .......... .......... .......... .......... .......... .......... .......... .......... .......... ....... 107
U6_234_4 .......... .......... .......... .......... .......... .......... .......... .......... .......... .......... ....... 107
U6_235_4 .......... .......... .......... .......... .......... .......... .......... .......... .......... .......... ....... 107
U6_236_4 .......... .......... .......... .......... .......... .......... .......... .......... .......... .......... ....... 107
U6_237_4 .......... .......... .......... .......... .......... .......... .......... .......... .......... .......... ....... 107
U6_238_4 .......... .......... .......... .......... .......... .......... .......... .......... .......... .......... ....... 107
U6_240_4 .......... .......... .......... .......... .......... .......... .......... .......... .......... .......... ....... 107
U6_241_4 .......... .......... .......... .......... .......... .......... .......... .......... .......... .......... ....... 107
U6_242_4 .......... .......... .......... .......... .......... .......... .......... .......... .......... .......... ....... 107
U6_243_4 .......... .......... .......... .......... .......... .......... .......... .......... .......... .......... ....... 107
U6_244_4 .......... .......... .......... .......... .......... .......... .......... .......... .......... .......... ....... 107
U6_246_4 .......... .......... .......... .......... .......... .......... .......... .......... .......... .......... ....... 107
U6_247_4 .......... .......... .......... .......... .......... .......... .......... .......... .......... .......... ....... 107
U6_248_4 .......... .......... .......... .......... .......... .......... .......... .......... .......... .......... ....... 107
U6_250_4 .......... .......... .......... .......... .......... .......... .......... .......... .......... .......... ....... 107
U6_251_4 .......... .......... .......... .......... .......... .......... .......... .......... .......... .......... ....... 107
U6_253_4 .......... .......... .......... .......... .......... .......... .......... .......... .......... .......... ....... 107
U6_254_4 .......... .......... .......... .......... .......... .......... .......... .......... .......... .......... ....... 107
U6_262_4 .......... .......... .......... .......... .......... .......... .......... .......... .......... .......... ....... 107
U6_263_4 .......... .......... .......... .......... .......... .......... .......... .......... .......... .......... ....... 107
U6_266_4 .......... .......... .......... .......... .......... .......... .......... .......... .......... .......... ....... 107
U6_268_4 .......... .......... .......... .......... .......... .......... .......... .......... .......... .......... ....... 107
U6_277_4 .......... .......... .......... .......... .......... .......... .......... .......... .......... .......... ....... 107
U6_278_4 .......... .......... .......... .......... .......... .......... .......... .......... .......... .......... ....... 107
U6_284_4 .......... .......... .......... .......... .......... .......... .......... .......... .......... .......... ....... 107
U6_292_4 .......... .......... .......... .......... .......... .......... .......... .......... .......... .......... ....... 107
U6_295_4 .......... .......... .......... .......... .......... .......... .......... .......... .......... .......... ....... 107
U6_298_4 .......... .......... .......... .......... .......... .......... .......... .......... .......... .......... ....... 107
U6_299_4 .......... .......... .......... .......... .......... .......... .......... .......... .......... .......... ....... 107
U6_303_4 .......... .......... .......... .......... .......... .......... .......... .......... .......... .......... ....... 107
U6_307_4 .......... .......... .......... .......... .......... .......... .......... .......... .......... .......... ....... 107
U6_311_4 .......... .......... .......... .......... .......... .......... .......... .......... .......... .......... ....... 107
U6_317_4 .......... .......... .......... .......... .......... .......... .......... .......... .......... .......... ....--- 104
U6_322_4 .......... .......... .......... .......... .......... .......... .......... .......... .......... .......... ....--- 104
U6_323_4 .......... .......... .......... .......... .......... .......... .......... .......... .......... .......... ....--- 104
U6_336_4 .......... .......... .......... .......... .......... .......... .......... .......... .......... .......... ....--- 104
U6_337_4 .......... .......... .......... .......... .......... .......... .......... .......... .......... .......... ....--- 104
U6_338_4 .......... .......... .......... .......... .......... .......... .......... .......... .......... .......... ....--- 104
U6_340_4 .......... .......... .......... .......... .......... .......... .......... .......... .......... .......... ....--- 104
U6_364_4 .......... .......... .......... .......... .......... .......... .......... .......... .......... .......... ....--- 104
U6_366_4 .......... .......... .......... .......... .......... .......... .......... .......... .......... .......... ....--- 104
U6_375_4 .......... .......... .......... .......... .......... .......... .......... .......... .......... .......... ....... 107
U6_378_4 .......... .......... .......... .......... .......... .......... .......... .......... .......... .......... ....... 107
U6_382_4 .......... .......... .......... .......... .......... .......... .......... .......... .......... .......... ....... 107
U6_384_4 .......... .......... .......... .......... .......... .......... .......... .......... .......... .......... ....... 107
U6_385_4 .......... .......... .......... .......... .......... .......... .......... .......... .......... .......... ....... 107
U6_386_4 .......... .......... .......... .......... .......... .......... .......... .......... .......... .......... ....... 107
U6_389_4 .......... .......... .......... .......... .......... .......... .......... .......... .......... .......... ....... 107
U6_391_4 .......... .......... .......... .......... .......... .......... .......... .......... .......... .......... ....... 107
U6_392_4 .......... .......... .......... .......... .......... .......... .......... .......... .......... .......... ....... 107
U6_393_4 .......... .......... .......... .......... .......... .......... .......... .......... .......... .......... ..----- 102
U6_395_4 .......... .......... .......... .......... .......... .......... .......... .......... .......... .......... ....... 107
U6_396_4 .......... .......... .......... .......... .......... .......... .......... .......... .......... .......... ....... 107
U6_397_4 .......... .......... .......... .......... .......... .......... .......... .......... .......... .......... ..----- 102
U6_398_4 .......... .......... .......... .......... .......... .......... .......... .......... .......... .......... ....... 107
U6_399_4 .......... .......... .......... .......... .......... .......... .......... .......... .......... .......... ....... 107
U6_400_4 .......... .......... .......... .......... .......... .......... .......... .......... .......... .......... ....... 107
U6_401_4 .......... .......... .......... .......... .......... .......... .......... .......... .......... .......... ....... 107
U6_403_4 .......... .......... .......... .......... .......... .......... .......... .......... .......... .......... ....... 107
U6_404_4 .......... .......... .......... .......... .......... .......... .......... .......... .......... .......... ....... 107
U6_405_4 .......... .......... .......... .......... .......... .......... .......... .......... .......... .......... ....... 107
U6_408_4 .......... .......... .......... .......... .......... .......... .......... .......... .......... .......... ....... 107
U6_409_4 .......... .......... .......... .......... .......... .......... .......... .......... .......... .......... ..----- 102
 U6_28_4 .......... .......... .......... .......... .......... .......... .......... .......... .......... .......... ....... 107
 U6_29_4 .......... .......... .......... .......... .......... .......... .......... .......... .......... .......... ....... 107
 U6_30_4 .......... .......... .......... .......... .......... .......... .......... .......... .......... .......... ....... 107
 U6_33_4 .......... .......... .......... .......... .......... .......... .......... .......... .......... .......... ....... 107
 U6_35_4 .......... .......... .......... .......... .......... .......... .......... .......... .......... .......... ..----- 102
 U6_39_4 .......... .......... .......... .......... .......... .......... .......... .......... .......... .......... ....... 107
 U6_40_4 .......... .......... .......... .......... .......... .......... .......... .......... .......... .......... ....... 107
 U6_43_4 .......... .......... .......... .......... .......... .......... .......... .......... .......... .......... ....... 107
 U6_45_4 .......... .......... .......... .......... .......... .......... .......... .......... .......... .......... ....... 107
 U6_46_4 .......... .......... .......... .......... .......... .......... .......... .......... .......... .......... ....... 107
 U6_47_4 .......... .......... .......... .......... .......... .......... .......... .......... .......... .......... ....... 107
 U6_50_4 .......... .......... .......... .......... .......... .......... .......... .......... .......... .......... ....... 107
 U6_52_4 .......... .......... .......... .......... .......... .......... .......... .......... .......... .......... ..----- 102
 U6_55_4 .......... .......... .......... .......... .......... .......... .......... .......... .......... .......... ....... 107
 U6_56_4 .......... .......... .......... .......... .......... .......... .......... .......... .......... .......... ..----- 102
 U6_58_4 .......... .......... .......... .......... .......... .......... .......... .......... .......... .......... ....... 107
 U6_59_4 .......... .......... .......... .......... .......... .......... .......... .......... .......... .......... ....... 107
 U6_62_4 .......... .......... .......... .......... .......... .......... .......... .......... .......... .......... ....... 107
 U6_64_4 .......... .......... .......... .......... .......... .......... .......... .......... .......... .......... ....... 107
 U6_65_4 .......... .......... .......... .......... .......... .......... .......... .......... .......... .......... ....--- 104
 U6_67_4 .......... .......... .......... .......... .......... .......... .......... .......... .......... .......... ....... 107
 U6_68_4 .......... .......... .......... .......... .......... .......... .......... .......... .......... .......... ....... 107
 U6_70_4 .......... .......... .......... .......... .......... .......... .......... .......... .......... .......... ....... 107
 U6_71_4 .......... .......... .......... .......... .......... .......... .......... .......... .......... .......... ....... 107
 U6_73_4 .......... .......... .......... .......... .......... .......... .......... .......... .......... .......... ....--- 104
 U6_79_4 .......... .......... .......... .......... .......... .......... .......... .......... .......... .......... ....--- 104
```

```
 U6_81_4  .......... .......... .......... .......... .......... .......... .......... .......... .......... .......... ....---  104
 U6_84_4  .......... .......... .......... .......... .......... .......... .......... .......... .......... .......... ....---  104
 U6_88_4  .......... .......... .......... .......... .......... .......... .......... .......... .......... .......... ....---  104
 U6_90_4  .......... .......... .......... .......... .......... .......... .......... .......... .......... .......... ....---  104
 U6_93_4  .......... .......... .......... .......... .......... .......... .......... .......... .......... .......... ....---  104
 U6_96_4  .......... .......... .......... .......... .......... .......... .......... .......... .......... .......... ....---  104
  U6_3_3  .......... .......... .......... .......... .......... .......... .......... .......... .......... .......... ....---  104
  U6_4_3  .......... .......... .......... .......... .......... .......... .......... .......... .......... .......... ....---  104
  U6_5_3  .......... .......... .......... .......... .......... .......... .......... .......... .......... .......... ....---  104
  U6_7_3  .......... .......... .......... .......... .......... .......... .......... .......... .......... .......... ....---  104
 U6_20_4  .......... .......... .......... .......... .......... .......... .......... .......... .......... .......... .......  107
 U6_22_4  .......... .......... .......... .......... .......... .......... .......... .......... .......... .......... .......  107
 U6_24_4  .......... .......... .......... .......... .......... .......... .......... .......... .......... .......... .......  107
 U6_27_4  .......... .......... .......... .......... .......... .......... .......... .......... .......... .......... ..-----  102
U6_508_4  .......... .......... .......... .......... .......... .......... .......... .......... .......... .......... .......  107
U6_512_4  .......... .......... .......... .......... .......... .......... .......... .......... .......... .......... ....---  104
U6_513_4  .......... .......... .......... .......... .......... .......... .......... .......... .......... .......... ....---  104
U6_488_4  .......... .......... .......... .......... .......... .......... .......... .......... .......... .......... .......  107
U6_492_4  .......... .......... .......... .......... .......... .......... .......... .......... .......... .......... .......  107
U6_493_4  .......... .......... .......... .......... .......... .......... .......... .......... .......... .......... ...----  103
U6_494_4  .......... .......... .......... .......... .......... .......... .......... .......... .......... .......... .......  107
U6_497_4  .......... .......... .......... .......... .......... .......... .......... .......... .......... .......... .......  107
U6_500_4  .......... .......... .......... .......... .......... .......... .......... .......... .......... .......... .......  107
U6_501_4  .......... .......... .......... .......... .......... .......... .......... .......... .......... .......... .......  107
U6_504_4  .......... .......... .......... .......... .......... .......... .......... .......... .......... .......... .......  107
U6_411_4  .......... .......... .......... .......... .......... .......... .......... .......... .......... .......... .......  107
U6_412_4  .......... .......... .......... .......... .......... .......... .......... .......... .......... .......... .......  107
U6_415_4  .......... .......... .......... .......... .......... .......... .......... .......... .......... .......... .......  107
U6_417_4  .......... .......... .......... .......... .......... .......... .......... .......... .......... .......... .......  107
U6_418_4  .......... .......... .......... .......... .......... .......... .......... .......... .......... .......... .......  107
U6_419_4  .......... .......... .......... .......... .......... .......... .......... .......... .......... .......... ..-----  102
U6_422_4  .......... .......... .......... .......... .......... .......... .......... .......... .......... .......... .......  107
U6_428_4  .......... .......... .......... .......... .......... .......... .......... .......... .......... .......... .......  107
U6_431_4  .......... .......... .......... .......... .......... .......... .......... .......... .......... .......... .......  107
U6_432_4  .......... .......... .......... .......... .......... .......... .......... .......... .......... .......... .......  107
U6_437_4  .......... .......... .......... .......... .......... .......... .......... .......... .......... .......... .......  107
U6_440_4  .......... .......... .......... .......... .......... .......... .......... .......... .......... .......... .......  107
U6_441_4  .......... .......... .......... .......... .......... .......... .......... .......... .......... .......... .......  107
U6_443_4  .......... .......... .......... .......... .......... .......... .......... .......... .......... .......... .......  107
U6_444_4  .......... .......... .......... .......... .......... .......... .......... .......... .......... .......... .......  107
U6_446_4  .......... .......... .......... .......... .......... .......... .......... .......... .......... .......... .......  107
U6_449_4  .......... .......... .......... .......... .......... .......... .......... .......... .......... .......... ....---  104
U6_450_4  .......... .......... .......... .......... .......... .......... .......... .......... .......... .......... ....---  104
U6_453_4  .......... .......... .......... .......... .......... .......... .......... .......... .......... .......... ....---  104
U6_456_4  .......... .......... .......... .......... .......... .......... .......... .......... .......... .......... ....---  104
U6_457_4  .......... .......... .......... .......... .......... .......... .......... .......... .......... .......... ....---  104
U6_464_4  .......... .......... .......... .......... .......... .......... .......... .......... .......... .......... ....---  104
U6_465_4  .......... .......... .......... .......... .......... .......... .......... .......... .......... .......... ....---  104
U6_467_4  .......... .......... .......... .......... .......... .......... .......... .......... .......... .......... ....---  104
U6_472_4  .......... .......... .......... .......... .......... .......... .......... .......... .......... .......... ....---  104
U6_473_4  .......... .......... .......... .......... .......... .......... .......... .......... .......... .......... ....---  104
U6_474_4  .......... .......... .......... .......... .......... .......... .......... .......... .......... .......... ....---  104
U6_475_4  .......... .......... .......... .......... .......... .......... .......... .......... .......... .......... ....---  104
U6_480_4  .......... .......... .......... .......... .......... .......... .......... .......... .......... .......... ....---  104
U6_482_4  .......... .......... .......... .......... .......... .......... .......... .......... .......... .......... ....---  104
U6_486_4  .......... .......... .......... .......... .......... .......... .......... .......... .......... .......... .......  107
U6_487_4  .......... .......... .......... .......... .......... .......... .......... .......... .......... .......... .......  107
U6_283_4  .......... .......... .......... .......... .......... .......... .......... .......... .......... .......... ....T..  107
U6_286_4  .......... .......... .......... .......... .......... .......... .......... .......... .......... .......... ....T..  107
U6_288_4  .......... .......... .......... .......... .......... .......... .......... .......... .......... .......... ....T..  107
U6_315_4  .......... .......... .......... .......... .......... .......... .......... .......... .......... .......... ....---  104
U6_325_4  .......... .......... .......... .......... .......... .......... .......... .......... .......... .......... ....---  104
U6_327_4  .......... .......... .......... .......... .......... .......... .......... .......... .......... .......... ....---  104
U6_331_4  .......... .......... .......... .......... .......... .......... .......... .......... .......... .......... ....---  104
U6_334_4  .......... .......... .......... .......... .......... .......... .......... .......... .......... .......... ....---  104
U6_344_4  .......... .......... .......... .......... .......... .......... .......... .......... .......... .......... ....---  104
U6_345_4  .......... .......... .......... .......... .......... .......... .......... .......... .......... .......... ....---  104
U6_346_4  .......... .......... .......... .......... .......... .......... .......... .......... .......... .......... ....---  104
U6_347_4  .......... .......... .......... .......... .......... .......... .......... .......... .......... .......... ....---  104
U6_348_4  .......... .......... .......... .......... .......... .......... .......... .......... .......... .......... ....---  104
U6_353_4  .......... .......... .......... .......... .......... .......... .......... .......... .......... .......... ....---  104
U6_360_4  .......... .......... .......... .......... .......... .......... .......... .......... .......... .......... ....---  104
U6_362_4  .......... .......... .......... .......... .......... .......... .......... .......... .......... .......... ....---  104
U6_270_4  .......... .......... .......... .......... ........A. .......... .......... .......... .......... .......... .......  107
U6_489_4  .......... .......... .......... .......... .......--- .......... .......... .......... .......... .......... .......  104
U6_279_4  .......... ..A....... .......... .......... .......... .......... .......... .......... .......... .......... .......  107
U6_269_4  .......... .......... .......... .......... .......... .......... .......... .......... .......... G......... .......  107
U6_434_4  .......... .......... .......... ....T..... .......... .......... .......... .......... .......... .......... .......  107
U6_414_4  .......... .......... .......... .......... .......... .......... .......... .....T.... .......... .......... .......  107
U6_442_4  .......... .......... .......... .......... .......... .......... .......... .......... ..G....... .......... .......  107
U6_490_4  .......... .......... .......... .......... .......... .......... .......... ......T... .......... .......... .......  107
 U6_21_4  .......... .......... .......... .......... .......... .......... .......... .......... .......... .....A.... .......  107
U6_293_4  .......... .......... .......... .......A.. .......... .......... .......... .......... .......... .......... .......  107
U6_496_4  .......... .......... .......... .......... .......... .......... .......... .A........ .......... .......... .......  107
U6_296_4  .......... .......... .......... .......... .......... .......... ..T....... .......... .......... .......... .......  107
U6_300_4  .......... .......... .......... .......... .......... .......... ..T....... .......... .......... .......... .......  107
U6_312_4  .......... .......... .......... .......... .......... .......... ..T....... .......... .......... .......... .......  107
U6_304_4  .......... .......... .......... .......... .......... .......... ..T....... .......... .......... .......... .......  107
U6_308_4  .......... .......... .......... .......... .......... .......... ..T....... .......... .......... .......... .......  107
U6_259_4  .......... ......T... .......... .......... .......... .......... .......... .......... .......... .......... .......  107
U6_264_4  .......... ......T... .......... .......... .......... .......... .......... .......... .......... .......... .......  107
U6_297_4  .......... ......G... .......... .......... .......... .......... .......... .......... .......... .......... .......  107
U6_301_4  .......... ......G... .......... .......... .......... .......... .......... .......... .......... .......... .......  107
```

```
U6_313_4 .......... ......G... .......... .......... .......... .......... .......... .......... .......... .......... ....... 107
U6_305_4 .......... ......G... .......... .......... .......... .......... .......... .......... .......... .......... ....... 107
U6_309_4 .......... ......G... .......... .......... .......... .......... .......... .......... .......... .......... ....... 107
U6_213_4 .......... .......... .......... .......... .......... .A........ .......... .......... .......... .......... ....--- 104
U6_485_4 .......... .......... .......... .......... .......... .A........ .......... .......... .......... .......... ....... 107
 U6_31_4 .......... .......... .......... .......... .......... .......... .......... .......... .......A.. .......... ....... 107
 U6_48_4 .......... .......... .......... .......... .......... .......... .......... .......... .......A.. .......... ....... 107
U6_447_4 .......... .......... .......... .......... .......... .......... .......... .......... .......A.. .......... ....--- 104
U6_171_4 .......... .......... .......... .......... .......... .......... .......... .......... .......A.. .......... ....... 107
 U6_83_4 .......... .......... .......... .......... .......... .......... .......... .......... .......A.. .......... ....--- 104
 U6_95_4 .......... .......... .......... .......... .......... .......... .......... .......... .......A.. .......... ....--- 104
U6_329_4 .......... .......... .......... .......... .......... .......... .......... .......... .......A.. .......... ....--- 104
 U6_92_4 .......... .......... .......... .......... .......... .......... .....A.... .......... .......... .......... ....--- 104
U6_319_4 .......... .......... .......... .......... .......... .......... .....A.... .......... .......... .......... ....--- 104
U6_324_4 .......... .......... .......... .......... .......... .......... .....A.... .......... .......... .......... ....--- 104
U6_333_4 .......... .......... .......... .......... .......... .......... .....A.... .......... .......... .......... ....--- 104
U6_339_4 .......... .......... .......... .......... .......... .......... .....A.... .......... .......... .......... ....--- 104
U6_439_4 .......... .......... .......... .......... .......... .......... .....A.... .......... .......... .......... ....... 107
U6_365_4 .......... .......... .......... .......... .......... .......... .....T.... .......... .......... .......... ....--- 104
U6_377_4 .......... .......... .......... .......... .......... .......... .....T.... .......... .......... .......... ....... 107
U6_435_4 .......... .......... .......... .......... .......... .......... .....T.... .......... .......... .......... ....... 107
U6_200_4 .......... .......... .......... .......... .......... .......... ......A... .......... .......... .......... ....--- 104
U6_256_4 .......... .......... .......... .......... .......... .......... ......A... .......... .......... .......... ....... 107
U6_454_4 .......... .......... .......... .......... .......... .......... ......A... .......... .......... .......... ....--- 104
U6_511_4 .......... .......... .......... .......... .......... .......... ......A... .......... .......... .......... ....--- 104
U6_316_4 .......... .......... .......... .......... .......... .......... .......G.. .......... .......... .......... ....--- 104
U6_320_4 .......... .......... .......... .......... .......... .......... .......... .......... .......... .......A.. ....--- 104
U6_452_4 .......... .......... .......... .......... .......... .......... .......... .......... .....T.... .......... ....--- 104
 U6_77_4 .......... .......... .......... .......... .......... .......... .......... .......... .......... ........A. ....--- 104
U6_357_4 .......... .......... .......... .......... .......... .......... .......... .......... .......... .........T ....--- 104
U6_481_4 .......... .......... .......... .......... ...A...... .......... .......... .......... .......... .......... ....--- 104
U6_354_4 .......... .......... .......... .......... .......... .......... .......... ....C..... .......... .......... ....--- 104
U6_358_4 .......... .......... .......... .......... .......... .......... .......... ....C..... .......... .......... ....--- 104
U6_367_4 .......... .......... .......... .......... .......... .......... .......... ....C..... .......... .......... ....--- 104
U6_328_4 .......... .......... ....A..... .......... .......... .......... .......... .......... .......... .......... ....--- 104
U6_361_4 .......... .......... ....T..... .......... .......... .......... .......... .......... .......... .......... ....--- 104
  U6_6_3 .......... .......... .......... .......... .......... .......... .......... .......... .........T .......... ....--- 104
U6_178_4 .......... .......... .......... .......... .......... .......... .......... .......... .........A .......... ....--- 104
U6_184_4 .......... .......... .......... .......... .......... .......... .......... .......... .........A .......... ....--- 104
U6_468_4 -......... .......... .......... .......... ......G... .......... .......... .......... .......... .......... ....--- 103
 U6_15_4 .....T.... .......... .......... .......... .......... .......... .......... .......... .......... ....A..... ....... 107
U6_381_4 .....T.... .......... .......... .......... .......... .......... .......... .......... .......... ....A..... ....... 107
U6_314_4 .....T.... .......... .......... .......... .......... .......... .......... .......... ......A... .......... ....... 107
U6_148_4 .....T.... .......... .......... .......... .......... .......... .......... .......... .......... .......... ....... 107
U6_149_4 .....T.... .......... .......... .......... .......... .......... .......... .......... .......... .......... ....... 107
U6_150_4 .....T.... .......... .......... .......... .......... .......... .......... .......... .......... .......... ....... 107
U6_152_4 .....T.... .......... .......... .......... .......... .......... .......... .......... .......... .......... ....... 107
U6_153_4 .....T.... .......... .......... .......... .......... .......... .......... .......... .......... .......... ....... 107
U6_162_4 .....T.... .......... .......... .......... .......... .......... .......... .......... .......... .......... ....... 107
U6_239_4 .....T.... .......... .......... .......... .......... .......... .......... .......... .......... .......... ....... 107
U6_249_4 .....T.... .......... .......... .......... .......... .......... .......... .......... .......... .......... ....... 107
U6_280_4 .....T.... .......... .......... .......... .......... .......... .......... .......... .......... .......... ....... 107
U6_383_4 .....T.... .......... .......... .......... .......... .......... .......... .......... .......... .......... ....... 107
U6_388_4 .....T.... .......... .......... .......... .......... .......... .......... .......... .......... .......... ....... 107
U6_390_4 .....T.... .......... .......... .......... .......... .......... .......... .......... .......... .......... ....... 107
U6_394_4 .....T.... .......... .......... .......... .......... .......... .......... .......... .......... .......... ....... 107
U6_402_4 .....T.... .......... .......... .......... .......... .......... .......... .......... .......... .......... ....... 107
U6_407_4 .....T.... .......... .......... .......... .......... .......... .......... .......... .......... .......... ....... 107
U6_413_4 .....T.... .......... .......... .......... .......... .......... .......... .......... .......... .......... ....... 107
U6_420_4 .....T.... .......... .......... .......... .......... .......... .......... .......... .......... .......... ....... 107
U6_421_4 .....T.... .......... .......... .......... .......... .......... .......... .......... .......... .......... ....... 107
U6_423_4 .....T.... .......... .......... .......... .......... .......... .......... .......... .......... .......... ....... 107
U6_424_4 .....T.... .......... .......... .......... .......... .......... .......... .......... .......... .......... ....... 107
U6_429_4 .....T.... .......... .......... .......... .......... .......... .......... .......... .......... .......... ....... 107
U6_433_4 .....T.... .......... .......... .......... .......... .......... .......... .......... .......... .......... ....... 107
U6_502_4 .....T.... .......... .......... .......... .......... .......... .......... .......... .......... .......... ....... 107
 U6_23_4 .....T.... .......... .......... .......... .......... .......... .......... .......... .......... .......... ..----- 102
 U6_74_4 --...T.... .......... .......... .......... .......... .......... .......... .......... .......... .......... ....--- 102
 U6_76_4 .....T.... .......... .......... .......... .......... .......... .......... .......... .......... .......... ....--- 104
 U6_78_4 --...T.... .......... .......... .......... .......... .......... .......... .......... .......... .......... ....--- 102
 U6_80_4 .....T.... .......... .......... .......... .......... .......... .......... .......... .......... .......... ....--- 104
 U6_82_4 .....T.... .......... .......... .......... .......... .......... .......... .......... .......... .......... ....--- 104
 U6_86_4 .....T.... .......... .......... .......... .......... .......... .......... .......... .......... .......... ....--- 104
 U6_91_4 .....T.... .......... .......... .......... .......... .......... .......... .......... .......... .......... ....--- 104
 U6_94_4 .....T.... .......... .......... .......... .......... .......... .......... .......... .......... .......... ....--- 104
 U6_38_4 .....T.... .......... .......... .......... .......... .......... .......... .......... .......... .......... ....... 107
 U6_44_4 .....T.... .......... .......... .......... .......... .......... .......... .......... .......... .......... ....... 107
 U6_49_4 .....T.... .......... .......... .......... .......... .......... .......... .......... .......... .......... ....... 107
 U6_53_4 .....T.... .......... .......... .......... .......... .......... .......... .......... .......... .......... ....... 107
 U6_54_4 .....T.... .......... .......... .......... .......... .......... .......... .......... .......... .......... ....... 107
 U6_57_4 .....T.... .......... .......... .......... .......... .......... .......... .......... .......... .......... ....... 107
 U6_63_4 .....T.... .......... .......... .......... .......... .......... .......... .......... .......... .......... ....... 107
 U6_66_4 .....T.... .......... .......... .......... .......... .......... .......... .......... .......... .......... ....... 107
 U6_69_4 .....T.... .......... .......... .......... .......... .......... .......... .......... .......... .......... ....... 107
U6_132_4 .....T.... .......... .......... .......... .......... .......... .......... .......... .......... .......... ....... 107
U6_134_4 .....T.... .......... .......... .......... .......... .......... .......... .......... .......... .......... ....... 107
U6_138_4 .....T.... .......... .......... .......... .......... .......... .......... .......... .......... .......... ....... 107
U6_140_4 .....T.... .......... .......... .......... .......... .......... .......... .......... .......... .......... ....... 107
U6_143_4 .....T.... .......... .......... .......... .......... .......... .......... .......... .......... .......... ....... 107
U6_144_4 .....T.... .......... .......... .......... .......... .......... .......... .......... .......... .......... ....... 107
U6_145_4 .....T.... .......... .......... .......... .......... .......... .......... .......... .......... .......... ....... 107
 U6_19_4 .....T.... .......... .......... .......... .......... .......... .......... .......... .......... .......... ....... 107
 U6_32_4 .....T.... .......... .......... .......... .......... .......... .......... .......... .......... .......... ....... 107
```

```
  U6_36_4  .....T.... .......... .......... .......... .......... .......... .......... .......... .......... .......... .......  107
  U6_37_4  .....T.... .......... .......... .......... .......... .......... .......... .......... .......... .......... .......  107
 U6_406_4  .....T.... .......... .......... .......... .......... .......... .......... .......... .......... .......... ..-----  102
 U6_410_4  .....T.... .......... .......... .......... .......... .......... .......... .......... .......... .......... ..-----  102
 U6_321_4  .....T.... .......... .......... .......... .......... .......... .......... .......... .......... .......... ....---  104
 U6_332_4  .....T.... .......... .......... .......... .......... .......... .......... .......... .......... .......... ....---  104
 U6_356_4  .....T.... .......... .......... .......... .......... .......... .......... .......... .......... .......... ....---  104
 U6_359_4  --...T.... .......... .......... .......... .......... .......... .......... .......... .......... .......... ....---  102
 U6_102_4  .....T.... .......... .......... .......... .......... .......... .......... .......... .......... .......... ....---  104
 U6_108_4  .....T.... .......... .......... .......... .......... .......... .......... .......... .......... .......... ....---  104
 U6_113_4  .....T.... .......... .......... .......... .......... .......... .......... .......... .......... .......... ....---  104
 U6_115_4  .....T.... .......... .......... .......... .......... .......... .......... .......... .......... .......... ....---  104
 U6_118_4  .....T.... .......... .......... .......... .......... .......... .......... .......... .......... .......... ....---  104
 U6_130_4  .....T.... .......... .......... .......... .......... .......... .......... .......... .......... .......... ....---  104
 U6_180_4  .....T.... .......... .......... .......... .......... .......... .......... .......... .......... .......... ....---  104
 U6_194_4  .....T.... .......... .......... .......... .......... .......... .......... .......... .......... .......... ....---  104
 U6_196_4  .....T.... .......... .......... .......... .......... .......... .......... .......... .......... .......... ....---  104
 U6_198_4  .....T.... .......... .......... .......... .......... .......... .......... .......... .......... .......... ....---  104
 U6_201_4  .....T.... .......... .......... .......... .......... .......... .......... .......... .......... .......... ....---  104
 U6_207_4  .....T.... .......... .......... .......... .......... .......... .......... .......... .......... .......... ....---  104
 U6_208_4  .....T.... .......... .......... .......... .......... .......... .......... .......... .......... .......... ....---  104
 U6_210_4  .....T.... .......... .......... .......... .......... .......... .......... .......... .......... .......... ....---  104
 U6_211_4  .....T.... .......... .......... .......... .......... .......... .......... .......... .......... .......... ....---  104
 U6_318_4  .....T.... .......... .......... .......... .......... .......... .......... .......... .......... .......... ....---  104
 U6_455_4  .....T.... .......... .....A.... .......... .......... .......... .......... .......... .......... .......... ....---  104
  U6_97_4  .....T.... .......... .......... .......... .......... .......... .....A.... .......... .......... .......... ....---  104
 U6_448_4  .....T.... .......... .......... .......... .......... .......... .......... .......... .......... .........C ....---  104
 U6_189_4  .....T.... .......... .......... .......... .......... .......... .....T.... .......... .......... .......... ....---  104
 U6_192_4  .....T.... .......... .......... .......... .......... .......... .....T.... .......... .......... .......... ....---  104
 U6_510_4  .....T.... .......... .......... .......... .......... .......... .....T.... .......... .......... .......... ....---  104
 U6_287_4  .....T.... .......... ..C....... .......... .......... .......... .....T.... .......... .......... .......... .......  107
 U6_374_4  .......... .......... .......... .......... .......... ........C. .......... .......... ......T... .......... .......  107
 U6_154_4  .......... .......... .......... .......... .......... .......... .......... .......... ......T... .......... .......  107
 U6_169_4  .......... .......... .......... .......... .......... .......... .......... .......... ......T... .......... .......  107
 U6_128_4  .......... .......... .......... .......... .......... .......... .......... .......... ......T... .......... ....---  104
 U6_416_4  .......... .......... .......... .......... .......... .......... .......... .......... ......T... .......... .......  107
 U6_438_4  .......... .......... .......... .......... .......... .......... .......... .......... ......T... .......... .......  107
 U6_491_4  .......... .......... .......... .......... .......... .......... .......... .......... ......T... .......... .......  107
 U6_495_4  .......... .......... .......... .......... .......... .......... .......... .......... ......T... .......... .......  107
 U6_289_4  .......... .......... .......... .......... .......... .......... .......... .......... ......T... .....T.... .......  107
 U6_376_4  .......... .A........ ..A....... .......... .......... .......... .......... .......... .......... .......... .......  107
  U6_89_4  .......... .A........ .......... .......... .......... .......... .......... .......... .......... .......... ....---  104
 U6_505_4  .......... .A........ .......... .......... .......... .......... .......... .......... .......... .......... .......  107
 U6_509_4  .......... .A........ .......... .......... .......... .......... .......... .......... .......... .......... .......  107
 U6_484_4  ....A..... .......... .......... .......... .......... .......... .......... .......... .......... .......... .......  107
 U6_157_4  ....AT.... .......... .......... .......... .......... .......... .......... .......... .......... .......... ....---  104
  U6_42_4  .....T.... .......... .......... .......... .......... .......... .......... A......... .......... .......... .......  107
  U6_61_4  .....T.... .......... .......... .......... .......... .......... .......... A......... .......... .......... .......  107
 U6_104_4  .....T.... .......... .......... .......... .......... .......... .......... A......... .......... .......... ....---  104
 U6_471_4  .......... .......... .......... .......... .......... .......... .......... A......... .......... .......... ....---  104
 U6_252_4  .......... .......... .......... .......... .......... .......... .......... ........T. .......... .......... .......  107
 U6_460_4  .......... .......... .......... .......... .......... .......... .......... ........T. .......... ....A..... ....---  104
 U6_425_4  ......C... .......... .......... .......... .......... .......... .......... .......... .......... .......... .......  107
 U6_255_4  .....TC... .......... .......... .......... .......... .......... .......... .......... .......... .......... .......  107
  U6_72_4  ......C... .......... .......... .......... .......... .......... .......... .......... .......... ....T..... ....---  104
  U6_75_4  ......C... .......... .......... .......... .......... .......... .......... .......... .......... ....T..... ....---  104
 U6_330_4  .......... .......... .......... .......... .......... .......... ....AA.... .......... .......... .......... ....---  104
 U6_335_4  .......... .......... .......... .......... .......... .......... ....AA.... .......... .......... .......... ....---  104
 U6_355_4  .......... .......... .......... .......... .......... .......... ....A..... .......... .......... .......... ....---  104
 U6_125_4  .......... .......... .......... ......T... .......... .......... .......... .......... .......... .......... ....---  104
  U6_16_4  .......... .......... .......... ......T... .......... .......... .......... .......... .......... ...T...... ..-----  102
  U6_18_4  .......... .......... .......... ......T... .......... .......... .......... .......... .......... ...T...... ..-----  102
 U6_387_4  .......... .......... .......... ......T... .......... .......... .......... .......... .......... ...T...... ..-----  102
  U6_26_4  .......... .......... .......... ......T... .......... .......... .......... .......... .......... ...T...... ..-----  102
  U6_41_4  .......... .......... .......... ......T... .......... .......... .......... .......... .......... ...T...... ..-----  102
  U6_60_4  .......... .......... .......... ......T... .......... .......... .......... .......... .......... ...T...... ..-----  102
 U6_105_4  .......... .......... .......... ......T... .......... .......... .......... .......... .......... ...T...... ..-----  102
 U6_168_4  .......... .......... .......... .......... .......... .......... .......... .......... .......... ..TT...... .......  107
 U6_257_4  .......... .......... .......... .......... .......... .......... .......... .......... .......... ...T...... .......  107
 U6_372_4  .......... .......... .......... .......... .......... .......... .......... .......... .......... ...T...... .......  107
 U6_430_4  .......... .......... .......... .......... .......... .......... .......... .......... .......... ...T...... .......  107
 U6_445_4  .......... .......... .......... .......... .......... .......... .......... .......... .......... ...T...... .......  107
 U6_506_4  .......... .......... .......... .......... .......... .......... .......... .......... .......... ...T...... .......  107
  U6_99_4  .......... .......... .......... .......... .......... .......... .......... .......... .......... ...T...... ....---  104
 U6_479_4  .......... .......... .......... .......... .......... .......... .....T.... .......... .......... ...T...... ....---  104
   U6_9_3  .....T.... .......... .......... ......A... .......... .......... .......... .......... .......... ...T...... ....---  104
 U6_514_4  .....T.... .......... .......... ......A... .......... .......... .......... .......... .......... ...T...... ....---  104
 U6_451_4  .....T.... .......... .......... .......... .......... .......... .......... .......... .......... ...T...... ....---  104
  U6_34_4  .....T.... .......... .......... ......T... .......... .......... .......... .......... .......... ...T...... ..-----  102
  U6_51_4  .....T.... .......... .......... ......T... .......... .......... .......... .......... .......... ...T...... ..-----  102
 U6_326_4  .......... ...C...... .......... .......... .......... .......... .....A.... .......... .......... .......... ....---  104
 U6_273_4  .......... .......... .......... ...T...... .......... .......... .......... .......... .......... .......... .G.....  107
 U6_267_4  .......... .......... .......... ..A....... .......... .......... .......... ..C....... .......... .......... .......  107
 U6_151_4  .......... .......... .......... .......... .......... .......... .......... .......... .......... ..T...C... .......  107
 U6_461_4  .......... T......... .......... .......... .......... .......... .......... .......... .......... .......... .T..---  104
 U6_466_4  .......... T......... .......... .......... .......... .......... .......... .......... .......... .......... .T..---  104
 U6_458_4  .......... ......T... .......... .......... .......... .........T .......... .......... .......... .......... ....---  104
 U6_459_4  .......... ......T... .......... .......... .......... .........T .......... .......... .......... .......... ....---  104
 U6_463_4  .......... ......T... .......... .......... .......... .........T .......... .......... .......... .......... ....---  104
  U6_10_3  .......... .......... .....A...G ..T....... .......... .......... .......... .......... .......... .......... ....---  104
 U6_216_4  .......... .......... .....A.... ..T....... .......... .......... .......... .......... .......... .......... ....---  104
 U6_350_4  .......... .......... .....A.... ..T....... .......... .......... .......... .......... .......... .......... ....---  104
 U6_363_4  .......... .......... .....A.... .......... .......... .......... .......... .......... .......... ....A..... ....---  104
```

```
  U6_170_4 .......... .A........ .......... .......... .......... .......... .....T.... .......... .......... .......... ....... 107
  U6_282_4 .......... .A........ .......... .......... .T........ .......... .....T.... .......... .......... .......... ....... 107
  U6_274_4 .....T.... ........T. .......... .......... .......... .......... .......... .......... .......... .......... .G..... 107
   U6_12_3 .......... .......... .......... .......... .......... .......... .......... .......... .......... ....A..... .T..--- 104
  U6_221_4 .......... .......... .......... .......... .......... .......... .......... .......... .......... ....A..... .T..--- 104
  U6_470_4 .......... .......... .......... .......... .......... .......... .......... .......... .......... ....A..... .T..--- 104
   U6_13_4 .......... .......... .......... .......... .......C.. .......... .......... .......... .......... ....A..... .T..--- 104
  U6_136_4 .......... .......... .......... .......... .......C.. .......... .......... .......... .......... ....A..... .T..--- 104
  U6_478_4 .......... .......... .......... .......... .......C.. .......... .......... .......... .......... ....A..... .T..--- 104
  U6_219_4 .......... .......... .......... .......... .......C.. .......... .......... .......... .......... ....A..... .T..--- 104
  U6_220_4 .......... .......... .......... .......... .......C.. .......... .......... .......... .......... ....A..... .T..--- 104
  U6_290_4 .......... .......... .......... .......... .......C.. .......... .......... .......... .......... ....A..... .T..--- 104
  U6_379_4 .......... .......... .......... .......... .......C.. .......... .......... .......... .......... ....A..... .T..--- 104
  U6_371_4 .......... T......... .......... .......... .......... .......... .....T.... .......... .......... ....A..... ....--- 104
  U6_175_4 .......... .......... .......... .......... .......... .......... .....T.T.. .......... .........A .......... ....... 107
  U6_260_4 .......... .......... .......... .......... .......... .......... .....T.... .......... ..G....... G......... ....... 107
  U6_426_4 .......... .......... .........G .......... ...A...... .......... .......... .......... .......... ....A..... ....... 107
  U6_370_4 .......... .......... .......... .......A.. .......... .......... ......A... .......... .......... ....A..... ....--- 104
  U6_265_4 .......... .......... .......... .......... .......... .......... .......... ........G. G......A.. .......... ....... 107
  U6_228_4 .....T.... .......... .G........ .......... ....G..... .......... .......... ........T. .......... .......... ....... 107
  U6_167_4 -......... .......... .......... ...T..T... ........A. .......... .......... .......... .......... ...T...... ....... 106
  U6_499_4 .......... .......... .......... .......... .......... .......... .....G.... .......... .....A.... .G........ .G..... 107
  U6_373_4 .......... .......... .......... ...A...... .......... .....TC... .......... .......... .........T .......... ....... 107
  U6_215_4 .....T.... .A........ .......... ......A... .......... ........T. .......... .......... .......... ...T...... ....--- 104
  U6_476_4 .....T.... .A........ .......... ......A... .......... ........T. .......... .......... .......... ...T...... ....--- 104
  U6_349_4 .....T.... .A........ .......... ......A... .......... .......... .......... .......... .......... ...T...... ....--- 104
  U6_427_4 -....T.... .A........ .......... ......A... .......... .......... ......A... .......... .......... .......... ....... 106
  U6_174_4 .....T.... .A........ .......... .......A.. ........A. ........-. .......... .......... .........A .......... ....... 106
    U6_1_3 .....T.... T.C....... .......... ....G..... .......... .......... .......... .......... ......T... ...T...... ....--- 104
  U6_176_4 .....T.... T......... .......... ....G..... .......... .......... .......... .......... .......... ...T...... ....--- 104
  U6_271_4 .....T.... T......... .......... ....G..... .......... .......... .....T.... .......... .......... ...T...... ....... 107
  U6_352_4 -....TA... T......... .......... ....G..... .......... .......... .......... .......... .......... ...T...... ....--- 103
  U6_351_4 .....T.... T......... .......... ....G..... .......... .......... .....A..T. .......... .......... ...T...... ....--- 104
  U6_166_4 .......... .......... .......... .......... .......... C.......A. .......... .......... .T........ ..TT...... ....... 107
  U6_498_4 .......... .......... .......... ......T... ........C. ....T..... ...G...... .......... ...G...... .......... ....... 107
  U6_275_4 ...T....CT A......... .......... .......... .......... .......... .......... .......... .......... .......... ....T.. 107
  U6_272_4 ...T....CT A......... .......... .......... ........C. .......... ...G...... .......... .......... .......... ....... 107
   U6_14_4 ......A... .......... .......... ......T... .......... .........T .......... .......... .........T .....A.... ....... 107
   U6_17_4 ......A... .......... .......... ......T... .......... .........T .......... .......... .........T .....A.... ....... 107
  U6_137_4 ......A... .......... .......... ......T... .......... .........T .......... .......... .........T .....A.... ....... 107
  U6_222_4 ......A... .......... .......... ......T... .......... .........T .......... .......... .........T .....A.... ....... 107
  U6_258_4 ......A... .......... .......... ......T... .......... .........T .......... .......... .........T .....A.... ....... 107
  U6_218_4 ......A... .......... .......... ......T... .......... .........T .......... .......... .........T .....A.... ....--- 104
  U6_291_4 ......A... .......... .......... ......T... .......... .........T .......... .......... ........AT .....A.... ....... 107
  U6_380_4 ......A... .......... .......... ......T... .......... .........T .......... .......... .G.......T .....A.... ....... 107
  U6_483_4 ......A... .......... .......... ......T... .......... .........T .......... .......... .....G...T .....A.... ....... 107
  U6_515_4 ......A... .......... .......... ......T... .......... .........T T......... .......... .........T .....A.... ....--- 104
   U6_11_3 ......A... .......... .......... ......T... .......... .........T .......... .......... .........A .....A.... ....--- 104
  U6_217_4 ......A... .......... .......... ......T... .......... .........T .......... .......... .........A .....A.... ....--- 104
  U6_369_4 ......A... .......... .......... ......T... .......... .........T .......... .......... .........A .....A.... ....--- 104
  U6_477_4 -.....A... .......... .......... ......T... .......... .........T .......... .......... ....C....T .....A.... ....--- 103
  U6_462_4 ...A..A... .......... .......... ......T... .......... .........T .....T.... .......... .........T .....A.... ....--- 104
  U6_469_4 ...A..A... .......... .......... ......T... .......... .........T .....T.... .......... .........T .....A.... ....--- 104
 U6_519_20 .......... ......-.-. C......... ..T....... .......... .......... .T..T.A... ....T...T. .......... .........G ....T-- 103
  U6_516_6 .....T...T ...CA..... .......... .....A.... .......... .......... .......C.. .......... .....T.... .....T.... .C..TTA 107
  U6_517_9 .....T...T ...CA..... .......... .....A.... .......... .......... .......C.. .......... .....T.... .....T.... .C..TTA 107
 U6_518_11 .....T...T ...CA..... .......... .....A.... .......... .......... .......C.. .......... .....T.... .....T.... .C..TTA 107
 U6_520_21 .....T...T ...CA..... .......... .....A.... .......... .......... .......C.. .......... .....T.... .....T...T C...TT- 106
```
