## Supplemental File SF2 for "An alternative spliceosome defined by distinct snRNAs in early zebrafish embryogenesis"

```
                                   20         40         60         80        100        120        140        160
U1Maternal ATACTTACCT GGCAGGGGAG ATACCATGAT CAAGAAGGTG GTTCACCCAG GGCGAGGCTT -GGCCATTGC ACTCCGGCCA CGCTGACCCC TGCGAATT-C CCCAAATGTG GGAATCTCGA CTGCATAATT TCTGGTAGTG GGGG-ACTGC GTTCGCGCTC TCCCCTG 164
U1_80_4    ...A...... .......... .......... .......... .......... .......... -......... .......... .......... ........-. .......... .......... .......... .........A ....-..... .....T.... ....... 164
U1_96_4    ...A...... .......... .......... .......... .......... .......... -......... .......... .......... ........-. .......... .......... .......... .........A ....-..... .....T.... ....... 164
U1_91_4    .......... .......... .......... .......... .......... .......... -......... .......... .......... ........-. .......... .......... .......... .........A ....-..... .....T.... ....... 164
U1_85_4    ...A...... .......... .......... .......... .......... .......... -......... .......... .......... ........-. .......... .......T.. .......... .........A ....-..... .......... ....... 164
U1_69_4    ........T. .......... .....G.... .......... .......... .......... -......... .......... .......... ........-. .......... .......... .......... .......... ....-..... .......... ....... 164
U1_70_4    ........T. .......... .....G.... .......... .......... .......... -......... .......... .......... ........-. .......... .......... .......... .......... ....-..... .......... ....... 164
U1_124_4   ........T. .......... .......... .....G.... .......... .......... -......... .......... .......... ........-. .......... .......... .......... .......... ....-..... .......... ....... 164
U1_103_4   .......... .......... .......... .......... .......... .......... -......... .......... .......... ........-. .......... .......... .......... ...A...... ....-..... .......... ....... 164
U1_119_4   .......... .......... .......... .......... .......... .......... -......... .......... .......... ........-. .......... .......... .......... ...A.A.... ....-..... .......... ....... 164
U1_137_4   .......... .......... .......... .......... .......... .......... -......... .......... .......... ........-. .......... .......... .G........ .......... ....-..... .......... ....... 164
U1_145_4   .......... .......... .......... .......... .......... .......... -......... G......... .......... ........-. .......... .......... .G........ .......... ....-..... .......... ....... 164
U1_5_4     .......... T......... .......... .......... .......... .......... -......... .......... .......... ........-. .......... .......... .......... .......... ....-..... .......... ....... 164
U1_22_4    .......... T......... .......... .......... .......... .......... -......... .......... .......... ........-. .......... .......... .......... .......... ....-..... .......... ....... 164
U1_29_4    .......... T......... .......... .......... .......... .......... -......... .......... .......... ........-. .......... .......... .......... .......... ....-..... .......... ....... 164
U1_9_4     .......... T......... .......... .......C.. .......... .......... -......... .......... .......... ........-. .......... .......... .......... .......... ....-..... .......... ....... 164
U1_27_4    .......... T......... .......... .......C.. .......... .......... -......... .......... .......... ........-. .......... .......... .......... .......... ....-..... .......... ....... 164
U1_35_4    .......... T......... .......... .......C.. .......... .......... -......... .......... .......... ........-. .......... .......... .......... .......... ....-..... .......... ....... 164
U1_84_4    .......... .......... .......... .......... .......... .......... -......... .......... .......... ........-. .......... .......... .......... .........A ....-..... .......... ....... 164
U1_105_4   .......... .......... .......... .......... .......... .......... -......... .......... .......... ........-. .......... .......... .......... .........A ....-..... .......... ....... 164
U1_150_4   .......... T......... .......... .......... .......... .......... -......... .......... .......... ........-. .......... .......... .......... .........A ....-..... .......... ....... 164
U1_79_4    .......... ....A..... .......... .......... .......... .......... -......... .......... .......... ........-. .......... .......... .......... .......... ....-..... .......... ....... 164
U1_86_4    .......... ....A..... .......... .......... .......... .......... -......... .......... .......... ........-. .......... .......... .......... .......... ....-..... .......... ....... 164
U1_94_4    .......... ....A..... .......... .......... .......... .......... -......... .......... .......... ........-. .......... .......... .......... .......... ....-..... .......... ....... 164
U1_138_4   .......... ....A..... .......... .......... .......... .......... -......... ......A... .......... ........-. .......... .......... .......... .......... ....-..... .......... ....... 164
U1_18_4    .......... .......... .......... .......... .......... .......... -......... .......... .....G.... ........-. .......... .......... .......... .......... ....-..... .......... ....... 164
U1_19_4    .......... .......... .......... .......... .......... .......... -......... .......... .....G.... ........-. .......... .......... .......... .......... ....-..... .......... ....... 164
U1_4_4     .......... .......... .......... .......... .......... .......... -......... .......... .....G.... ........-. .......... .......... .......... .......... ....-..... .......... ....... 164
U1_7_4     .......... .......... .......... .......... .......... .......... -......... .......... .....G.... ........-. .......... .......... .......... .......... ....-..... .......... ....... 164
U1_11_4    .......... .......... .......... .......... .......... .......... -......... .......... .....G.... ........-. .......... .......... .......... .......... ....-..... .......... ....... 164
U1_12_4    .......... .......... .......... .......... .......... .......... -......... .......... .....G.... ........-. .......... .......... .......... .......... ....-..... .......... ....... 164
U1_31_4    .......... .......... .......... .......... .......... .......... -......... .......... .....G.... ........-. .......... .......... .......... .......... ....-..... .......... ....... 164
U1_37_4    .......... .......... .......... .......... .......... .......... -......... .......... .....G.... ........-. .......... .......... .......... .......... ....-..... .......... ....... 164
U1_23_4    .......... .......... .......... .......... .......... .......... -......... .......... .....G.... ........-. .......... .......... .......... .......... ....-..... .......... ....... 164
U1_26_4    .......... .......... .......... .......... .......... .......... -......... .......... .....G.... ........-. .......... .......... .......... .......... ....-..... .......... ....... 164
U1_28_4    .......... .......... .......... .......... .......... .......... -......... .......... .....G.... ........-. .......... .......... .......... .......... ....-..... .......... ....... 164
U1_21_4    .......... .......... .......... .......... .......... .......... -......... .......... .....G..G. ........-. .......... .......... .......... .......... ....-..... .......... ....... 164
U1_24_4    .......... .......... .......... .......... .......... .......... -......... .......... .....G.... ........-. .......... .......... A......... .......... ....-..... .......... ....... 164
U1_30_4    .......... .......... .......... .......... .......... .......... -......... .......... .....G.... ........-. .......... .......... A......... .......... ....-..... .......... ....... 164
U1_3_4     .......... .......... .......... .......... .......... .......... -......... .......... .......... ........-. .......... .......... .......... .......... ....-..... .......... ......T 164
U1_6_4     .......... .......... .......... .......... .......... .......... -......... .......... .......... ........-. .......... .......... .......... .......... ....-..... .......... ......T 164
U1_34_4    .......... .......... .......... .......... .......... .......... -......... .......... .......... ........-. .......... .......... ......T... .......... ....-..... .......... ....... 164
U1_92_4    ...A...... .......... .......... .......... .......... .......... -......... .......... .......... ........-. .......... .......... .......... .......... ....-..... .......... ....... 164
U1_20_4    .......... .......... .......... .......... .......... .......... -......... .......... .......... ........-. .......... .......... .......... .......... ....-..... .......... ....... 164
U1_25_4    .......... .......... .......... .......... .......... .......... -......... .......... .......... ........-. .......... .......... .......... .......... ....-..... .......... ....... 164
U1_32_4    .......... .......... .......... .......... .......... .......... -......... .......... .......... ........-. .......... .......... .......... .......... ....-..... .......... ....... 164
U1_33_4    .......... .......... .......... .......... .......... .......... -......... .......... .......... ........-. .......... .......... .......... .......... ....-..... .......... ....... 164
U1_36_4    .......... .......... .......... .......... .......... .......... -......... .......... .......... ........-. .......... .......... .......... .......... ....-..... .......... ....... 164
U1_38_4    .......... .......... .......... .......... .......... .......... -......... .......... .......... ........-. .......... .......... .......... .......... ....-..... .......... ....... 164
U1_40_4    .......... .......... .......... .......... .......... .......... -......... .......... .......... ........-. .......... .......... .......... .......... ....-..... .......... ....... 164
U1_41_4    .......... .......... .......... .......... .......... .......... -......... .......... .......... ........-. .......... .......... .......... .......... ....-..... .......... ....... 164
U1_42_4    .......... .......... .......... .......... .......... .......... -......... .......... .......... ........-. .......... .......... .......... .......... ....-..... .......... ....... 164
U1_43_4    .......... .......... .......... .......... .......... .......... -......... .......... .......... ........-. .......... .......... .......... .......... ....-..... .......... ....... 164
U1_44_4    .......... .......... .......... .......... .......... .......... -......... .......... .......... ........-. .......... .......... .......... .......... ....-..... .......... ....... 164
U1_45_4    .......... .......... .......... .......... .......... .......... -......... .......... .......... ........-. .......... .......... .......... .......... ....-..... .......... ....... 164
U1_46_4    .......... .......... .......... .......... .......... .......... -......... .......... .......... ........-. .......... .......... .......... .......... ....-..... .......... ....... 164
U1_47_4    .......... .......... .......... .......... .......... .......... -......... .......... .......... ........-. .......... .......... .......... .......... ....-..... .......... ....... 164
U1_48_4    .......... .......... .......... .......... .......... .......... -......... .......... .......... ........-. .......... .......... .......... .......... ....-..... .......... ....... 164
U1_49_4    .......... .......... .......... .......... .......... .......... -......... .......... .......... ........-. .......... .......... .......... .......... ....-..... .......... ....... 164
U1_50_4    .......... .......... .......... .......... .......... .......... -......... .......... .......... ........-. .......... .......... .......... .......... ....-..... .......... ....... 164
U1_51_4    .......... .......... .......... .......... .......... .......... -......... .......... .......... ........-. .......... .......... .......... .......... ....-..... .......... ....... 164
U1_52_4    .......... .......... .......... .......... .......... .......... -......... .......... .......... ........-. .......... .......... .......... .......... ....-..... .......... ....... 164
U1_53_4    .......... .......... .......... .......... .......... .......... -......... .......... .......... ........-. .......... .......... .......... .......... ....-..... .......... ....... 164
U1_54_4    .......... .......... .......... .......... .......... .......... -......... .......... .......... ........-. .......... .......... .......... .......... ....-..... .......... ....... 164
U1_55_4    .......... .......... .......... .......... .......... .......... -......... .......... .......... ........-. .......... .......... .......... .......... ....-..... .......... ....... 164
U1_56_4    .......... .......... .......... .......... .......... .......... -......... .......... .......... ........-. .......... .......... .......... .......... ....-..... .......... ....... 164
U1_57_4    .......... .......... .......... .......... .......... .......... -......... .......... .......... ........-. .......... .......... .......... .......... ....-..... .......... ....... 164
U1_58_4    .......... .......... .......... .......... .......... .......... -......... .......... .......... ........-. .......... .......... .......... .......... ....-..... .......... ....... 164
U1_59_4    .......... .......... .......... .......... .......... .......... -......... .......... .......... ........-. .......... .......... .......... .......... ....-..... .......... ....... 164
U1_60_4    .......... .......... .......... .......... .......... .......... -......... .......... .......... ........-. .......... .......... .......... .......... ....-..... .......... ....... 164
U1_61_4    .......... .......... .......... .......... .......... .......... -......... .......... .......... ........-. .......... .......... .......... .......... ....-..... .......... ....... 164
U1_65_4    .......... .......... .......... .......... .......... .......... -......... .......... .......... ........-. .......... .......... .......... .......... ....-..... .......... ....... 164
U1_68_4    .......... .......... .......... .......... .......... .......... -......... .......... .......... ........T. .......... .......... .......... .......... ....-..... .......... ....... 165
U1_72_4    .......... .......... .......... .......... .......... .......... -......... .......... .......... ........-. .......... .......... .......... .......... ....-..... .......... ....... 164
U1_73_4    .......... .......... .......... .......... .......... .......... -......... .......... .......... ........-. .......... .......... .......... .......... ....-..... .......... ....... 164
U1_74_4    .......... .......... .......... .......... .......... .......... -......... .......... .......... ........-. .......... .......... .......... .......... ....-..... .......... ....... 164
U1_75_4    .......... .......... .......... .......... .......... .......... -......... .......... .......... ........-. .......... .......... .......... .......... ....-..... .......... ....... 164
U1_76_4    .......... .......... .......... .......... .......... .......... -......... .......... .......... ........-. .......... .......... .......... .......... ....-..... .......... ....... 164
U1_81_4    .......... .......... .......... .......... .......... .......... -......... .......... .......... ........-. .......... .......... .......... .......... ....-..... .......... ....... 164
U1_87_4    .......... .......... .......... .......... .......... .......... -......... .......... .......... ........-. .......... .......... .......... .......... ....-..... .......... ....... 164
U1_88_4    .......... .......... .......... .......... .......... .......... -......... .......... .......... ........-. .......... .......... .......... .......... ....-..... .......... ....... 164
U1_89_4    .......... .......... .......... .......... .......... .......... -......... .......... .......... ........-. .......... .......... .......... .......... ....-..... .......... ....... 164
U1_97_4    .......... .......... .......... .......... .......... .......... -......... .......... .......... ........-. .......... .......... .......... .......... ....-..... .......... ....... 164
U1_98_4    .......... .......... .......... .......... .......... .......... -......... .......... .......... ........-. .......... .......... .......... .......... ....-..... .......... ....... 164
U1_101_4   .......... .......... .......... .......... .......... .......... -......... .......... .......... ........-. .......... .......... .......... .......... ....-..... .......... ....... 164
U1_106_4   .......... .......... .......... .......... .......... .......... -......... .......... .......... ........-. .......... .......... .......... .......... ....-..... .......... ....... 164
U1_125_4   .......... .......... .......... .......... .......... .......... -......... .......... .......... ........-. .......... .......... .......... .......... ....-..... .......... ....... 164
U1_131_4   .......... .......... .......... .......... .......... .......... -......... .......... .......... ........-. .......... .......... .......... .......... ....-..... .......... ....... 164
U1_132_4   .......... .......... .......... .......... .......... .......... -......... .......... .......... ........-. .......... .......... .......... .......... ....-..... .......... ....... 164
U1_134_4   .......... .......... .......... .......... .......... .......... -......... .......... .......... ........-. .......... .......... .......... .......... ....-..... .......... ....... 164
U1_135_4   .......... .......... .......... .......... .......... .......... -......... .......... .......... ........-. .......... .......... .......... .......... ....-..... .......... ....... 164
```

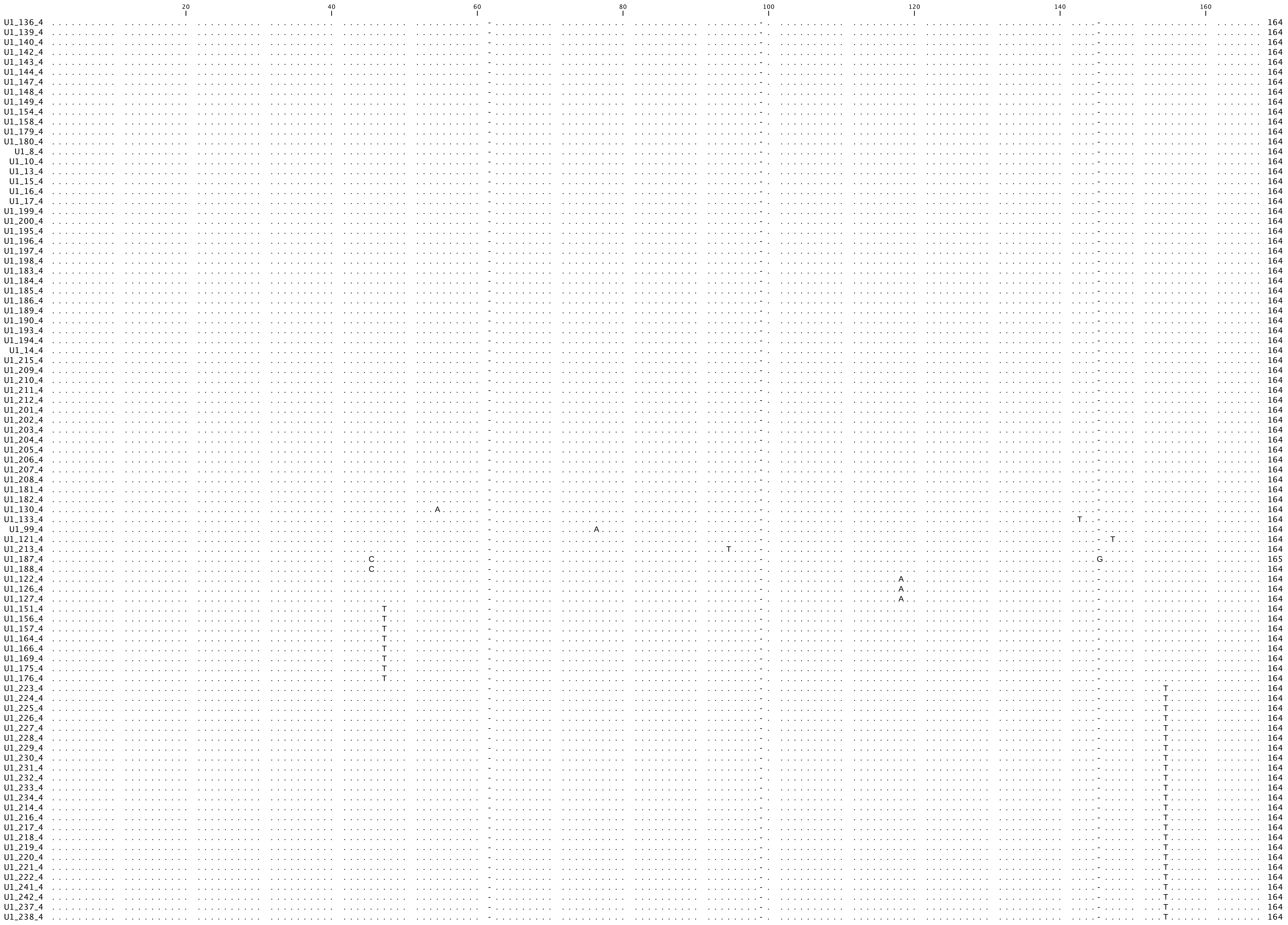
20 40 60 80 100 120 140 160
U1_136_4 164
U1_139_4 164
U1_140_4 164
U1_142_4 164
U1_143_4 164
U1_144_4 164
U1_147_4 164
U1_148_4 164
U1_149_4 164
U1_154_4 164
U1_158_4 164
U1_179_4 164
U1_180_4 164
U1_8_4 164
U1_10_4 164
U1_13_4 164
U1_15_4 164
U1_16_4 164
U1_17_4 164
U1_199_4 164
U1_200_4 164
U1_195_4 164
U1_196_4 164
U1_197_4 164
U1_198_4 164
U1_183_4 164
U1_184_4 164
U1_185_4 164
U1_186_4 164
U1_189_4 164
U1_190_4 164
U1_193_4 164
U1_194_4 164
U1_14_4 164
U1_215_4 164
U1_209_4 164
U1_210_4 164
U1_211_4 164
U1_212_4 164
U1_201_4 164
U1_202_4 164
U1_203_4 164
U1_204_4 164
U1_205_4 164
U1_206_4 164
U1_207_4 164
U1_208_4 164
U1_181_4 164
U1_182_4 164
U1_130_4 A 164
U1_133_4 T 164
U1_99_4 A 164
U1_121_4 T 164
U1_213_4 T 164
U1_187_4 C G 165
U1_188_4 C 164
U1_122_4 A 164
U1_126_4 A 164
U1_127_4 A 164
U1_151_4 T 164
U1_156_4 T 164
U1_157_4 T 164
U1_164_4 T 164
U1_166_4 T 164
U1_169_4 T 164
U1_175_4 T 164
U1_176_4 T 164
U1_223_4 T 164
U1_224_4 T 164
U1_225_4 T 164
U1_226_4 T 164
U1_227_4 T 164
U1_228_4 T 164
U1_229_4 T 164
U1_230_4 T 164
U1_231_4 T 164
U1_232_4 T 164
U1_233_4 T 164
U1_234_4 T 164
U1_214_4 T 164
U1_216_4 T 164
U1_217_4 T 164
U1_218_4 T 164
U1_219_4 T 164
U1_220_4 T 164
U1_221_4 T 164
U1_222_4 T 164
U1_241_4 T 164
U1_242_4 T 164
U1_237_4 T 164
U1_238_4 T 164

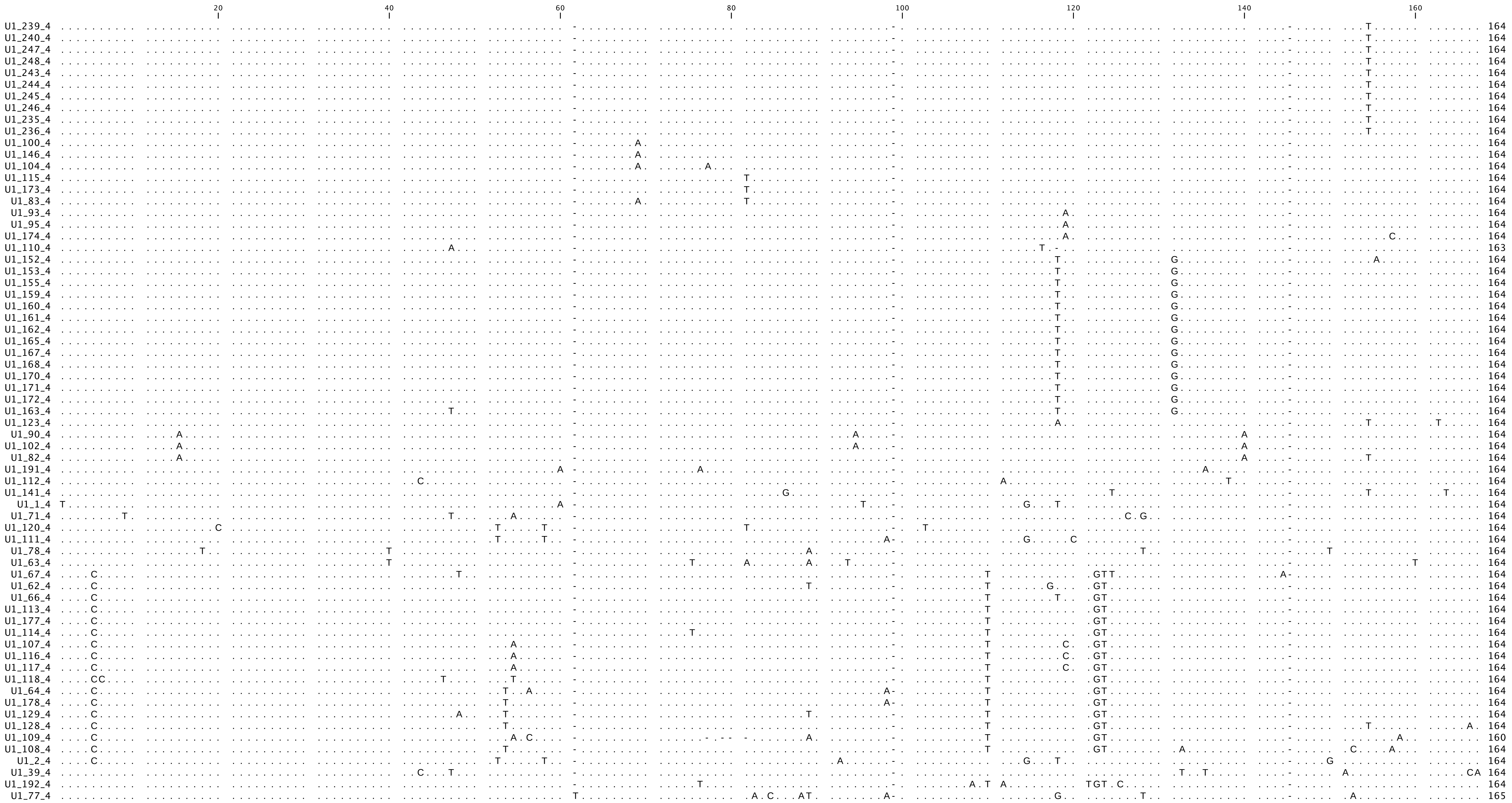
20 40 60 80 100 120 140 160
U1_239_4 164
U1_240_4 164
U1_247_4 164
U1_248_4 164
U1_243_4 164
U1_244_4 164
U1_245_4 164
U1_246_4 164
U1_235_4 164
U1_236_4 164
U1_100_4 164
U1_146_4 164
U1_104_4 164
U1_115_4 164
U1_173_4 164
U1_83_4 164
U1_93_4 164
U1_95_4 164
U1_174_4 164
U1_110_4 163
U1_152_4 164
U1_153_4 164
U1_155_4 164
U1_159_4 164
U1_160_4 164
U1_161_4 164
U1_162_4 164
U1_165_4 164
U1_167_4 164
U1_168_4 164
U1_170_4 164
U1_171_4 164
U1_172_4 164
U1_163_4 164
U1_123_4 164
U1_90_4 164
U1_102_4 164
U1_82_4 164
U1_191_4 164
U1_112_4 164
U1_141_4 164
U1_1_4 164
U1_71_4 164
U1_120_4 164
U1_111_4 164
U1_78_4 164
U1_63_4 164
U1_67_4 164
U1_62_4 164
U1_66_4 164
U1_113_4 164
U1_177_4 164
U1_114_4 164
U1_107_4 164
U1_116_4 164
U1_117_4 164
U1_118_4 164
U1_64_4 164
U1_178_4 164
U1_129_4 164
U1_128_4 164
U1_109_4 160
U1_108_4 164
U1_2_4 164
U1_39_4 164
U1_192_4 164
U1_77_4 165

```
                                20                   40                   60                   80                   100                  120                  140                  160
                                 |                    |                    |                    |                    |                    |                    |                    |
U1Somatic  ATACTTACCT GGCAGGGGAG ACACCATGAT CAAGAAGGTG GTTCACCCAG GGTGAGGCTC AGCCATTGCA CTGTCGGCTG TGTTGACCTC TGCGAATTCC CCAAATGTGG GAATCTCGAC TGCATAATTT CTGGTAGTGG GGGACTGCGT TCGCGCTCTC CCCTG 165
U1_249_7   .......... .......... .......... ..G....... .......... .......... .A........ ...C...T.. ..C.....C. .......... .......... .......... .......... .......... .......... .......... ..... 165
U1_251_7   .......... .......... .......... ..G....... .......... .......... .A........ ...C...T.. ..C.....C. .......... .......... .......... .......... .......... .......... .......... ..... 165
U1_250_7   .......... .......... .......... ..G....... .......... .......... .T........ ...C...T.. ..C.....C. .......... .......... .......... .......... .......... .......... .......... ..... 165
U1_252_7   .......... .......... .......... ..G....... .......... .......... .T........ ...C...T.. ..C.....C. .......... .......... .......... .......... .......... .......... .......... ..... 165
U1_254_8   .......... .......... .......... .......... .......... .......... .......... .......... .......... .......... .......... .......... .......... .......... .......... .......... ..... 165
U1_257_9   .......... .......... .......... .......... .......... .......... .......... .......... .......... .......... .......... .......... .......... .......... .......... .......... ..... 165
U1_255_9   .......... .......... .......... .......... .......... .......... .......... .......... .......... .......... .......... .......... .......... .......... .......... .......... ..... 165
U1_256_9   .......... .......... .......... .......... .......... .......... .......... .......... .......... .......... .......... .......... .......... .......... .......... .......... ..... 165
U1_253_8   .G........ .......... .T........ ..GA...A.. .......G.. ..C......T G......... ..-C.A.TCA A.C....TC. .......... .......... .......... ....C..... .......... ........A. .......... ..... 164
U1_259_20  T......... .......... .T........ .......... .......... ..C......T G......... ..-C....CA C.C.....C. .......... .......... .......... .......... .......... .......... .......... ..... 164
U1_258_20  .......... .......... .T........ .......... .......... ..CA.....T G......... ..-C....CA C.C.....C. .......... .......... .......... .......... .......... .......... .......... ..... 164
```

```
                        20        40        60        80        100       120       140       160       180
U2Maternal  ATCGCTTCTC GGCCTTTTGG CTAAGATCAA GTGTAGTATC TGTTCTTATC AGTTTAATAT CTGATACGTG CCCTACCCGG GCACTATATA TTAAATTGAT TTTTGCAGCT GGGAGATGGG TTAGGGGCTT GCTCCACCCA CTCCACGCAT CGACCTGGTA TTGCAGTACC TCCGGGAACG GTGCACCATT - 190
U2_1_2      .......... .......... .......... .......... .......... .......... .......... .......... .......... .......... .......... .......... .......... .......... .......... .......... .......... .......... ......TCA. T 191
U2_2_2      .......... .......... .......... .......... .......... .......... .......... .......... .......... .......... .......... .......... .......... .......... .......... .......... .......... .......... ......TCA. T 191
```

```
                                      20                  40                  60                  80                 100                 120                 140                 160                 180
                                       |                   |                   |                   |                   |                   |                   |                   |                   |
U2Somatic  ATCGCTTCTC GGCCTTTTGG CTAAGATCAA GTGTAGTATC TGTTCTTATC AGTTTAATAT CTGATACGTG CCCTACCCGG GCACCATATA TTAAATTGAT TTTTGGAACA GGGAGATGGA ATAGGGGCTT GCTCCGTCCA CTCCACGCAT CGACCCGGTA TTGCAGTACT TCCGGGAACG GTGCACCACC T 191
U2_3_6     .......... .......... .......... .......... .......... .......... .......... .......... .......... .......... .......G.. .......... .......... .......... .......... .......... .......... .......... .......C.. . 191
U2_6_13    .......... .......... .......... .......... .......... .......... .......... .......... .......... .......... .......... .......... .......... .......... .......... .......... .........C .......... .......C.. . 191
U2_7_13    .......... .......... .......... .......... .......... .......... .......... .......... .......... .......... .......... .......... .......... .......... .......... .......... .........C .......... .......C.. . 191
U2_4_7     .......... .......... .......... .......... .......... .......... .......... .......... .......... .......... ........T. .......... .......... .......... .......... .......... .......... .......... .......C.. . 191
U2_5_7     .......... .......... .......... .......... .......... .......... .......... .......... .......... .......... .......... .......... .......... .......... .......T.. .......... .........A .......... .......C.. . 191
```

```
                                     20                    40                    60                    80                    100                   120                   140
U4Maternal ATTCTTTCGC AGAGGCAATA TCATAGCCTA TGAGGCCTAG CCGAGGCGTG ATCATTGCTG GTTGAAAACT TTACCCAATA CCCCGCCGAG ATGGCTTGAA ATACAGTCAG CTCTGGCAAT TTTTGACAGC CTCCTAGAAG G 141
U4_6_17    .......... .......... ......T... .......... .......... .......... .......... .......... .......... .......... .......... .......... .......... .......... . 141
U4_4_17    .......... .......... .......... .......... .......... .......... .......... .......... .......... .......... .......... .......... .......... .......... . 141
U4_5_17    .......... .......... .......... .......... .......... .......... .......... .......... .......... .......... .......... .......... .......... .......... . 141
U4_8_17    .......... .......... .......... .......... .......... .......... .......... .......... .......... .......... .......... .......... .......... .......... . 141
U4_10_17   .......... .......... .......... .......... .......... .......... .......... .......... .......... .......... .......... .......... .......... .......... . 141
U4_14_17   .......... .......... .......... .......... .......... .......... .......... .......... .......... .......... .......... .......... .......... .......... . 141
U4_17_17   .......... .......... .......... .......... .......... .......... .......... .......... .......... .......... .......... .......... .......... .......... . 141
U4_18_17   .......... .......... .......... .......... .......... .......... .......... .......... .......... .......... .......... .......... .......... .......... . 141
U4_19_17   .......... .......... .......... .......... .......... .......... .......... .......... .......... .......... .......... .......... .......... .......... . 141
U4_20_17   .......... .......... .......... .......... .......... .......... .......... .......... .......... .......... .......... .......... .......... .......... . 141
U4_21_17   .......... .......... .......... .......... .......... .......... .......... .......... .......... .......... .......... .......... .......... .......... . 141
U4_22_17   .......... .......... .......... .......... .......... .......... .......... .......... .......... .......... .......... .......... .......... .......... . 141
U4_16_17   .......... .......... .......T.. .......... .......... .......... .......... .......... .......... .......... .......... .......... .......... .......... . 141
U4_23_17   .......... .......... .......T.. .......... .......... .......... .......... .......... .......... .......... .......... .......... .......... .......... . 141
U4_3_17    .......... .......... .......... .......... ......T... .......... .......... .......... .......... .......... .......... .......... .......... .......... . 141
U4_12_17   .......... .......... .......... .......... .......... .......... T......... .......... .......... .......... .......... .......... .......... .......... . 141
U4_24_17   .......... .......... .......... .......... ......T... .......... .......... .......... .......... .......... .......... ........G. .......... .......... . 141
U4_7_17    .......... .......... .......... .......... ......T... .......... .......C.. .......... .......... .......... .......... ........G. .......... .......... . 141
U4_9_17    .......... .......... .......... .......... .......... .......... .......... ..G....... ......T... .......... .......... .......... .......... .......... . 141
U4_13_17   .......... .........G .......... .......... .......... .......... .......... ..G....... ......T... .......... .......... .......... .......... .......... . 141
U4_11_17   .......... .A........ .......... .......... .......... .......... .......... .......... ...T...... ...T...... .......... .......... .......... .......... . 141
U4_15_17   .......... .A........ .......... .......... .......... .......... .......... .......... ...T...... ...T...... .......... .......... .......... .......... . 141
U4_25_17   ...T...... .......... .......... .......... .T....T... .......... .......... .......... .......... .........G .......... .......... .......... .......... . 141
```

```
                                      20                   40                   60                   80                   100                  120                  140
                                       |                    |                    |                    |                    |                    |                    |
U4Somatic  AGCTTTGCGC AGTGGCAGTA TCGTAGCCTA TGAGGTTTAT CCGAGGCGCG ATTATTGCTA ATTGAAAACT TTACCCAATA CCCCGCCGTG ACGACTTGAA ATATAGTCGG CACTGGCAAT TTTTGACGGT CTCGACAGAG A 141
   U4_1_4  .......... .......... .......... .......... .......... .......... .......... .......... .......... .......... .......... .......... .......... .......... . 141
 U4_26_25  .......... .......... .......... .......... .......... .......... .......... .......... .......... .......... .......... .......... .......A.. ..TCT.GA.. . 141
   U4_2_4  .T.C...... .......... ........C. .......... .......... .........G G......... .......... .......... .......... .......... .T........ .......... ...A.A.... . 141
```

```
                              20                  40                  60                  80                  100
                               |                   |                   |                   |                   |
U5Maternal ACAGCTCGAG TTTCTCTTCA ATCACGCACA AATCTTTCGC CTTTTACTAA AGATTTCCGT GGGGAGGAAC TATTGTGAGT TTTGTTTATT TTTGGGTGCT CTGCTATCTG CAGAGCTG 118
U5_8_5     -......... .......... .......... .......... .......... .......... .......... .......... .......... .......... .......... .......- 116
U5_11_5    -......... .......... .......... .......... .......... .......... .......... .......... .......... .......... .......... .......- 116
U5_12_5    -......... .......... .......... .......... .......... .......... .......... .......... .......... .......... .......... .......- 116
U5_13_5    -......... .......... .......... .......... .......... .......... .......... .......... .......... .......... .......... .......- 116
U5_14_5    -......... .......... .......... .......... .......... .......... .......... .......... .......... .......... .......... .......- 116
U5_15_5    -......... .......... .......... .......... .......... .......... .......... .......... .......... .......... .......... .......- 116
U5_16_5    -......... .......... .......... .......... .......... .......... .......... .......... .......... .......... .......... .......- 116
U5_17_5    -......... .......... .......... .......... .......... .......... .......... .......... .......... .......... .......... .......- 116
U5_18_5    -......... .......... .......... .......... .......... .......... .......... .......... .......... .......... .......... .......- 116
U5_19_5    -......... .......... .......... .......... .......... .......... .......... .......... .......... .......... .......... .......- 116
U5_20_5    -......... .......... .......... .......... .......... .......... .......... .......... .......... .......... .......... .......- 116
U5_21_5    -......... .......... .......... .......... .......... .......... .......... .......... .......... .......... .......... .......- 116
U5_22_5    -......... .......... .......... .......... .......... .......... .......... .......... .......... .......... .......... .......- 116
U5_23_5    -......... .......... .......... .......... .......... .......... .......... .......... .......... .......... .......... .......- 116
U5_24_5    -......... .......... .......... .......... .......... .......... .......... .......... .......... .......... .......... .......- 116
U5_25_5    -......... .......... .......... .......... .......... .......... .......... .......... .......... .......... .......... .......- 116
U5_5_5     -......... .......... .......... .......... .......... .......... .......... .......... .......... .......... .......... .......- 116
U5_6_5     -......... .......... .......... .......... .......... .......... .......... .......... .......... .......... .......... .......- 116
U5_7_5     -......... .......... .......... .......... .......... .......... .......... .......... .......... .......... .......... .......- 116
U5_129_5   -......... .......... .......... .......... .......... .......... .......... .......... .......... .......... .......... .......- 116
U5_131_5   -......... .......... .......... .......... .......... .......... .......... .......... .......... .......... .......... .......- 116
U5_125_5   -......... .......... .......... .......... .......... .......... .......... .......... .......... .......... .......... .......- 116
U5_126_5   -......... .......... .......... .......... .......... .......... .......... .......... .......... .......... .......... .......- 116
U5_127_5   -......... .......... .......... .......... .......... .......... .......... .......... .......... .......... .......... .......- 116
U5_128_5   -......... .......... .......... .......... .......... .......... .......... .......... .......... .......... .......... .......- 116
U5_26_5    -......... .......... .......... .......... .......... .......... .......... .......... .......... .......... .......... .......- 116
U5_27_5    -......... .......... .......... .......... .......... .......... .......... .......... .......... .......... .......... .......- 116
U5_28_5    -......... .......... .......... .......... .......... .......... .......... .......... .......... .......... .......... .......- 116
U5_29_5    -......... .......... .......... .......... .......... .......... .......... .......... .......... .......... .......... .......- 116
U5_30_5    -......... .......... .......... .......... .......... .......... .......... .......... .......... .......... .......... .......- 116
U5_31_5    -......... .......... .......... .......... .......... .......... .......... .......... .......... .......... .......... .......- 116
U5_32_5    -......... .......... .......... .......... .......... .......... .......... .......... .......... .......... .......... .......- 116
U5_33_5    -......... .......... .......... .......... .......... .......... .......... .......... .......... .......... .......... .......- 116
U5_34_5    -......... .......... .......... .......... .......... .......... .......... .......... .......... .......... .......... .......- 116
U5_35_5    -......... .......... .......... .......... .......... .......... .......... .......... .......... .......... .......... .......- 116
U5_36_5    -......... .......... .......... .......... .......... .......... .......... .......... .......... .......... .......... .......- 116
U5_37_5    -......... .......... .......... .......... .......... .......... .......... .......... .......... .......... .......... .......- 116
U5_38_5    -......... .......... .......... .......... .......... .......... .......... .......... .......... .......... .......... .......- 116
U5_39_5    -......... .......... .......... .......... .......... .......... .......... .......... .......... .......... .......... .......- 116
U5_40_5    -......... .......... .......... .......... .......... .......... .......... .......... .......... .......... .......... .......- 116
U5_41_5    -......... .......... .......... .......... .......... .......... .......... .......... .......... .......... .......... .......- 116
U5_42_5    -......... .......... .......... .......... .......... .......... .......... .......... .......... .......... .......... .......- 116
U5_43_5    -......... .......... .......... .......... .......... .......... .......... .......... .......... .......... .......... .......- 116
U5_44_5    -......... .......... .......... .......... .......... .......... .......... .......... .......... .......... .......... .......- 116
U5_45_5    -......... .......... .......... .......... .......... .......... .......... .......... .......... .......... .......... .......- 116
U5_46_5    -......... .......... .......... .......... .......... .......... .......... .......... .......... .......... .......... .......- 116
U5_47_5    -......... .......... .......... .......... .......... .......... .......... .......... .......... .......... .......... .......- 116
U5_48_5    -......... .......... .......... .......... .......... .......... .......... .......... .......... .......... .......... .......- 116
U5_49_5    -......... .......... .......... .......... .......... .......... .......... .......... .......... .......... .......... .......- 116
U5_50_5    -......... .......... .......... .......... .......... .......... .......... .......... .......... .......... .......... .......- 116
U5_51_5    -......... .......... .......... .......... .......... .......... .......... .......... .......... .......... .......... .......- 116
U5_52_5    -......... .......... .......... .......... .......... .......... .......... .......... .......... .......... .......... .......- 116
U5_53_5    -......... .......... .......... .......... .......... .......... .......... .......... .......... .......... .......... .......- 116
U5_54_5    -......... .......... .......... .......... .......... .......... .......... .......... .......... .......... .......... .......- 116
U5_55_5    -......... .......... .......... .......... .......... .......... .......... .......... .......... .......... .......... .......- 116
U5_56_5    -......... .......... .......... .......... .......... .......... .......... .......... .......... .......... .......... .......- 116
U5_57_5    -......... .......... .......... .......... .......... .......... .......... .......... .......... .......... .......... .......- 116
U5_58_5    -......... .......... .......... .......... .......... .......... .......... .......... .......... .......... .......... .......- 116
U5_59_5    -......... .......... .......... .......... .......... .......... .......... .......... .......... .......... .......... .......- 116
U5_60_5    -......... .......... .......... .......... .......... .......... .......... .......... .......... .......... .......... .......- 116
U5_61_5    -......... .......... .......... .......... .......... .......... .......... .......... .......... .......... .......... .......- 116
U5_62_5    -......... .......... .......... .......... .......... .......... .......... .......... .......... .......... .......... .......- 116
U5_63_5    -......... .......... .......... .......... .......... .......... .......... .......... .......... .......... .......... .......- 116
U5_64_5    -......... .......... .......... .......... .......... .......... .......... .......... .......... .......... .......... .......- 116
U5_65_5    -......... .......... .......... .......... .......... .......... .......... .......... .......... .......... .......... .......- 116
U5_66_5    -......... .......... .......... .......... .......... .......... .......... .......... .......... .......... .......... .......- 116
U5_67_5    -......... .......... .......... .......... .......... .......... .......... .......... .......... .......... .......... .......- 116
U5_68_5    -......... .......... .......... .......... .......... .......... .......... .......... .......... .......... .......... .......- 116
U5_69_5    -......... .......... .......... .......... .......... .......... .......... .......... .......... .......... .......... .......- 116
U5_70_5    -......... .......... .......... .......... .......... .......... .......... .......... .......... .......... .......... .......- 116
U5_71_5    -......... .......... .......... .......... .......... .......... .......... .......... .......... .......... .......... .......- 116
U5_72_5    -......... .......... .......... .......... .......... .......... .......... .......... .......... .......... .......... .......- 116
U5_73_5    -......... .......... .......... .......... .......... .......... .......... .......... .......... .......... .......... .......- 116
U5_74_5    -......... .......... .......... .......... .......... .......... .......... .......... .......... .......... .......... .......- 116
U5_75_5    -......... .......... .......... .......... .......... .......... .......... .......... .......... .......... .......... .......- 116
U5_77_5    -......... .......... .......... .......... .......... .......... .......... .......... .......... .......... .......... .......- 116
U5_78_5    -......... .......... .......... .......... .......... .......... .......... .......... .......... .......... .......... .......- 116
U5_80_5    -......... .......... .......... .......... .......... .......... .......... .......... .......... .......... .......... .......- 116
U5_81_5    -......... .......... .......... .......... .......... .......... .......... .......... .......... .......... .......... .......- 116
U5_83_5    -......... .......... .......... .......... .......... .......... .......... .......... .......... .......... .......... .......- 116
U5_84_5    -......... .......... .......... .......... .......... .......... .......... .......... .......... .......... .......... .......- 116
U5_85_5    -......... .......... .......... .......... .......... .......... .......... .......... .......... .......... .......... .......- 116
U5_86_5    -......... .......... .......... .......... .......... .......... .......... .......... .......... .......... .......... .......- 116
U5_87_5    -......... .......... .......... .......... .......... .......... .......... .......... .......... .......... .......... .......- 116
U5_88_5    -......... .......... .......... .......... .......... .......... .......... .......... .......... .......... .......... .......- 116
U5_89_5    -......... .......... .......... .......... .......... .......... .......... .......... .......... .......... .......... .......- 116
U5_90_5    -......... .......... .......... .......... .......... .......... .......... .......... .......... .......... .......... .......- 116
U5_91_5    -......... .......... .......... .......... .......... .......... .......... .......... .......... .......... .......... .......- 116
U5_92_5    -......... .......... .......... .......... .......... .......... .......... .......... .......... .......... .......... .......- 116
U5_93_5    -......... .......... .......... .......... .......... .......... .......... .......... .......... .......... .......... .......- 116
```

```
                                     20         40         60         80         100
                                      |          |          |          |          |
U5_94_5   -......... .......... .......... .......... .......... .......... .......... .......... .......... .......... .......... .......-  116
U5_95_5   -......... .......... .......... .......... .......... .......... .......... .......... .......... .......... .......... .......-  116
U5_96_5   -......... .......... .......... .......... .......... .......... .......... .......... .......... .......... .......... .......-  116
U5_97_5   -......... .......... .......... .......... .......... .......... .......... .......... .......... .......... .......... .......-  116
U5_98_5   -......... .......... .......... .......... .......... .......... .......... .......... .......... .......... .......... .......-  116
U5_99_5   -......... .......... .......... .......... .......... .......... .......... .......... .......... .......... .......... .......-  116
U5_100_5  -......... .......... .......... .......... .......... .......... .......... .......... .......... .......... .......... .......-  116
U5_101_5  -......... .......... .......... .......... .......... .......... .......... .......... .......... .......... .......... .......-  116
U5_102_5  -......... .......... .......... .......... .......... .......... .......... .......... .......... .......... .......... .......-  116
U5_103_5  -......... .......... .......... .......... .......... .......... .......... .......... .......... .......... .......... .......-  116
U5_104_5  -......... .......... .......... .......... .......... .......... .......... .......... .......... .......... .......... .......-  116
U5_105_5  -......... .......... .......... .......... .......... .......... .......... .......... .......... .......... .......... .......-  116
U5_106_5  -......... .......... .......... .......... .......... .......... .......... .......... .......... .......... .......... .......-  116
U5_107_5  -......... .......... .......... .......... .......... .......... .......... .......... .......... .......... .......... .......-  116
U5_108_5  -......... .......... .......... .......... .......... .......... .......... .......... .......... .......... .......... .......-  116
U5_109_5  -......... .......... .......... .......... .......... .......... .......... .......... .......... .......... .......... .......-  116
U5_110_5  -......... .......... .......... .......... .......... .......... .......... .......... .......... .......... .......... .......-  116
U5_111_5  -......... .......... .......... .......... .......... .......... .......... .......... .......... .......... .......... .......-  116
U5_112_5  -......... .......... .......... .......... .......... .......... .......... .......... .......... .......... .......... .......-  116
U5_113_5  -......... .......... .......... .......... .......... .......... .......... .......... .......... .......... .......... .......-  116
U5_114_5  -......... .......... .......... .......... .......... .......... .......... .......... .......... .......... .......... .......-  116
U5_115_5  -......... .......... .......... .......... .......... .......... .......... .......... .......... .......... .......... .......-  116
U5_116_5  -......... .......... .......... .......... .......... .......... .......... .......... .......... .......... .......... .......-  116
U5_117_5  -......... .......... .......... .......... .......... .......... .......... .......... .......... .......... .......... .......-  116
U5_118_5  -......... .......... .......... .......... .......... .......... .......... .......... .......... .......... .......... .......-  116
U5_119_5  -......... .......... .......... .......... .......... .......... .......... .......... .......... .......... .......... .......-  116
U5_120_5  -......... .......... .......... .......... .......... .......... .......... .......... .......... .......... .......... .......-  116
U5_121_5  -......... .......... .......... .......... .......... .......... .......... .......... .......... .......... .......... .......-  116
U5_122_5  -......... .......... .......... .......... .......... .......... .......... .......... .......... .......... .......... .......-  116
U5_123_5  -......... .......... .......... .......... .......... .......... .......... .......... .......... .......... .......... .......-  116
U5_124_5  -......... .......... .......... .......... .......... .......... .......... .......... .......... .......... .......... .......-  116
U5_4_5    -......A.. .......... .......... .......... .......... .......... .......... .......... .......... .......... .......... .......-  116
U5_82_5   -......... .......... .......... .......... .......... .......... .........T .......... .......... .......... .......... .......-  116
U5_130_5  -......... .......... .......... .......... .......... .......... ........G. .......... .......... .......... .......... .......-  116
U5_10_5   -......... .......A.. .......... .......... .......... .......... .......... .......... .......... .......... .......... .......-  116
U5_9_5    -......... .......... .......... .......... .......... .T........ .......... .......... .......... .......... .......... .......-  116
U5_79_5   -......... .......... .......... .......... .......... .T........ .......... .......... .......... .......... .......... .......-  116
U5_1_5    -......... .......... .......... .......... .......... .......... .......... ........A. .......... .......... .......... .......-  116
U5_2_5    -......... .......... .....T.... .......... .......... .......... .......... ........A. .......... .......... .......... .......-  116
U5_3_5    -......... .......... .A........ .......... .......... .......... .......... .......... .A........ .......... .......... .......-  116
U5_76_5   -.....T... .....G.... .......... .......... .......... .......... .......... .......... .......... .......... .......... .......-  116
U5_132_5  -T.....A.. .......... .......T.. .........T T......... .......... .......... .......... .......... .......... .......... .......-  116
```

```
                                 20                   40                   60                   80                   100
                                  |                    |                    |                    |                    |
U5_134_20  --ACACTCAT GTTTTTCCTC AGATCGTATA AATCTTTCGC CTTTTACTAA AGATTTCCGT GGGGAAGAAC ACTATGAGTA TAAACTAATT TTTTGATGCC CTGCCGATAG GCGGGGCT 116
U5Somatic  GT........ .......... .......... .......... .......... .......... .......... .......... .......... .......... .......C.. ........ 118
U5_135_20  --........ ....CA.... .......... .......... .......... .......... .....G.... ..A....... .T........ .......... ....AACAT. CG...C.- 115
U5_136_20  --........ ....C..T.. .......... .......... .......... .......... .....G.... ..A....... ...C...... .......... .....T..C. .T...... 116
U5_139_20  --.......A ....C..T.. .......... .......... .......... .......... .....G.... T.A....... ...T...... .......... .....T..C. .T...... 116
U5_133_9   --........ ....C..T.. .T...A.... .G........ .......... ........A. ..A..G.... ..CT...... .TT....... ........T. ...GTTCA.T .TA..AT. 116
U5_138_20  --........ ....C..T.. .......... .......... .......... .......... .....G.... ..A....... ...T...... ...GATGC.T TGCTTATACC ........ 116
U5_137_20  -AG.T..TGA C......T.. .......... .......... .......... .......... .....G.... ..A....... .......... ...GATGC.. TGCTTATACC .......- 116
```

```
                              20                    40                    60                    80                   100
                               |                     |                     |                     |                     |
U6Maternal GTGCTCGCTA CGGTGGCACA TATACTAAAA TTGGATCGAT ACAGAGAAGA TTAGCATGGC CCCTGCGAAA GGATGACACG CAAATCCGTG AAGCGCTCCA TATTTCT 107
  U6_276_4 -GTG...... .......... .......... .......... .......... .........T .......... ........T. .......... .......... ....G.. 106
  U6_341_4 ..A....... .......... .......... .........A .......... .......... .......... .......... .......... .......... ....--- 104
  U6_343_4 ..A....... .......... .......... .........A .......... .......... .......... .......... .......... .......... ....--- 104
  U6_294_4 .......... .......... .......... .......... ........T. .......... .......... .......... .......... .......T.. ....G.. 107
  U6_302_4 .......... .......... .......... .......... ........T. .......... .......... .......... .......... .......T.. ....G.. 107
  U6_306_4 .......... .......... .......... .......... ........T. .......... .......... .......... .......... .......T.. ....G.. 107
  U6_310_4 .......... .......... .......... .......... ........T. .......... .......... .......... .......... .......T.. ....G.. 107
    U6_8_3 .......... .......... .......... .......... .....A.... .......... .......... .......... .......... .......... ....--- 104
  U6_281_4 .......... .......... .......... ...C...... .....A.... .......... .......... .......... .......... .......... ....G.. 107
   U6_85_4 .......... .......... .......... .......... .......... .......... .......... ..G....... .......... .......... ....--- 104
  U6_101_4 .......... .......... .......... .......... .......... .......... .......... ..G....... .......... .......... ....--- 104
  U6_121_4 .....T.... .......... .......... .......... .......... .......... .......... ..G....... .......... .......... ....--- 104
  U6_172_4 .......... .......... .......... ...C...... .......... .......... .......... .......... .......... .......... ....... 107
  U6_503_4 .......... .......... .......... ...C...... .......... .......... .......... .......... .......... .......... ....G.. 107
  U6_507_4 .......... .......... .......... ...C...... .......... .......... .......... .......... .......... .......... ....G.. 107
  U6_259_4 .......... ......T... .......... .......... .......... .......... .......... .......... .......... .......... ....G.. 107
  U6_264_4 .......... ......T... .......... .......... .......... .......... .......... .......... .......... .......... ....G.. 107
  U6_297_4 .......... ......G... .......... .......... .......... .......... .......... .......... .......... .......... ....G.. 107
  U6_301_4 .......... ......G... .......... .......... .......... .......... .......... .......... .......... .......... ....G.. 107
  U6_313_4 .......... ......G... .......... .......... .......... .......... .......... .......... .......... .......... ....G.. 107
  U6_305_4 .......... ......G... .......... .......... .......... .......... .......... .......... .......... .......... ....G.. 107
  U6_309_4 .......... ......G... .......... .......... .......... .......... .......... .......... .......... .......... ....G.. 107
   U6_25_4 .......... .......... .......... .......... .......... .......... .......... .........A .......... .......... ....G.. 107
  U6_245_4 .......... .......... .......... .......... .......... .......... .......... .........T .......... .......... ....G.. 107
  U6_146_4 .......... .......... .T........ .......... .......... .......... .......... .......... .......... .......... ....G.. 107
  U6_165_4 .......... .......... .T........ .......... .......... .......... .......... .......... .......... .......... ....G.. 107
  U6_269_4 .......... .......... .......... .......... .......... .......... .......... .......... .......... G......... ....G.. 107
  U6_179_4 .......... .......... .......... .......... .......... ........A. .......... .......... .......... .......... ....--- 104
  U6_185_4 .......... .......... .......... .......... .......... ........A. .......... .......... .......... .......... ....--- 104
  U6_436_4 .......... .......... .......... .......... .......... ........T. .......... .......... .......... .......... ....G.. 107
  U6_125_4 .......... .......... .......... ......T... .......... .......... .......... .......... .......... .......... ....--- 104
    U6_3_3 .......... .......... .......... .......... .......... .......... .......... .......... .......... .......... ....--- 104
    U6_4_3 .......... .......... .......... .......... .......... .......... .......... .......... .......... .......... ....--- 104
    U6_5_3 .......... .......... .......... .......... .......... .......... .......... .......... .......... .......... ....--- 104
   U6_84_4 .......... .......... .......... .......... .......... .......... .......... .......... .......... .......... ....--- 104
   U6_88_4 .......... .......... .......... .......... .......... .......... .......... .......... .......... .......... ....--- 104
   U6_90_4 .......... .......... .......... .......... .......... .......... .......... .......... .......... .......... ....--- 104
   U6_93_4 .......... .......... .......... .......... .......... .......... .......... .......... .......... .......... ....--- 104
  U6_111_4 .......... .......... .......... .......... .......... .......... .......... .......... .......... .......... ....--- 104
  U6_112_4 .......... .......... .......... .......... .......... .......... .......... .......... .......... .......... ....--- 104
  U6_114_4 .......... .......... .......... .......... .......... .......... .......... .......... .......... .......... ....--- 104
  U6_116_4 .......... .......... .......... .......... .......... .......... .......... .......... .......... .......... ....--- 104
  U6_117_4 .......... .......... .......... .......... .......... .......... .......... .......... .......... .......... ....--- 104
  U6_119_4 .......... .......... .......... .......... .......... .......... .......... .......... .......... .......... ....--- 104
  U6_120_4 .......... .......... .......... .......... .......... .......... .......... .......... .......... .......... ....--- 104
  U6_122_4 .......... .......... .......... .......... .......... .......... .......... .......... .......... .......... ....--- 104
  U6_283_4 .......... .......... .......... .......... .......... .......... .......... .......... .......... .......... ....... 107
  U6_286_4 .......... .......... .......... .......... .......... .......... .......... .......... .......... .......... ....... 107
  U6_288_4 .......... .......... .......... .......... .......... .......... .......... .......... .......... .......... ....... 107
  U6_315_4 .......... .......... .......... .......... .......... .......... .......... .......... .......... .......... ....--- 104
  U6_323_4 .......... .......... .......... .......... .......... .......... .......... .......... .......... .......... ....--- 104
  U6_325_4 .......... .......... .......... .......... .......... .......... .......... .......... .......... .......... ....--- 104
  U6_327_4 .......... .......... .......... .......... .......... .......... .......... .......... .......... .......... ....--- 104
  U6_331_4 .......... .......... .......... .......... .......... .......... .......... .......... .......... .......... ....--- 104
  U6_340_4 .......... .......... .......... .......... .......... .......... .......... .......... .......... .......... ....--- 104
  U6_344_4 .......... .......... .......... .......... .......... .......... .......... .......... .......... .......... ....--- 104
  U6_345_4 .......... .......... .......... .......... .......... .......... .......... .......... .......... .......... ....--- 104
  U6_346_4 .......... .......... .......... .......... .......... .......... .......... .......... .......... .......... ....--- 104
  U6_347_4 .......... .......... .......... .......... .......... .......... .......... .......... .......... .......... ....--- 104
  U6_348_4 .......... .......... .......... .......... .......... .......... .......... .......... .......... .......... ....--- 104
  U6_353_4 .......... .......... .......... .......... .......... .......... .......... .......... .......... .......... ....--- 104
  U6_360_4 .......... .......... .......... .......... .......... .......... .......... .......... .......... .......... ....--- 104
   U6_33_4 .......... .......... .......... .......... .......... .......... .......... .......... .......... .......... ....G.. 107
   U6_35_4 .......... .......... .......... .......... .......... .......... .......... .......... .......... .......... ..----- 102
   U6_39_4 .......... .......... .......... .......... .......... .......... .......... .......... .......... .......... ....G.. 107
   U6_40_4 .......... .......... .......... .......... .......... .......... .......... .......... .......... .......... ....G.. 107
   U6_43_4 .......... .......... .......... .......... .......... .......... .......... .......... .......... .......... ....G.. 107
   U6_45_4 .......... .......... .......... .......... .......... .......... .......... .......... .......... .......... ....G.. 107
   U6_46_4 .......... .......... .......... .......... .......... .......... .......... .......... .......... .......... ....G.. 107
   U6_47_4 .......... .......... .......... .......... .......... .......... .......... .......... .......... .......... ....G.. 107
   U6_50_4 .......... .......... .......... .......... .......... .......... .......... .......... .......... .......... ....G.. 107
   U6_52_4 .......... .......... .......... .......... .......... .......... .......... .......... .......... .......... ..----- 102
   U6_55_4 .......... .......... .......... .......... .......... .......... .......... .......... .......... .......... ....G.. 107
   U6_56_4 .......... .......... .......... .......... .......... .......... .......... .......... .......... .......... ..----- 102
   U6_58_4 .......... .......... .......... .......... .......... .......... .......... .......... .......... .......... ....G.. 107
   U6_59_4 .......... .......... .......... .......... .......... .......... .......... .......... .......... .......... ....G.. 107
   U6_62_4 .......... .......... .......... .......... .......... .......... .......... .......... .......... .......... ....G.. 107
   U6_64_4 .......... .......... .......... .......... .......... .......... .......... .......... .......... .......... ....G.. 107
   U6_65_4 .......... .......... .......... .......... .......... .......... .......... .......... .......... .......... ....--- 104
   U6_67_4 .......... .......... .......... .......... .......... .......... .......... .......... .......... .......... ....G.. 107
   U6_68_4 .......... .......... .......... .......... .......... .......... .......... .......... .......... .......... ....G.. 107
   U6_70_4 .......... .......... .......... .......... .......... .......... .......... .......... .......... .......... ....G.. 107
   U6_71_4 .......... .......... .......... .......... .......... .......... .......... .......... .......... .......... ....G.. 107
   U6_73_4 .......... .......... .......... .......... .......... .......... .......... .......... .......... .......... ....--- 104
   U6_79_4 .......... .......... .......... .......... .......... .......... .......... .......... .......... .......... ....--- 104
   U6_81_4 .......... .......... .......... .......... .......... .......... .......... .......... .......... .......... ....--- 104
   U6_96_4 .......... .......... .......... .......... .......... .......... .......... .......... .......... .......... ....--- 104
   U6_98_4 .......... .......... .......... .......... .......... .......... .......... .......... .......... .......... ....--- 104
  U6_100_4 .......... .......... .......... .......... .......... .......... .......... .......... .......... .......... ....--- 104
  U6_103_4 .......... .......... .......... .......... .......... .......... .......... .......... .......... .......... ....--- 104
```

```
                            20         40         60         80         100
                            |          |          |          |          |
U6_106_4 .......... .......... .......... .......... .......... .......... .......... .......... .......... .......... ....--- 104
U6_107_4 .......... .......... .......... .......... .......... .......... .......... .......... .......... .......... ....--- 104
U6_109_4 .......... .......... .......... .......... .......... .......... .......... .......... .......... .......... ....--- 104
U6_110_4 .......... .......... .......... .......... .......... .......... .......... .......... .......... .......... ....--- 104
U6_124_4 .......... .......... .......... .......... .......... .......... .......... .......... .......... .......... ....--- 104
U6_126_4 .......... .......... .......... .......... .......... .......... .......... .......... .......... .......... ....--- 104
U6_127_4 .......... .......... .......... .......... .......... .......... .......... .......... .......... .......... ....--- 104
U6_129_4 .......... .......... .......... .......... .......... .......... .......... .......... .......... .......... ....--- 104
U6_131_4 .......... .......... .......... .......... .......... .......... .......... .......... .......... .......... ....--- 104
U6_133_4 .......... .......... .......... .......... .......... .......... .......... .......... .......... .......... ....G.. 107
U6_135_4 .......... .......... .......... .......... .......... .......... .......... .......... .......... .......... ....G.. 107
U6_139_4 .......... .......... .......... .......... .......... .......... .......... .......... .......... .......... ....G.. 107
U6_141_4 .......... .......... .......... .......... .......... .......... .......... .......... .......... .......... ....G.. 107
U6_142_4 .......... .......... .......... .......... .......... .......... .......... .......... .......... .......... ....G.. 107
U6_147_4 .......... .......... .......... .......... .......... .......... .......... .......... .......... .......... ....G.. 107
U6_155_4 .......... .......... .......... .......... .......... .......... .......... .......... .......... .......... ....G.. 107
U6_156_4 .......... .......... .......... .......... .......... .......... .......... .......... .......... .......... ....G.. 107
U6_158_4 .......... .......... .......... .......... .......... .......... .......... .......... .......... .......... ....G.. 107
U6_159_4 .......... .......... .......... .......... .......... .......... .......... .......... .......... .......... ....G.. 107
U6_160_4 .......... .......... .......... .......... .......... .......... .......... .......... .......... .......... ....G.. 107
U6_161_4 .......... .......... .......... .......... .......... .......... .......... .......... .......... .......... ....G.. 107
U6_163_4 .......... .......... .......... .......... .......... .......... .......... .......... .......... .......... ....G.. 107
U6_164_4 .......... .......... .......... .......... .......... .......... .......... .......... .......... .......... ....G.. 107
U6_173_4 .......... .......... .......... .......... .......... .......... .......... .......... .......... .......... ....G.. 107
U6_177_4 .......... .......... .......... .......... .......... .......... .......... .......... .......... .......... ....--- 104
U6_181_4 .......... .......... .......... .......... .......... .......... .......... .......... .......... .......... ....--- 104
U6_182_4 .......... .......... .......... .......... .......... .......... .......... .......... .......... .......... ....--- 104
U6_183_4 .......... .......... .......... .......... .......... .......... .......... .......... .......... .......... ....--- 104
U6_186_4 .......... .......... .......... .......... .......... .......... .......... .......... .......... .......... ....--- 104
U6_187_4 .......... .......... .......... .......... .......... .......... .......... .......... .......... .......... ....--- 104
U6_188_4 .......... ..-....... .......... .......... .......... .......... .......... .......... .......... .......... ....--- 103
U6_190_4 .......... .......... .......... .......... .......... .......... .......... .......... .......... .......... ....--- 104
U6_191_4 .......... ..-....... .......... .......... .......... .......... .......... .......... .......... .......... ....--- 103
U6_193_4 .......... .......... .......... .......... .......... .......... .......... .......... .......... .......... ....--- 104
U6_195_4 .......... .......... .......... .......... .......... .......... .......... .......... .......... .......... ....--- 104
U6_197_4 .......... .......... .......... .......... .......... .......... .......... .......... .......... .......... ....--- 104
U6_199_4 .......... .......... .......... .......... .......... .......... .......... .......... .......... .......... ....--- 104
U6_202_4 .......... .......... .......... .......... .......... .......... .......... .......... .......... .......... ....--- 104
U6_203_4 .......... .......... .......... .......... .......... .......... .......... .......... .......... .......... ....--- 104
U6_204_4 .......... .......... .......... .......... .......... .......... .......... .......... .......... .......... ....--- 104
U6_205_4 .......... .......... .......... .......... .......... .......... .......... .......... .......... .......... ....--- 104
U6_206_4 .......... .......... .......... .......... .......... .......... .......... .......... .......... .......... ....--- 104
U6_209_4 .......... .......... .......... .......... .......... .......... .......... .......... .......... .......... ....--- 104
U6_212_4 .......... .......... .......... .......... .......... .......... .......... .......... .......... .......... ....--- 104
U6_214_4 .......... .......... .......... .......... .......... .......... .......... .......... .......... .......... ....--- 104
U6_223_4 .......... .......... .......... .......... .......... .......... .......... .......... .......... .......... ....G.. 107
U6_224_4 .......... .......... .......... .......... .......... .......... .......... .......... .......... .......... ....G.. 107
U6_225_4 .......... .......... .......... .......... .......... .......... .......... .......... .......... .......... ....G.. 107
U6_226_4 .......... .......... .......... .......... .......... .......... .......... .......... .......... .......... ....G.. 107
U6_227_4 .......... .......... .......... .......... .......... .......... .......... .......... .......... .......... ....G.. 107
U6_229_4 .......... .......... .......... .......... .......... .......... .......... .......... .......... .......... ....G.. 107
U6_230_4 .......... .......... .......... .......... .......... .......... .......... .......... .......... .......... ....G.. 107
U6_231_4 .......... .......... .......... .......... .......... .......... .......... .......... .......... .......... ....G.. 107
U6_232_4 .......... .......... .......... .......... .......... .......... .......... .......... .......... .......... ....G.. 107
U6_233_4 .......... .......... .......... .......... .......... .......... .......... .......... .......... .......... ....G.. 107
U6_234_4 .......... .......... .......... .......... .......... .......... .......... .......... .......... .......... ....G.. 107
U6_235_4 .......... .......... .......... .......... .......... .......... .......... .......... .......... .......... ....G.. 107
U6_236_4 .......... .......... .......... .......... .......... .......... .......... .......... .......... .......... ....G.. 107
U6_237_4 .......... .......... .......... .......... .......... .......... .......... .......... .......... .......... ....G.. 107
U6_238_4 .......... .......... .......... .......... .......... .......... .......... .......... .......... .......... ....G.. 107
U6_240_4 .......... .......... .......... .......... .......... .......... .......... .......... .......... .......... ....G.. 107
U6_241_4 .......... .......... .......... .......... .......... .......... .......... .......... .......... .......... ....G.. 107
U6_242_4 .......... .......... .......... .......... .......... .......... .......... .......... .......... .......... ....G.. 107
U6_243_4 .......... .......... .......... .......... .......... .......... .......... .......... .......... .......... ....G.. 107
U6_244_4 .......... .......... .......... .......... .......... .......... .......... .......... .......... .......... ....G.. 107
U6_246_4 .......... .......... .......... .......... .......... .......... .......... .......... .......... .......... ....G.. 107
U6_247_4 .......... .......... .......... .......... .......... .......... .......... .......... .......... .......... ....G.. 107
U6_248_4 .......... .......... .......... .......... .......... .......... .......... .......... .......... .......... ....G.. 107
U6_250_4 .......... .......... .......... .......... .......... .......... .......... .......... .......... .......... ....G.. 107
U6_251_4 .......... .......... .......... .......... .......... .......... .......... .......... .......... .......... ....G.. 107
U6_253_4 .......... .......... .......... .......... .......... .......... .......... .......... .......... .......... ....G.. 107
U6_254_4 .......... .......... .......... .......... .......... .......... .......... .......... .......... .......... ....G.. 107
U6_262_4 .......... .......... .......... .......... .......... .......... .......... .......... .......... .......... ....G.. 107
U6_263_4 .......... .......... .......... .......... .......... .......... .......... .......... .......... .......... ....G.. 107
U6_266_4 .......... .......... .......... .......... .......... .......... .......... .......... .......... .......... ....G.. 107
U6_268_4 .......... .......... .......... .......... .......... .......... .......... .......... .......... .......... ....G.. 107
U6_277_4 .......... .......... .......... .......... .......... .......... .......... .......... .......... .......... ....G.. 107
U6_278_4 .......... .......... .......... .......... .......... .......... .......... .......... .......... .......... ....G.. 107
U6_284_4 .......... .......... .......... .......... .......... .......... .......... .......... .......... .......... ....G.. 107
U6_292_4 .......... .......... .......... .......... .......... .......... .......... .......... .......... .......... ....G.. 107
U6_295_4 .......... .......... .......... .......... .......... .......... .......... .......... .......... .......... ....G.. 107
U6_298_4 .......... .......... .......... .......... .......... .......... .......... .......... .......... .......... ....G.. 107
U6_299_4 .......... .......... .......... .......... .......... .......... .......... .......... .......... .......... ....G.. 107
U6_303_4 .......... .......... .......... .......... .......... .......... .......... .......... .......... .......... ....G.. 107
U6_307_4 .......... .......... .......... .......... .......... .......... .......... .......... .......... .......... ....G.. 107
U6_311_4 .......... .......... .......... .......... .......... .......... .......... .......... .......... .......... ....G.. 107
U6_317_4 .......... .......... .......... .......... .......... .......... .......... .......... .......... .......... ....--- 104
U6_322_4 .......... .......... .......... .......... .......... .......... .......... .......... .......... .......... ....--- 104
U6_334_4 .......... .......... .......... .......... .......... .......... .......... .......... .......... .......... ....--- 104
U6_336_4 .......... .......... .......... .......... .......... .......... .......... .......... .......... .......... ....--- 104
U6_337_4 .......... .......... .......... .......... .......... .......... .......... .......... .......... .......... ....--- 104
```

```
                                             20         40         60         80         100
                                              |          |          |          |          |
U6_338_4  .......... .......... .......... .......... .......... .......... .......... .......... .......... .......... ....---  104
U6_362_4  .......... .......... .......... .......... .......... .......... .......... .......... .......... .......... ....---  104
U6_364_4  .......... .......... .......... .......... .......... .......... .......... .......... .......... .......... ....---  104
U6_366_4  .......... .......... .......... .......... .......... .......... .......... .......... .......... .......... ....---  104
U6_375_4  .......... .......... .......... .......... .......... .......... .......... .......... .......... .......... ....G..  107
U6_378_4  .......... .......... .......... .......... .......... .......... .......... .......... .......... .......... ....G..  107
U6_382_4  .......... .......... .......... .......... .......... .......... .......... .......... .......... .......... ....G..  107
U6_384_4  .......... .......... .......... .......... .......... .......... .......... .......... .......... .......... ....G..  107
U6_385_4  .......... .......... .......... .......... .......... .......... .......... .......... .......... .......... ....G..  107
  U6_7_3  .......... .......... .......... .......... .......... .......... .......... .......... .......... .......... ....---  104
 U6_20_4  .......... .......... .......... .......... .......... .......... .......... .......... .......... .......... ....G..  107
 U6_22_4  .......... .......... .......... .......... .......... .......... .......... .......... .......... .......... ....G..  107
 U6_24_4  .......... .......... .......... .......... .......... .......... .......... .......... .......... .......... ....G..  107
 U6_27_4  .......... .......... .......... .......... .......... .......... .......... .......... .......... .......... ..-----  102
 U6_28_4  .......... .......... .......... .......... .......... .......... .......... .......... .......... .......... ....G..  107
 U6_29_4  .......... .......... .......... .......... .......... .......... .......... .......... .......... .......... ....G..  107
 U6_30_4  .......... .......... .......... .......... .......... .......... .......... .......... .......... .......... ....G..  107
U6_504_4  .......... .......... .......... .......... .......... .......... .......... .......... .......... .......... ....G..  107
U6_508_4  .......... .......... .......... .......... .......... .......... .......... .......... .......... .......... ....G..  107
U6_512_4  .......... .......... .......... .......... .......... .......... .......... .......... .......... .......... ....---  104
U6_513_4  .......... .......... .......... .......... .......... .......... .......... .......... .......... .......... ....---  104
U6_487_4  .......... .......... .......... .......... .......... .......... .......... .......... .......... .......... ....G..  107
U6_488_4  .......... .......... .......... .......... .......... .......... .......... .......... .......... .......... ....G..  107
U6_492_4  .......... .......... .......... .......... .......... .......... .......... .......... .......... .......... ....G..  107
U6_493_4  .......... .......... .......... .......... .......... .......... .......... .......... .......... .......... ...----  103
U6_494_4  .......... .......... .......... .......... .......... .......... .......... .......... .......... .......... ....G..  107
U6_497_4  .......... .......... .......... .......... .......... .......... .......... .......... .......... .......... ....G..  107
U6_500_4  .......... .......... .......... .......... .......... .......... .......... .......... .......... .......... ....G..  107
U6_501_4  .......... .......... .......... .......... .......... .......... .......... .......... .......... .......... ....G..  107
U6_446_4  .......... .......... .......... .......... .......... .......... .......... .......... .......... .......... ....G..  107
U6_449_4  .......... .......... .......... .......... .......... .......... .......... .......... .......... .......... ....---  104
U6_450_4  .......... .......... .......... .......... .......... .......... .......... .......... .......... .......... ....---  104
U6_453_4  .......... .......... .......... .......... .......... .......... .......... .......... .......... .......... ....---  104
U6_456_4  .......... .......... .......... .......... .......... .......... .......... .......... .......... .......... ....---  104
U6_457_4  .......... .......... .......... .......... .......... .......... .......... .......... .......... .......... ....---  104
U6_464_4  .......... .......... .......... .......... .......... .......... .......... .......... .......... .......... ....---  104
U6_465_4  .......... .......... .......... .......... .......... .......... .......... .......... .......... .......... ....---  104
U6_467_4  .......... .......... .......... .......... .......... .......... .......... .......... .......... .......... ....---  104
U6_472_4  .......... .......... .......... .......... .......... .......... .......... .......... .......... .......... ....---  104
U6_473_4  .......... .......... .......... .......... .......... .......... .......... .......... .......... .......... ....---  104
U6_474_4  .......... .......... .......... .......... .......... .......... .......... .......... .......... .......... ....---  104
U6_475_4  .......... .......... .......... .......... .......... .......... .......... .......... .......... .......... ....---  104
U6_480_4  .......... .......... .......... .......... .......... .......... .......... .......... .......... .......... ....---  104
U6_482_4  .......... .......... .......... .......... .......... .......... .......... .......... .......... .......... ....---  104
U6_486_4  .......... .......... .......... .......... .......... .......... .......... .......... .......... .......... ....G..  107
U6_386_4  .......... .......... .......... .......... .......... .......... .......... .......... .......... .......... ....G..  107
U6_389_4  .......... .......... .......... .......... .......... .......... .......... .......... .......... .......... ....G..  107
U6_391_4  .......... .......... .......... .......... .......... .......... .......... .......... .......... .......... ....G..  107
U6_392_4  .......... .......... .......... .......... .......... .......... .......... .......... .......... .......... ....G..  107
U6_393_4  .......... .......... .......... .......... .......... .......... .......... .......... .......... .......... ..-----  102
U6_395_4  .......... .......... .......... .......... .......... .......... .......... .......... .......... .......... ....G..  107
U6_396_4  .......... .......... .......... .......... .......... .......... .......... .......... .......... .......... ....G..  107
U6_397_4  .......... .......... .......... .......... .......... .......... .......... .......... .......... .......... ..-----  102
U6_398_4  .......... .......... .......... .......... .......... .......... .......... .......... .......... .......... ....G..  107
U6_399_4  .......... .......... .......... .......... .......... .......... .......... .......... .......... .......... ....G..  107
U6_400_4  .......... .......... .......... .......... .......... .......... .......... .......... .......... .......... ....G..  107
U6_401_4  .......... .......... .......... .......... .......... .......... .......... .......... .......... .......... ....G..  107
U6_403_4  .......... .......... .......... .......... .......... .......... .......... .......... .......... .......... ....G..  107
U6_404_4  .......... .......... .......... .......... .......... .......... .......... .......... .......... .......... ....G..  107
U6_405_4  .......... .......... .......... .......... .......... .......... .......... .......... .......... .......... ....G..  107
U6_408_4  .......... .......... .......... .......... .......... .......... .......... .......... .......... .......... ....G..  107
U6_409_4  .......... .......... .......... .......... .......... .......... .......... .......... .......... .......... ..-----  102
U6_411_4  .......... .......... .......... .......... .......... .......... .......... .......... .......... .......... ....G..  107
U6_412_4  .......... .......... .......... .......... .......... .......... .......... .......... .......... .......... ....G..  107
U6_415_4  .......... .......... .......... .......... .......... .......... .......... .......... .......... .......... ....G..  107
U6_417_4  .......... .......... .......... .......... .......... .......... .......... .......... .......... .......... ....G..  107
U6_418_4  .......... .......... .......... .......... .......... .......... .......... .......... .......... .......... ....G..  107
U6_419_4  .......... .......... .......... .......... .......... .......... .......... .......... .......... .......... ..-----  102
U6_422_4  .......... .......... .......... .......... .......... .......... .......... .......... .......... .......... ....G..  107
U6_428_4  .......... .......... .......... .......... .......... .......... .......... .......... .......... .......... ....G..  107
U6_431_4  .......... .......... .......... .......... .......... .......... .......... .......... .......... .......... ....G..  107
U6_432_4  .......... .......... .......... .......... .......... .......... .......... .......... .......... .......... ....G..  107
U6_437_4  .......... .......... .......... .......... .......... .......... .......... .......... .......... .......... ....G..  107
U6_440_4  .......... .......... .......... .......... .......... .......... .......... .......... .......... .......... ....G..  107
U6_441_4  .......... .......... .......... .......... .......... .......... .......... .......... .......... .......... ....G..  107
U6_443_4  .......... .......... .......... .......... .......... .......... .......... .......... .......... .......... ....G..  107
U6_444_4  .......... .......... .......... .......... .......... .......... .......... .......... .......... .......... ....G..  107
U6_270_4  .......... .......... .......... .......... ........A. .......... .......... .......... .......... .......... ....G..  107
U6_489_4  .......... .......... .......... .......... .......--- .......... .......... .......... .......... .......... ....G..  104
U6_279_4  .......... ..A....... .......... .......... .......... .......... .......... .......... .......... .......... ....G..  107
U6_316_4  .......... .......... .......... .......... .......... .......... .......G.. .......... .......... .......... ....---  104
U6_320_4  .......... .......... .......... .......... .......... .......... .......... .......... .......... .......A.. ....---  104
U6_452_4  .......... .......... .......... .......... .......... .......... .......... .......... .....T.... .......... ....---  104
 U6_77_4  .......... .......... .......... .......... .......... .......... .......... .......... .......... ........A. ....---  104
U6_357_4  .......... .......... .......... .......... .......... .......... .......... .......... .......... .........T ....---  104
U6_481_4  .......... .......... .......... .......... ...A...... .......... .......... .......... .......... .......... ....---  104
U6_354_4  .......... .......... .......... .......... .......... .......... .......... ....C..... .......... .......... ....---  104
U6_358_4  .......... .......... .......... .......... .......... .......... .......... ....C..... .......... .......... ....---  104
U6_367_4  .......... .......... .......... .......... .......... .......... .......... ....C..... .......... .......... ....---  104
U6_328_4  .......... .......... ....A..... .......... .......... .......... .......... .......... .......... .......... ....---  104
U6_361_4  .......... .......... ....T..... .......... .......... .......... .......... .......... .......... .......... ....---  104
```

```
                                  20          40          60          80         100
                                   |           |           |           |           |
U6_6_3    .......... .......... .......... .......... .......... .......... .......... .......... .........T .......... ....--- 104
U6_178_4  .......... .......... .......... .......... .......... .......... .......... .......... .........A .......... ....--- 104
U6_184_4  .......... .......... .......... .......... .......... .......... .......... .......... .........A .......... ....--- 104
U6_200_4  .......... .......... .......... .......... .......... .......... ......A... .......... .......... .......... ....--- 104
U6_256_4  .......... .......... .......... .......... .......... .......... ......A... .......... .......... .......... ....G.. 107
U6_454_4  .......... .......... .......... .......... .......... .......... ......A... .......... .......... .......... ....--- 104
U6_511_4  .......... .......... .......... .......... .......... .......... ......A... .......... .......... .......... ....--- 104
U6_92_4   .......... .......... .......... .......... .......... .......... .....A.... .......... .......... .......... ....--- 104
U6_319_4  .......... .......... .......... .......... .......... .......... .....A.... .......... .......... .......... ....--- 104
U6_324_4  .......... .......... .......... .......... .......... .......... .....A.... .......... .......... .......... ....--- 104
U6_333_4  .......... .......... .......... .......... .......... .......... .....A.... .......... .......... .......... ....--- 104
U6_339_4  .......... .......... .......... .......... .......... .......... .....A.... .......... .......... .......... ....--- 104
U6_439_4  .......... .......... .......... .......... .......... .......... .....A.... .......... .......... .......... ....G.. 107
U6_365_4  .......... .......... .......... .......... .......... .......... .....T.... .......... .......... .......... ....--- 104
U6_377_4  .......... .......... .......... .......... .......... .......... .....T.... .......... .......... .......... ....G.. 107
U6_435_4  .......... .......... .......... .......... .......... .......... .....T.... .......... .......... .......... ....G.. 107
U6_434_4  .......... .......... .......... ....T..... .......... .......... .......... .......... .......... .......... ....G.. 107
U6_414_4  .......... .......... .......... .......... .......... .......... .......... .....T.... .......... .......... ....G.. 107
U6_442_4  .......... .......... .......... .......... .......... .......... .......... .......... ..G....... .......... ....G.. 107
U6_490_4  .......... .......... .......... .......... .......... .......... .......... ......T... .......... .......... ....G.. 107
U6_21_4   .......... .......... .......... .......... .......... .......... .......... .......... .......... .....A.... ....G.. 107
U6_293_4  .......... .......... .......... .......A.. .......... .......... .......... .......... .......... .......... ....G.. 107
U6_496_4  .......... .......... .......... .......... .......... .......... .......... .A........ .......... .......... ....G.. 107
U6_296_4  .......... .......... .......... .......... .......... .......... ..T....... .......... .......... .......... ....G.. 107
U6_300_4  .......... .......... .......... .......... .......... .......... ..T....... .......... .......... .......... ....G.. 107
U6_312_4  .......... .......... .......... .......... .......... .......... ..T....... .......... .......... .......... ....G.. 107
U6_304_4  .......... .......... .......... .......... .......... .......... ..T....... .......... .......... .......... ....G.. 107
U6_308_4  .......... .......... .......... .......... .......... .......... ..T....... .......... .......... .......... ....G.. 107
U6_213_4  .......... .......... .......... .......... .......... .A........ .......... .......... .......... .......... ....--- 104
U6_485_4  .......... .......... .......... .......... .......... .A........ .......... .......... .......... .......... ....G.. 107
U6_31_4   .......... .......... .......... .......... .......... .......... .......... .......... .......A.. .......... ....G.. 107
U6_48_4   .......... .......... .......... .......... .......... .......... .......... .......... .......A.. .......... ....G.. 107
U6_447_4  .......... .......... .......... .......... .......... .......... .......... .......... .......A.. .......... ....--- 104
U6_171_4  .......... .......... .......... .......... .......... .......... .......... .......... .......A.. .......... ....G.. 107
U6_83_4   .......... .......... .......... .......... .......... .......... .......... .......... .......A.. .......... ....--- 104
U6_95_4   .......... .......... .......... .......... .......... .......... .......... .......... .......A.. .......... ....--- 104
U6_329_4  .......... .......... .......... .......... .......... .......... .......... .......... .......A.. .......... ....--- 104
U6_468_4  -......... .......... .......... .......... ......G... .......... .......... .......... .......... .......... ....--- 103
U6_15_4   .....T.... .......... .......... .......... .......... .......... .......... .......... .......... ....A..... ....G.. 107
U6_381_4  .....T.... .......... .......... .......... .......... .......... .......... .......... .......... ....A..... ....G.. 107
U6_314_4  .....T.... .......... .......... .......... .......... .......... .......... .......... ......A... .......... ....G.. 107
U6_148_4  .....T.... .......... .......... .......... .......... .......... .......... .......... .......... .......... ....G.. 107
U6_149_4  .....T.... .......... .......... .......... .......... .......... .......... .......... .......... .......... ....G.. 107
U6_150_4  .....T.... .......... .......... .......... .......... .......... .......... .......... .......... .......... ....G.. 107
U6_152_4  .....T.... .......... .......... .......... .......... .......... .......... .......... .......... .......... ....G.. 107
U6_153_4  .....T.... .......... .......... .......... .......... .......... .......... .......... .......... .......... ....G.. 107
U6_162_4  .....T.... .......... .......... .......... .......... .......... .......... .......... .......... .......... ....G.. 107
U6_239_4  .....T.... .......... .......... .......... .......... .......... .......... .......... .......... .......... ....G.. 107
U6_249_4  .....T.... .......... .......... .......... .......... .......... .......... .......... .......... .......... ....G.. 107
U6_280_4  .....T.... .......... .......... .......... .......... .......... .......... .......... .......... .......... ....G.. 107
U6_383_4  .....T.... .......... .......... .......... .......... .......... .......... .......... .......... .......... ....G.. 107
U6_388_4  .....T.... .......... .......... .......... .......... .......... .......... .......... .......... .......... ....G.. 107
U6_390_4  .....T.... .......... .......... .......... .......... .......... .......... .......... .......... .......... ....G.. 107
U6_394_4  .....T.... .......... .......... .......... .......... .......... .......... .......... .......... .......... ....G.. 107
U6_402_4  .....T.... .......... .......... .......... .......... .......... .......... .......... .......... .......... ....G.. 107
U6_407_4  .....T.... .......... .......... .......... .......... .......... .......... .......... .......... .......... ....G.. 107
U6_413_4  .....T.... .......... .......... .......... .......... .......... .......... .......... .......... .......... ....G.. 107
U6_420_4  .....T.... .......... .......... .......... .......... .......... .......... .......... .......... .......... ....G.. 107
U6_421_4  .....T.... .......... .......... .......... .......... .......... .......... .......... .......... .......... ....G.. 107
U6_423_4  .....T.... .......... .......... .......... .......... .......... .......... .......... .......... .......... ....G.. 107
U6_424_4  .....T.... .......... .......... .......... .......... .......... .......... .......... .......... .......... ....G.. 107
U6_429_4  .....T.... .......... .......... .......... .......... .......... .......... .......... .......... .......... ....G.. 107
U6_433_4  .....T.... .......... .......... .......... .......... .......... .......... .......... .......... .......... ....G.. 107
U6_502_4  .....T.... .......... .......... .......... .......... .......... .......... .......... .......... .......... ....G.. 107
U6_23_4   .....T.... .......... .......... .......... .......... .......... .......... .......... .......... .......... ..----- 102
U6_74_4   --...T.... .......... .......... .......... .......... .......... .......... .......... .......... .......... ....--- 102
U6_76_4   .....T.... .......... .......... .......... .......... .......... .......... .......... .......... .......... ....--- 104
U6_78_4   --...T.... .......... .......... .......... .......... .......... .......... .......... .......... .......... ....--- 102
U6_80_4   .....T.... .......... .......... .......... .......... .......... .......... .......... .......... .......... ....--- 104
U6_82_4   .....T.... .......... .......... .......... .......... .......... .......... .......... .......... .......... ....--- 104
U6_86_4   .....T.... .......... .......... .......... .......... .......... .......... .......... .......... .......... ....--- 104
U6_91_4   .....T.... .......... .......... .......... .......... .......... .......... .......... .......... .......... ....--- 104
U6_94_4   .....T.... .......... .......... .......... .......... .......... .......... .......... .......... .......... ....--- 104
U6_38_4   .....T.... .......... .......... .......... .......... .......... .......... .......... .......... .......... ....G.. 107
U6_44_4   .....T.... .......... .......... .......... .......... .......... .......... .......... .......... .......... ....G.. 107
U6_49_4   .....T.... .......... .......... .......... .......... .......... .......... .......... .......... .......... ....G.. 107
U6_53_4   .....T.... .......... .......... .......... .......... .......... .......... .......... .......... .......... ....G.. 107
U6_54_4   .....T.... .......... .......... .......... .......... .......... .......... .......... .......... .......... ....G.. 107
U6_57_4   .....T.... .......... .......... .......... .......... .......... .......... .......... .......... .......... ....G.. 107
U6_63_4   .....T.... .......... .......... .......... .......... .......... .......... .......... .......... .......... ....G.. 107
U6_66_4   .....T.... .......... .......... .......... .......... .......... .......... .......... .......... .......... ....G.. 107
U6_69_4   .....T.... .......... .......... .......... .......... .......... .......... .......... .......... .......... ....G.. 107
U6_132_4  .....T.... .......... .......... .......... .......... .......... .......... .......... .......... .......... ....G.. 107
U6_134_4  .....T.... .......... .......... .......... .......... .......... .......... .......... .......... .......... ....G.. 107
U6_138_4  .....T.... .......... .......... .......... .......... .......... .......... .......... .......... .......... ....G.. 107
U6_140_4  .....T.... .......... .......... .......... .......... .......... .......... .......... .......... .......... ....G.. 107
U6_143_4  .....T.... .......... .......... .......... .......... .......... .......... .......... .......... .......... ....G.. 107
U6_144_4  .....T.... .......... .......... .......... .......... .......... .......... .......... .......... .......... ....G.. 107
U6_145_4  .....T.... .......... .......... .......... .......... .......... .......... .......... .......... .......... ....G.. 107
U6_19_4   .....T.... .......... .......... .......... .......... .......... .......... .......... .......... .......... ....G.. 107
U6_32_4   .....T.... .......... .......... .......... .......... .......... .......... .......... .......... .......... ....G.. 107
```

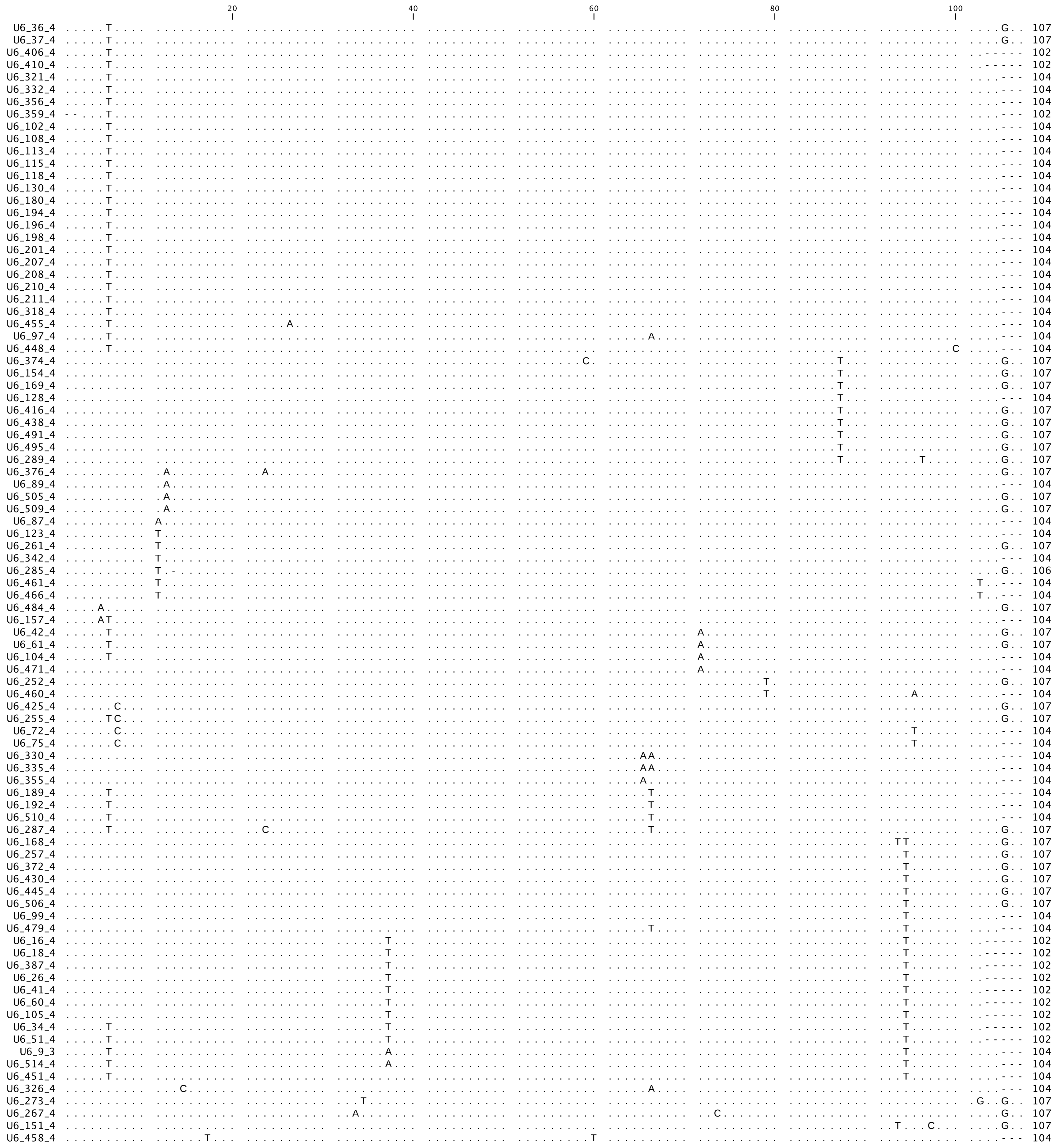
20
40
60
80
100
U6_36_4 107
U6_37_4 107
U6_406_4 102
U6_410_4 102
U6_321_4 104
U6_332_4 104
U6_356_4 104
U6_359_4 102
U6_102_4 104
U6_108_4 104
U6_113_4 104
U6_115_4 104
U6_118_4 104
U6_130_4 104
U6_180_4 104
U6_194_4 104
U6_196_4 104
U6_198_4 104
U6_201_4 104
U6_207_4 104
U6_208_4 104
U6_210_4 104
U6_211_4 104
U6_318_4 104
U6_455_4 104
U6_97_4 104
U6_448_4 104
U6_374_4 107
U6_154_4 107
U6_169_4 107
U6_128_4 104
U6_416_4 107
U6_438_4 107
U6_491_4 107
U6_495_4 107
U6_289_4 107
U6_376_4 107
U6_89_4 104
U6_505_4 107
U6_509_4 107
U6_87_4 104
U6_123_4 104
U6_261_4 107
U6_342_4 104
U6_285_4 106
U6_461_4 104
U6_466_4 104
U6_484_4 107
U6_157_4 104
U6_42_4 107
U6_61_4 107
U6_104_4 104
U6_471_4 104
U6_252_4 107
U6_460_4 104
U6_425_4 107
U6_255_4 107
U6_72_4 104
U6_75_4 104
U6_330_4 104
U6_335_4 104
U6_355_4 104
U6_189_4 104
U6_192_4 104
U6_510_4 104
U6_287_4 107
U6_168_4 107
U6_257_4 107
U6_372_4 107
U6_430_4 107
U6_445_4 107
U6_506_4 107
U6_99_4 104
U6_479_4 104
U6_16_4 102
U6_18_4 102
U6_387_4 102
U6_26_4 102
U6_41_4 102
U6_60_4 102
U6_105_4 102
U6_34_4 102
U6_51_4 102
U6_9_3 104
U6_514_4 104
U6_451_4 104
U6_326_4 104
U6_273_4 107
U6_267_4 107
U6_151_4 107
U6_458_4 104

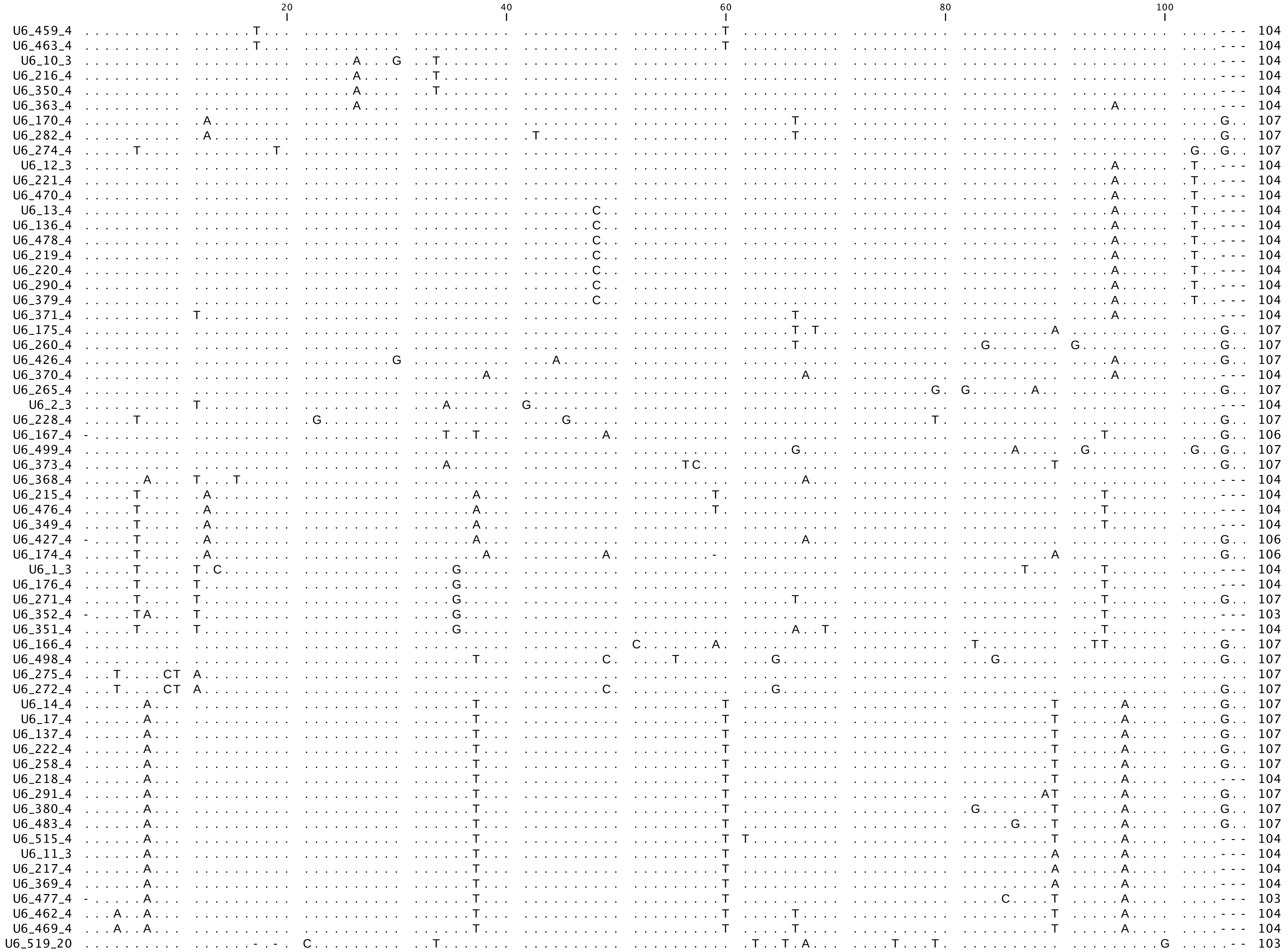
20
40
60
80
100
U6_459_4 104
U6_463_4 104
U6_10_3 104
U6_216_4 104
U6_350_4 104
U6_363_4 104
U6_170_4 107
U6_282_4 107
U6_274_4 107
U6_12_3 104
U6_221_4 104
U6_470_4 104
U6_13_4 104
U6_136_4 104
U6_478_4 104
U6_219_4 104
U6_220_4 104
U6_290_4 104
U6_379_4 104
U6_371_4 104
U6_175_4 107
U6_260_4 107
U6_426_4 107
U6_370_4 104
U6_265_4 107
U6_2_3 104
U6_228_4 107
U6_167_4 106
U6_499_4 107
U6_373_4 107
U6_368_4 104
U6_215_4 104
U6_476_4 104
U6_349_4 104
U6_427_4 106
U6_174_4 106
U6_1_3 104
U6_176_4 104
U6_271_4 107
U6_352_4 103
U6_351_4 104
U6_166_4 107
U6_498_4 107
U6_275_4 107
U6_272_4 107
U6_14_4 107
U6_17_4 107
U6_137_4 107
U6_222_4 107
U6_258_4 107
U6_218_4 104
U6_291_4 107
U6_380_4 107
U6_483_4 107
U6_515_4 104
U6_11_3 104
U6_217_4 104
U6_369_4 104
U6_477_4 103
U6_462_4 104
U6_469_4 104
U6_519_20 103

```
                                 20                   40                   60                   80                   100
                                  |                    |                    |                    |                    |
 U6Somatic GTGCTTGCTT CGGCAGCACA TATACTAAAA TTGGAACGAT ACAGAGAAGA TTAGCATGGC CCCTGCGCAA GGATGACACG CAAATTCGTG AAGCGTTCCA TCTTTTA 107
  U6_516_6 .......... .......... .......... .......... .......... .......... .......... .......... .......... .......... ....... 107
  U6_517_9 .......... .......... .......... .......... .......... .......... .......... .......... .......... .......... ....... 107
 U6_518_11 .......... .......... .......... .......... .......... .......... .......... .......... .......... .......... ....... 107
 U6_520_21 .......... .......... .......... .......... .......... .......... .......... .......... .......... .........T CA....- 106
```
